# Astrocyte regional specialization is shaped by postnatal development

**DOI:** 10.1101/2024.10.11.617802

**Authors:** Margaret E. Schroeder, Dana M. McCormack, Lukas R. Metzner, Jinyoung Kang, Katelyn X. Li, Eunah Yu, Lisa Melamed, Kirsten M. Levandowski, Heather Zaniewski, Qiangge Zhang, Edward S. Boyden, Fenna M. Krienen, Guoping Feng

## Abstract

Astrocytes are an abundant class of glial cells with critical roles in neural circuit assembly and function. Though many studies have uncovered significant molecular distinctions between astrocytes from different brain regions, how this regionalization unfolds over development is not fully understood. We used single-nucleus RNA sequencing to characterize the molecular diversity of brain cells across six developmental stages and four brain regions in the mouse and marmoset brain. Our analysis of over 170,000 single astrocyte nuclei revealed striking regional heterogeneity among astrocytes, particularly between telencephalic and diencephalic regions, at all developmental time points surveyed in both species. At the stages sampled, most of the region patterning was private to astrocytes and not shared with neurons or other glial types. Though astrocytes were already regionally patterned in late embryonic stages, this region-specific astrocyte gene expression signature changed dramatically over postnatal development, and its composition suggests that regional astrocytes further specialize postnatally to support their local neuronal circuits. Across mouse and marmoset, we found hundreds of species differentially expressed genes, as well as divergence in the expression of astrocytic region- and age-differentially expressed genes and the timing of astrocyte maturation relative to birth between the species. Finally, we used expansion microscopy to show that astrocyte morphology is also regionally specialized across cortex, striatum, and thalamus in the mouse.

## Introduction

The mammalian brain is composed of thousands of heterogeneous molecularly-defined cell types^1,2^. This heterogeneity is prominent between cells from different anatomical regions that arise from distinct developmental compartments. This regional specialization is critical for circuit formation and proper brain function. In recent years, this heterogeneity has been cataloged through large-scale single-cell and single-nucleus RNA sequencing (scRNAseq and snRNAseq, respectively), which enables molecular profiling in unprecedented detail and scale^3^. The past decade has seen the publication of multiple brain cell type transcriptomic atlases, including of the entire adult mouse brain^1,2^, the adult human brain^4^, the developing mouse^5^ and human^6^ brains, and the adult marmoset brain^7,8^. Most, but not all of these atlases have focused primarily on characterizing neurons, long considered the brain’s principal cell type. Indeed, several studies used cell sorting methods to enrich for neurons^9^.

Astrocytes, an abundant class of glia, play critical roles in neuronal circuit assembly and function in healthy and pathological states^10–14^. While their morphological heterogeneity has long been appreciated^15,16^, their molecular heterogeneity, particularly across brain regions, was only revealed more recently by microarray^17^ and bulk RNA sequencing studies^18–20^, and later by single-cell RNA sequencing studies^8,21–25^. Early lineage tracing studies in the mouse spinal cord and brain revealed that astrocyte precursors from different embryonic domains are molecularly distinct^26,27^. Adult mouse astrocytes maintain epigenetic marks from their region-restricted radial glia ancestors^28^, which may contribute to the significant heterogeneity of adult astrocyte populations. There is also abundant evidence supporting the role of extrinsic cues in astrocyte regionalization, including the formation of various cortical morphological subtypes from a shared astrocyte progenitor^29^, the up- or down-regulation of ion channels, transporters, receptors in response to neuronal inputs^30^, and the molecular and morphological adaptation of distinct developmentally-patterned septal astrocyte subtypes after cross-region heterotopic transplant^31^.

As has been found with neurons, it is likely that transcriptionally-defined astrocyte populations are developmentally influenced by their respective microenvironments and perform distinct functions. Yet, the developmental time course of astrocyte regional patterning, the composition of astrocyte subtypes over development, and the conservation of these features between rodents and primates remain unclear^32^. To address this knowledge gap, we applied snRNAseq to characterize astrocyte molecular diversity across six developmental stages and four brain regions in mouse and marmoset. To complement the transcriptomic studies, we characterized complex astrocyte morphology and protein localization at high resolution across brain regions using expansion microscopy.

We used single nucleus sequencing to generate a dataset of 1.4 million brain cell nuclei across multiple stages and brain regions in mouse and marmoset. A unified study, with data generated from a single lab using highly consistent methodology, has the advantage of reduced technical variation compared to datasets integrated across research groups, nuclei isolation protocols, and sequencing platforms, which is difficult to remove *in silico*^33,34^.

Our analysis shows that astrocytes are regionally patterned before birth and at all subsequent time points. Importantly, we found dramatic changes in the transcriptional signatures underlying astrocyte regional identity between birth and early adolescence in both species, highlighting the importance of postnatal regional cues in shaping astrocyte identity. We explored the functional implications of genes differentially expressed between astrocytes from different brain regions, and between astrocytes at different developmental time points. Furthermore, we identified both region-shared and region-divergent developmental transcriptional signatures in astrocytes.

Many of the region-, age-, and species-differentially expressed genes in astrocytes implicated morphogenesis pathways. Indeed, astrocyte morphology, which is highly ramified and complex, including sub-micron scale processes that contact synapses and blood vessels, is essential for their many functions^35,36^. Therefore, to assess whether astrocyte morphology is also regionally specialized, we used a new variant of expansion microscopy, ExR^37^ to characterize virally-labeled astrocyte morphology and nanoscale protein expression with enhanced resolution. We found that gray matter thalamic astrocytes in mice were significantly smaller and less complex than their striatal and cortical counterparts, alongside differences in protein expression.

## Results

### A multi-region transcriptomic atlas of the developing mouse and marmoset brain

To create the cross-region, cross-species, cross-development snRNAseq atlas, we dissected prefrontal cortex (PFC), motor cortex (MO), striatum, and thalamus from freshly harvested mouse and marmoset brains at late embryonic, neonatal, early adolescent, late adolescent, young adult, and aged timepoints and snap-froze the tissue (**Fig. 1A**). We collected tissues from 2 marmoset donors, one male and one female, at gestational day (GD)135, neonate, 7 months, 14 months, 30 months (4 donors, previously collected data in the lab) and 11+ years). For mouse, we collected tissue from 3 mouse biological replicates, at least one female, at E18.5, P4, P14, P32, P90, and 90 weeks (see **Table S1** for mapping of donors to biological replicates, see **Methods**). We generated single-nuclei suspensions from the snap-frozen tissue without enriching for any particular cell type, and generated single-nucleus transcriptomes using 10x Genomics Chromium v3.1 chemistry (see **Fig. S1** for sequencing coverage statistics). Though the adult (4 donors aged 29-32 months, together labeled 30 months) marmoset snRNAseq data was generated using a different nuclei isolation protocol and reference genome^7^, the data integrated very well across studies (**Fig. S2A**). The data were also well integrated across biological sex (**Fig. S2-3A**). Quantitative measures of integration quality^38,39^ suggest our integration is well-mixed across biological replicates while preserving true biological variability (donor mixing = 0.8981 for mouse and 0.9155 for marmoset; neighbor consistency= 0.6680 for mouse and 0.4694 for marmoset; and average silhouette width = 0.6437 for mouse and 0.6634 for marmoset; see **Methods**).

**Figure 1.**
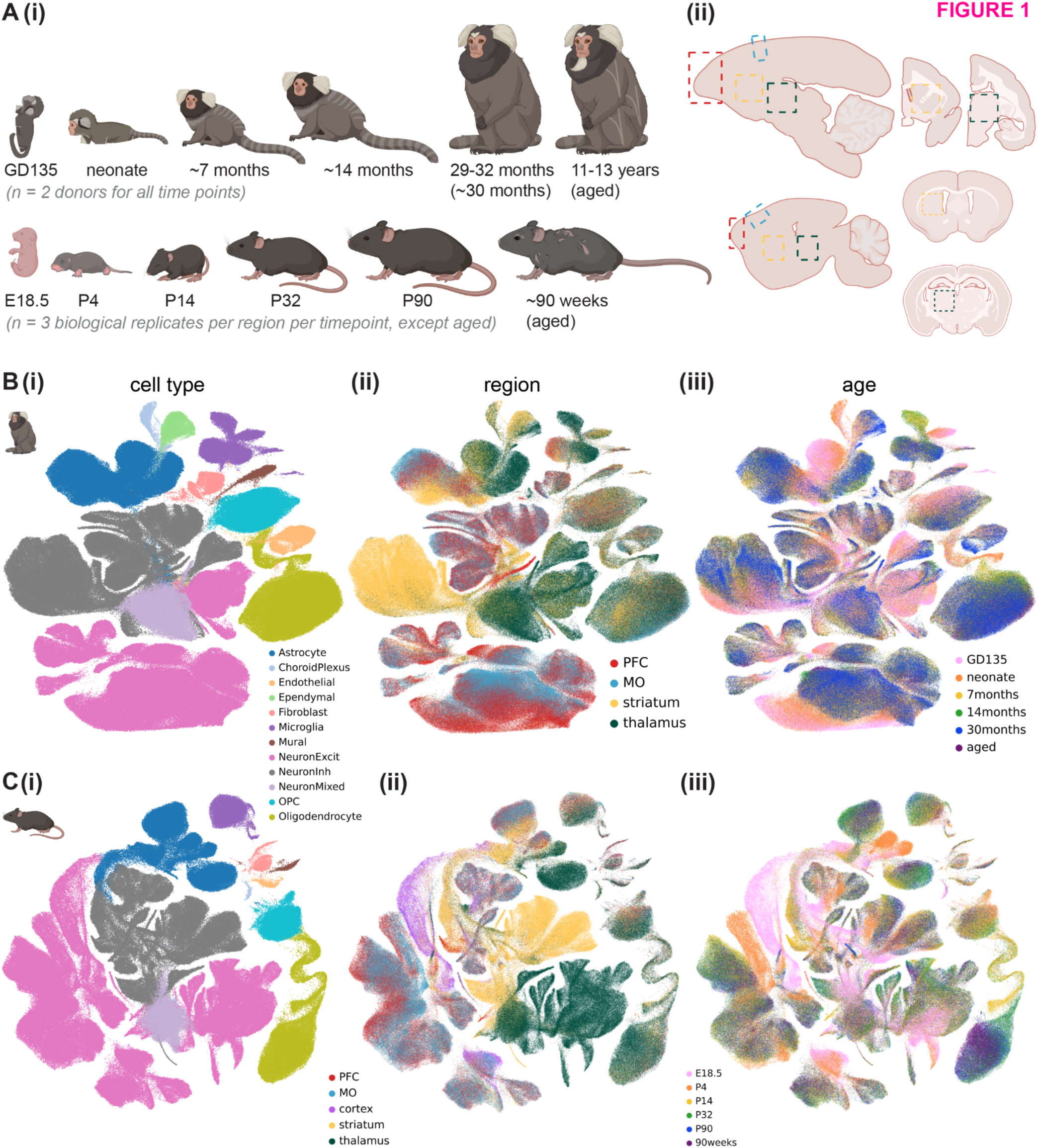
A multi-region transcriptomic atlas of brain cell diversity across postnatal development in marmoset and mouse. **A**, Cross-development, cross-region sampling strategy in marmoset (top row) and mouse (bottom row). **(i)** Developmental time points profiled (some approximate, see **Methods** and **Table S1**) GD, gestational day; E, embryonic day; P, postnatal day. **(ii)** Brain regions profiled, including prefrontal cortex (PFC, red dashed boxes), motor cortex (MO, blue dashed boxes), striatum (yellow dashed boxes), and thalamus (green dashed boxes), shown in either sagittal (left) or coronal slices (right, for subcortical regions only). Schematics generated using BioRender.com. **B-C**, Integrated UMAP embedding of marmoset (**B**, 881,832 nuclei) or mouse (**C**, 597,668 nuclei) nuclei from PFC, MO, striatum, and thalamus across all developmental time points assayed and a randomly downsampled portion of adult nuclei from our previous study^7^ colored by **(i)** assigned cell type **(ii)** dissected brain region, or **(iii)** developmental time point. Legend for **B-C(i)** is shared.

After rigorous quality control, including removal of ambient RNA, low-quality nuclei, and doublets (see **Methods**), we obtained 597,668 mouse nuclei and 881,832 marmoset nuclei, which were composed of 12 broad cell classes (**Fig. 1B(i), Fig. 1C(i); Fig. S2-3**). We annotated more granular cell type (Leiden^40^-determined) clusters within each cell class (**Fig. S4**, **Tables S2-4**). We employed the Allen Brain Cell Atlas’s (ABCA) MapMyCells^41^ portal^41^ to refine our annotations of neuronal subtypes and to help correct for modest cross-region contamination resulting from dissection error (**Fig. S5-6**, see **Methods**). Throughout the paper, “dissected” brain region refers to the original region label for the sample in which the nucleus was processed, while “region” or “assigned region” refers to the brain region assigned post-hoc in the case of cells informatically predicted to arise from neighboring structures.

The total neuron-to-astrocyte ratio (across regions and developmental time points) was modestly higher in mouse (6.28) than marmoset (5.24). To more quantitatively assess cell type composition differences across dissected region (cortex, striatum, or thalamus), age, and sex, we used single-cell compositional data analysis (scCODA)^42^, which implements a Bayesian model of cell type counts to address the issue of low sample sizes in snRNAseq data. scCODA confirmed many significant differences in cell type proportion between regions and ages for each species, including the expected low numbers of excitatory neurons in striatum, increasing oligodendrocyte abundance with age in both species, and minimal sex differences in cell type composition (**Fig. S2-3C**, **Tables S5-8**).

We found several cell type clusters that were enriched or depleted in developing (late embryonic or neonate) brains (**Fig. S4C,F**). For example, in both species, there were immature cortical excitatory neuron, microglia, and astrocyte clusters composed mostly of nuclei from late embryonic and fetal donors. The committed oligodendrocyte precursor (COP) and newly formed oligodendrocyte (NFOL) cluster was primarily composed of nuclei from the neonate time point in marmoset, with some nuclei even coming from late embryonic donors, but was primarily composed of nuclei from early adolescent (P14) donors in mouse, with no COP/NFOLs coming from E18.5 mouse, indicating earlier oligodendrocyte maturation in marmoset.

### Astrocyte regional heterogeneity is embryonically patterned and unfolds over postnatal development

We observed striking regional heterogeneity among astrocytes at all developmental time points sampled in both species, particularly between astrocytes of diencephalic (thalamus) and telencephalic (cortex and striatum) origin (**Fig. 2A**, **Fig. 3A**). This is in line with multiple studies demonstrating embryonic regional patterning of astrocytes^6,8,27^. These regional populations further divided into an immature population, primarily composed of nuclei from late embryonic and neonatal time points, and a mature population, composed of nuclei from late adolescent timepoints onward. These separate populations suggest embryonically-patterned regional astrocyte populations undergo significant changes from the time of birth (neonate or P4) to early adolescence (7 months or P14). Notably, abundant populations of immature astrocytes remained present in the mouse striatum through adulthood (P90).

**Figure 2.**
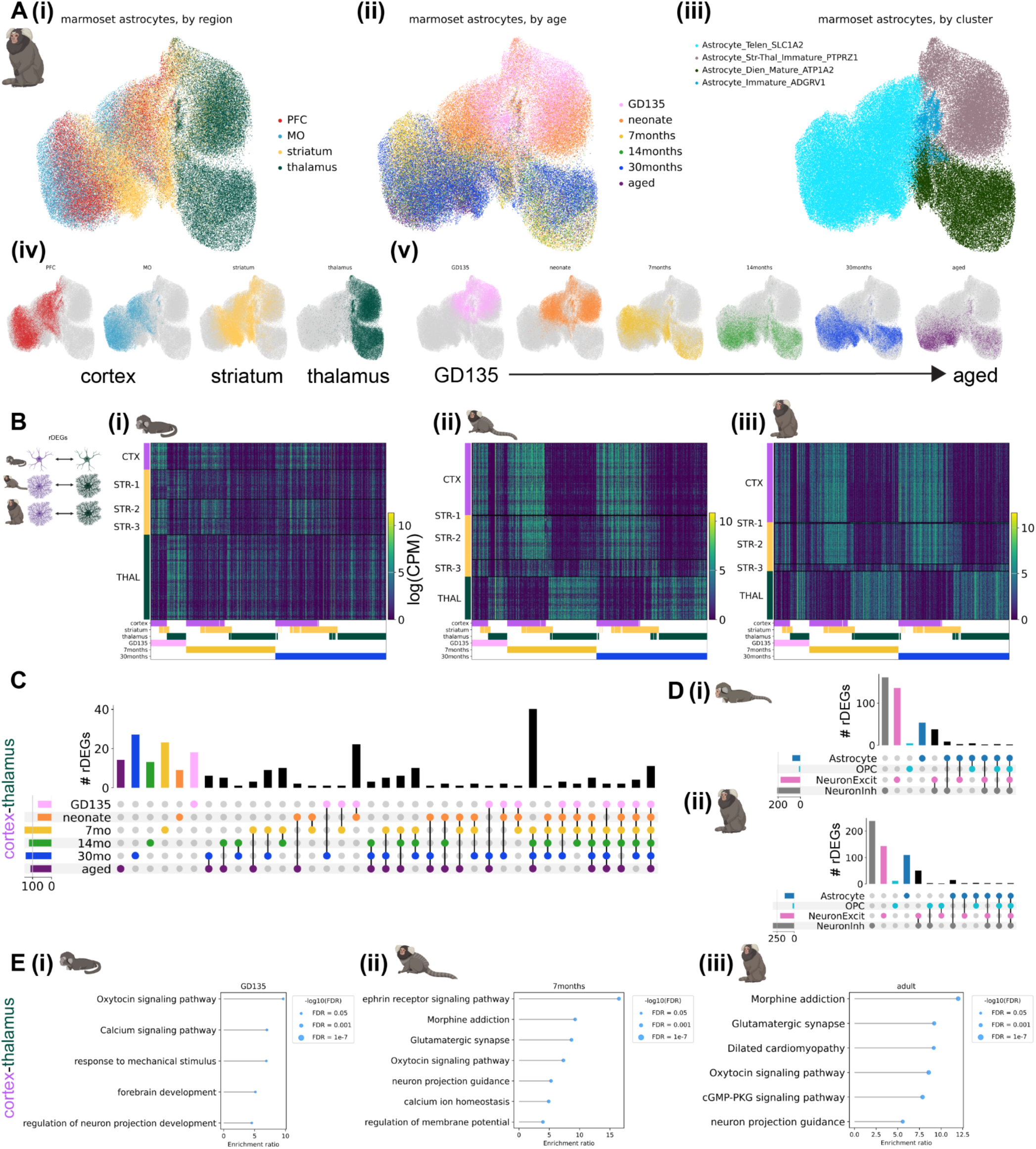
Developmental changes and cell-type specificity of astrocyte regional heterogeneity over postnatal development in the marmoset. **A,** Integrated UMAP embeddings of 103,009 marmoset astrocytes colored by **(i)** assigned brain region (one-hot color encoded in **(iv)**), **(ii)** developmental time point (one-hot color encoded in **(v)**, and **(iii)** Leiden cluster assignment. **B,** Expression heatmaps (rows are cells, columns are genes) of regional differentially expressed genes (rDEGs) between astrocytes from cortex, striatum, and thalamus at **(i)** GD135, **(ii)** 7 month, and **(iii)** 30 month-old marmosets in log counts per million (logCPM). The raster plots beneath each heatmap indicate the time point (GD135, 7 months, or 30 months) and region(s) of upregulation (cortex, striatum, and/or thalamus) for each rDEG. Genes are ordered first by the time point at which they are an rDEG, then by the regions in which they are most highly expressed, and are plotted more than once if they are present at more than one time point. The same set of genes are plotted in the same order in **(i-iii)**. Striatal astrocytes are ordered by subtype identity (see **Methods**). **C,** UpSet plot showing the number of unique and overlapping cortex-thalamus rDEGs between developmental time points. The colored dots below each vertical bar indicate which age(s) share that set of rDEGs, while the colored horizontal bars indicate the total number of cortex-thalamus rDEGs for each age. Overlap categories with 0 rDEGs are not shown. **D,** UpSet plot (as in **(C)**) showing the number of overlapping cortex-thalamus rDEGs between OPCs (light blue), astrocytes (dark blue), excitatory neurons (pink), and inhibitory neurons (gray) for **(i)** neonate and **(ii)** adult marmoset. **E,** Gene ontology (GO) and pathway analysis on cortex-thalamus (enriched in either region) astrocyte rDEGs via WebGestalt 2024 in **(i)** GD135, **(ii)** 7 month, and **(iii)** 30 month marmoset astrocytes. Lollipop plots show the enrichment ratio of GO Biological Process and KEGG pathways from an over-representation analysis with weighted set cover redundancy reduction, with tip size inversely proportional to the false discovery rate (FDR).

**Figure 3.**
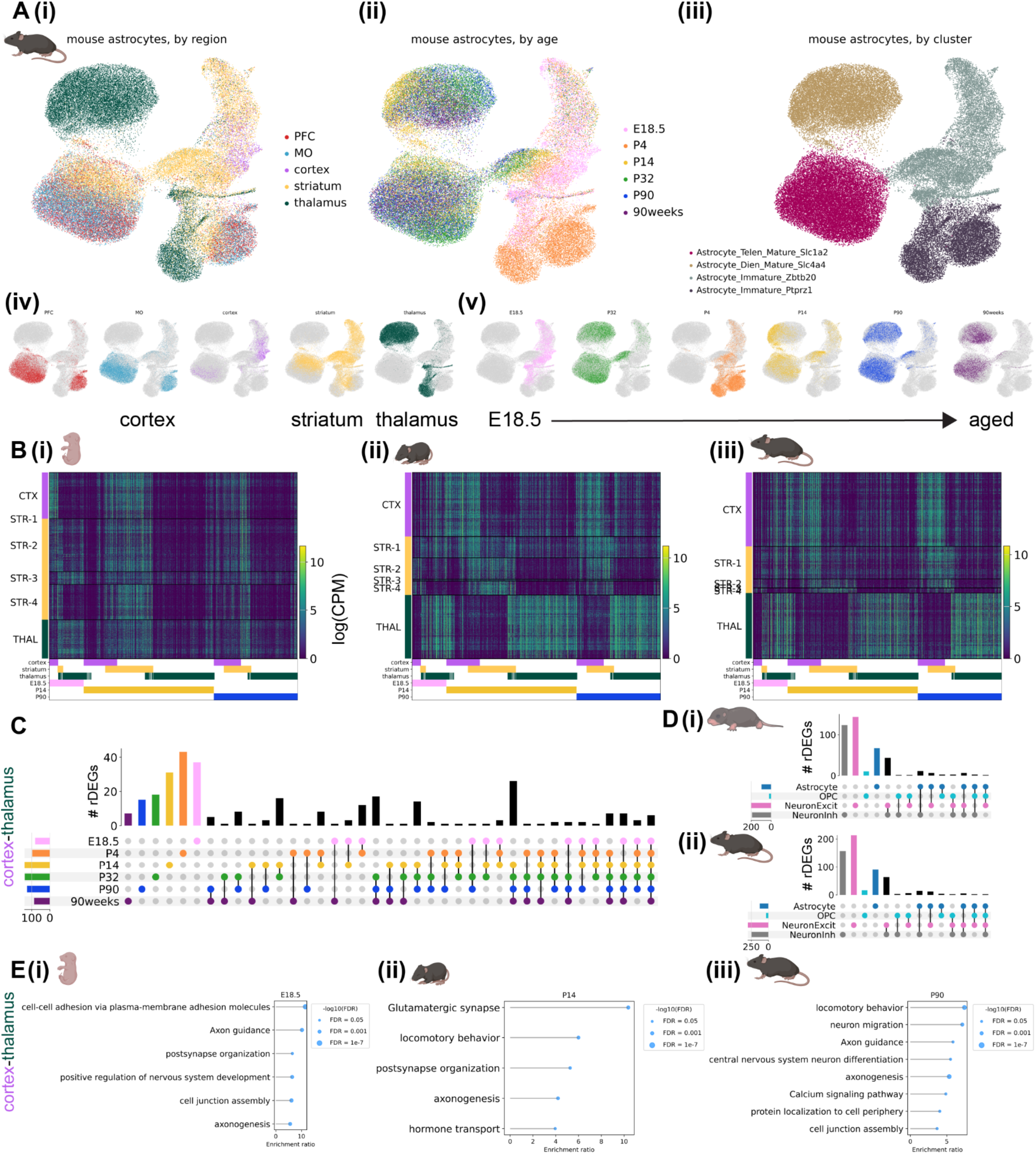
Developmental changes and cell-type specificity of astrocyte regional heterogeneity over postnatal development in the mouse. **A,** Integrated UMAP embeddings of 68,485 mouse astrocytes colored by **(i)** brain region (one-hot color encoded in **(iv)**), **(ii)** developmental time point (one-hot color encoded in **(v)**), and **(iii)** Leiden cluster assignment. **B,** Expression heatmap (rows are cells, columns are genes) of regional differentially expressed genes (rDEGs) between astrocytes from cortex, striatum, and thalamus at **(i)** E18.5, **(ii)** P14, and **(iii)** P90 mice in log counts per million (logCPM). The raster plots beneath each heatmap indicate the time point (GD135, 7 months, or 30 months) and region(s) of upregulation (cortex, striatum, and/or thalamus) for each rDEG. Genes are ordered first by the time point at which they are an rDEG, then by the regions in which they are most highly expressed, and are plotted more than once if they are present at more than one time point. The same set of genes are plotted in the same order in **(i-iii)**. Striatal astrocytes are ordered by subtype identity (see **Methods**). **C,** UpSet plot showing the number of overlapping cortex-thalamus rDEGs between developmental time points. The colored dots below each vertical bar indicate which age(s) share that set of rDEGs, while the colored horizontal bars indicate the total number of cortex-thalamus rDEGs for each age. Overlap categories with 0 rDEGs are not shown. **D,** UpSet plot (as in **(C)**) showing the number of overlapping cortex-thalamus rDEGs between OPCs (light blue), astrocytes (dark blue), inhibitory neurons (gray), and excitatory neurons (pink) for **(i)** P4 and **(ii)** P90 mouse. Overlap categories with 0 rDEGs are not shown. **E,** Gene ontology (GO) and pathway analysis on cortex-thalamus (enriched in either region) astrocyte rDEGs via WebGestalt 2024 in **(i)** E18.5, **(ii)** P32, and **(iii)** P90 month mouse astrocytes. Lollipop plots show the enrichment ratio of GO Biological Process and KEGG pathways from an over-representation analysis with weighted set cover redundancy reduction, with tip size inversely proportional to the false discovery rate (FDR).

#### Marmoset

Transcription factors and morphogen gradients set up initial boundaries between developmental compartments such as the telencephalon and diencephalon^43,44^. It could be that such early influences are present only transiently at initial astrocyte specification, or that later stages retain initial molecular distinctions and accumulate others over development. A combination of the two is also possible, where some genes follow one pattern (developmentally transient expression) or the other (sustained expression throughout the lifespan). We calculated **r**egional **d**ifferentially **e**xpressed **g**enes (rDEGs) at each developmental time point from metacells, 1-dimensional vectors of averaged normalized expression across all cells in a given grouping (see **Methods**), of each region. For marmoset, where each donor (biological replicate) was represented in each brain region, we calculated rDEGs separately for each donor and required that rDEGs be above threshold (minimum expression and log fold-change) requirements in both donors. We found 70 rDEGs whose expression differed between fetal cortical and thalamic astrocytes, 142 of such rDEGs by early adolescence (7 months), and 134 in adulthood (30 months, see **Fig. 2B(i-iii)** for the expression pattern of the union of these rDEGs at each timepoint). Focusing on cortex-thalamus rDEGs (which were most numerous, **Table S9**), we found that relatively few persisted across all developmental timepoints (**Fig. 2C**). 50% were shared between fetal and neonate, and 51% between late adolescent and aged, but only 4% continued to act as regional patterning signatures throughout the lifespan. We found that many more rDEGs were shared between late adolescent, young adult, and aged time points (61) than between fetal, neonate, and 7-month time points (23) and neonate, 7-month, and 14-month time points (25, **Fig. 2C**). These data suggest that regional astrocyte gene expression signatures emerge in the embryonic brain, change drastically over the course of early postnatal development and stabilize during adolescence into adulthood.

If early telencephalic and diencephalic patterning persists in astrocytes, cortex and striatum should retain common rDEGs compared to thalamus. To assess the degree of pairwise astrocyte rDEGs sharing across the 3 brain structures, we correlated the log fold-change difference in astrocyte regional gene expression between different region pairs (e.g., cortex-striatum vs. cortex-thalamus) for both rDEGs (log fold-change > 0.5) and non-rDEGs at fetal, early adolescent, and adult timepoints. We found that cortex-striatum vs. cortex-thalamus fold-changes exhibited a high degree of correlation in fetal marmoset (Pearson’s r = 0.78), which decreased over developmental time (Pearson’s r = 0.47 in adult marmoset, **Fig. S7A-C(i)**). At all 3 time points, striatum-thalamus vs. striatum-cortex fold-changes were negatively or uncorrelated (r = -0.43, -0.26, and 0.04 at GD135, 7 months, and 30 months respectively, **Fig. S7A-C(ii)**). Finally, thalamus-striatum vs. thalamus-cortex fold-changes were highly correlated (r > 0.80) at all 3 timepoints (**Fig. S7A-C(iii)**). Together, these results suggest that cortical and striatal astrocytes share transcriptional divergence from thalamic astrocytes at all ages, but become more transcriptionally similar later in development. At the same time, cortical and thalamic astrocytes both diverge from striatal astrocytes, but in distinct ways, as indicated by the negative correlation between striatum-thalamus and striatum-cortex fold-changes at GD135 and 7 months, and the lack of correlation at 30 months.

Dorsal radial glia populate the neocortex in a stereotyped progression, giving rise first to glutamatergic neurons, then to astrocytes, and finally to oligodendrocytes^45–47^. In the developing thalamus, radial glial progenitors likely follow the same cell type sequence^46,48^. As we showed previously for variable genes across the adult neocortex^7^, far more adult cortex-thalamus rDEGs are private to astrocytes than are shared with neurons or OPCs, despite their shared lineage^47^ (**Fig. 2D**). Surprisingly, this remained true even at the earliest stages we sampled (GD135 and neonate). Together these observations suggest that astrocytes gain regional identity early in their maturation, but that their continued regional identity is facilitated by distinct genes across their lifespan (**Fig. 2C**).

Many rDEGs nominate core cellular functions that may be further regionally specialized in astrocytes. For example, ephrins such as *EFNB2* (identified as rDEG in 30 month marmoset astrocytes) and *EFNA5* (7, 14, and 30 months and aged) are up-regulated in cortical astrocytes and ephrin receptor *EPHB1* (7, 14, and 30 months and aged) is upregulated in cortical and striatal astrocytes. Neuron-astrocyte signaling via ephrin ligands and receptors regulates axon guidance and synaptogenesis^49^. Thus, neuron-astrocyte ephrin signaling may be specialized in the telencephalon. Cyclic-AMP-related signaling molecules *ADCY1* (30 months and aged) and *ADCY8* (identified at all ages) are upregulated in thalamic astrocytes. As in neurons, astrocytic cAMP is an important second messenger following GPCR activation^50^, and modulates synaptic plasticity^51^. *ITPR1* (7, 14, and 30 months and aged), a calcium channel that controls calcium release from the endoplasmic reticulum, an important source of intracellular calcium during astrocyte signaling^52^, is also upregulated in thalamic astrocytes. These rDEGs suggest that thalamic astrocytes may have developed specialized pathways for calcium and cAMP signaling, potentially in response to the release of upstream GPCR ligands by thalamus-projecting and thalamic neurons. Additionally, astrocyte rDEGs included ion channels (e.g. *TRPM3* (GD135, 7, 14, and 30 months, and aged), a non-selective Ca^2+^ permeable ion channel and thalamic rDEG); synapse-related proteins (e.g., *SPARC, a thalamic rDEG at* 7, 14, and 30 months and aged, which regulates synaptogenesis^53^); neurotransmitter transporters and receptors (e.g., *SLC6A11*/GAT3 (7, 14, and 30 months and aged), a thalamic rDEG and GABA transporter, *SLC1A3*/GLAST (at GD135 and neonate), a glutamate transporter higher in the cortex and commonly used astrocyte marker gene, and *GRM3*/mGluR3 (all ages except GD135), a telencephalic rDEG and metabotropic glutamate receptor); and a thyroid hormone receptor (*SLCO1C1* (GD135, neonate, and 30 months), a cortical and later striatal rDEG). These rDEGS point more directly to astrocyte adaptation to the local synaptic and neuronal niche.

To characterize astrocyte rDEG pathways in a more unbiased manner, we used WebGestalt 2024^54,55^ over-representation analysis to test for enrichment of cortex-thalamus rDEGs (bidirectionally, i.e. upregulated either in cortex or thalamus) in GO Biological Process and KEGG pathways. Enriched pathways implicated oxytocin and calcium signaling and neuronal projection development for GD135 astrocyte rDEGs; ephrin signaling, synaptic transmission, and calcium ion homeostasis for 7-month astrocyte rDEGs; and glutamatergic synaptic transmission, oxytocin signaling, and cGMP-PKG signaling for adult marmoset astrocyte rDEGs (**Fig. 2E**). A summary of WebGestalt results for cortex-thalamus astrocyte rDEGs each age is provided in **Table S10**. To complement this pathway analysis and facilitate exploration of rDEG functions, we queried UniProt^56^ for each rDEG (see **Methods**) to return its full protein name, GO Cellular Compartment, GO Molecular Function, and GO Biological Process annotation(s). These annotations are included in **Table S9** for marmoset and **Table S10** for mouse. Together, these results suggest that astrocytes are regionally specialized with varied physiological adaptations necessary to support neuronal transmission and activity in their local environment.

Compared to cortex-thalamus expression differences, there were many fewer cortex-striatum rDEGs (0 in fetal marmoset, 12 in neonate, 8 in 7-month, 9 in 14-month, 25 in 30-month, and 12 in aged). At neonate, 7- and 14-month time points, at least half of these cortex-striatum rDEGs overlapped with cortex-thalamus rDEGs. At 14 months, these overlapped genes included *MYO16*, an unconventional myosin protein implicated in neurodevelopment^57^ (higher in striatum and thalamus); *UNC5C*, a netrin receptor family member involved in axon guidance^58,59^ (higher in cortex); *MAPK10*, a mitogen-activated protein kinase (higher in striatum and thalamus); *STXBP6*, a syntaxin binding protein which is part of the SNARE complex in neurons (higher in striatum and thalamus); *GRIK2*, a kainate-type ionotropic glutamate receptor subunit (higher in striatum and thalamus); *EYA2*, a transcriptional coactivator and phosphatase (higher in striatum and thalamus); *DYNC1I1*, a member of the cytoplasmic dynein 1 complex involved in intracellular transport (higher in striatum and thalamus); and *PTPRE*, a protein tyrosine phosphatase family member involved in cell signaling with various downstream consequences (higher in cortex). Each of these genes points to a biological process, such as glutamate sensing, phosphorylation, and exocytosis, for which striatal and thalamic astrocytes may be differentially invested.

#### Mouse

We calculated mouse astrocyte rDEGs using the previously described metacell method on expression data pooled across all biological replicates (see **Methods**). As with marmoset, mouse astrocyte gene expression varied across developmental timepoints, and most astrocyte rDEGs were not shared with other cell types (**Fig. 3A-D, Table S11**). E18.5 astrocyte cortex-thalamus rDEGs (79 total) included *Cacna2d1*, *Cntn5*, *Nrxn1*, *Creb5*, *Slco1c1*, and *Slc6a11*, and together were enriched for biological processes including cell-cell adhesion, axon guidance, and postsynaptic organization (**Fig. 3E(i), Table S12**). P14 astrocyte cortex-thalamus rDEGs included 26 of the rDEGs present at E18.5 (19% of total P14 rDEGs), in addition to rDEGs that only emerged at P14. These included voltage-gated calcium channel subunit *Cacna1a*, the glutamate-gated kainate receptor *Grik4*, the N-glycoprotein *Thsd7a*, the cholesterol transporter *Gramd1b*, and the inward-rectifying potassium channel *Kcnj6*. Together, P14 cortex-thalamus astrocyte rDEGs were enriched in glutamatergic synapse, hormone transport, and postsynaptic organization pathways (**Fig. 3E(ii), Table S12**). There were 124 cortex-thalamus astrocyte rDEGs at P90, which included many of the rDEGs present at earlier time points (17% of P90 rDEGs were present at E18.5 and 52% were present at P14), and were enriched in neuron migration, axon guidance, calcium signaling, and cell junction pathways (**Fig. 3E(iii), Table S12**). Compared to marmoset, mouse astrocytes had more cortex-striatum astrocyte rDEGs throughout development, especially at P14 (13 at E18.5, 24 at P4, 63 at P14, 15 at P90, and 33 at 90 weeks). However, at most 15 of these (at P14) overlapped with cortex-thalamus rDEGs, suggesting a more distinct transcriptional niche for mouse striatal astrocytes compared to cortex, as explored below.

As we did with marmoset, we assessed in mouse the degree to which regional imprinting of astrocyte gene expression persists across development by correlating log-fold change gene expression differences between region pairs at E18.5, P14, and P90. We found that cortex-striatum vs. cortex-thalamus fold-changes exhibited a high degree of correlation at E18.5 (Pearson’s r = 0.85), which decreased dramatically at P14 (r = 0.26) and increased again at P90 (r = 0.40, **Fig. S7D-F(i)**). Compared to marmoset, striatum-cortex vs. striatum-thalamus rDEGs were more positively correlated at juvenile and adult stages (P14 (r = 0.48) and P90 (r = 0.19, **Fig. S7D-F(ii)**)). Consistent with marmoset, thalamus-striatum vs. thalamus-cortex fold-changes were highly correlated (r > 0.70) at all 3 timepoints (**Fig. S7D-F(iii)**). Together, these results suggest that while cortical and striatal astrocytes are similarly divergent from thalamic astrocytes (likely reflecting the their distinct telencephalic-diencephalic origins), mouse striatal astrocytes develop and maintain a unique transcriptional signature distinct from cortex and thalamus.

To validate the existence of these regional astrocyte populations and the differential expression of selected rDEGs *in situ*, we conducted multiplexed RNA fluorescence *in situ* hybridization (FISH) using the RNAscope HiPlex or RNAscope Multiplex Fluorescent v2 kit (Advanced Cell Diagnostics) in neonate and adult animals of both species. We used CellProfiler 4.2.5^60,61^ to quantify the fraction of astrocytes positive for each target gene in each region and the fraction of each astrocyte nuclei covered by the probe for each target gene in each region (referred to as mean intensity, see **Methods**). Most rDEGs followed the expected regional and developmental expression pattern in marmoset astrocytes, including *SPARC*, which was enriched in diencephalic astrocytes (more so in adulthood), *FOXG1*, which marked telencephalic astrocytes, *GFAP,* which was elevated in thalamus in adult but not neonate, and *KCNH7*, which was a telencephalic rDEG in neonate but not adult (**Fig. S8-9**, see **Methods** and **Supplementary Note 1** for a discussion of the few rDEGs whose in situ expression differed from snRNAseq predictions).

Similarly, we found that most mouse astrocyte rDEGs followed the expected regional and developmental expression pattern in P4 and P90 mouse astrocytes, including *Clmn, Slco1c1, Csmd1*, *Sparc* (in adult mouse), and *Kcnd2* (**Figs. S10-11,** see **Table S13** for source data and statistics). For additional validation, we analyzed the differential expression of our selected mouse rDEGs in the Allen Mouse Brain Cell Atlas whole-brain MERSCOPE v1 dataset^1^, and found it to be largely consistent with our snRNAseq data (**Fig. S11B**, see **Methods**). This was also true for the whole list P90 mouse rDEGs, which also showed differential expression across a wider selection of brain regions (**Fig. S11C**).

To assess the extent of astrocyte intra-regional heterogeneity, we performed subclustering on cortical, striatal, and thalamic astrocytes from all developmental time points separately for each species (**Fig. S12-13**, see **Methods**). We found at least 4 astrocyte subclusters within each region, which primarily distinguished protoplasmic and fibrous/interlaminar subtypes (the latter being identified by *GFAP*, *AQP4*, and/or *ID3* expression^8,23,62,63^) and immature and mature astrocytes. In both species, the majority of astrocytes in the cortex and striatum were protoplasmic. In the marmoset thalamus, a larger proportion of astrocytes were *GFAP+*, *AQP4+*, or *ID3+* (**Fig. S12C, Table S14**), suggesting a higher proportion of fibrous astrocytes, consistent with the greater abundance of white matter in this region (**Fig. S2B-C**). Nevertheless, it is unclear the extent to which the definitions of protoplasmic, fibrous, and intralaminar apply outside of the cortex. The mouse striatum had the most intra-regional heterogeneity, with 12 subclusters (**Fig. S13B**), in large part due to immature populations including *Top2a+* rostral migratory stream progenitors. As in previous studies, we found that *CRYM/Crym* marks a subset of striatal astrocytes^19^ and *SPARC/Sparc* marks thalamic astrocytes in both species^18^. Several of our subclusters mapped specifically (>70% of cells in the subcluster) to a single adult mouse Allen Brain Cell Atlas (ABCA) cluster, though many mapped to several ABCA clusters, especially immature and mixed fibrous/protoplasmic subclusters (**Table S15**). Taken together with our FISH data, which was obtained in gray matter regions, our subclustering analysis suggests that most of the astrocytes in our study, and therefore likely most of the resulting rDEGs, arise from protoplasmic or gray matter astrocytes.

### Shared and subtype-specific predicted mechanisms of neuron-astrocyte communication

Many of the astrocyte rDEGs implicated neuron-astrocyte communication, suggesting that the regional molecular identity of astrocytes may arise in part from customized interactions with the vast diversity of specialized neuronal types across the mammalian brain^1,4,64^. Our previous analysis showed that rDEGs are not substantially shared across neurons and glia (**Figs. 2-3D**), which rules out the influence of pan-cell type regional patterning. Neurons and astrocytes communicate via myriad signaling pathways. We assessed whether neuron and astrocyte cluster pairs sampled from the same region were over-enriched for known ligand-receptor (L-R) interactions using CellPhoneDB^65^. To increase the specificity of our predicted L-R results, we restricted CellPhoneDB analysis to neurons and astrocytes only (see **Methods**).

Across most brain regions and ages in the marmoset, we found neurexin and neuroligin (NRXN/NLGN) family members, contactin (CNTN) family members, fibroblast growth factor and receptor (FGF/FGFR) family members, and neural cell adhesion molecule (NCAM) family members to be the most enriched predicted neuron-astrocyte and astrocyte-neuron L-R molecules (**Table S16**). Despite the commonality of these L-R pairs between astrocytes and all neuronal subtypes, for each neuronal subtype in a given region, we found unique or near-unique L-R and R-L pairs with astrocytes. For example, in the fetal marmoset thalamus, *SLT3*→*ROBO2* is specific to midbrain-derived GRIK1+ thalamic inhibitory neurons and immature thalamic astrocytes, while *AFDN*→*EPHA7* signaling is specific to immature astrocytes and TRN GABAergic neurons, compared to the other neuronal subtypes examined (**Fig. 4A**). Later in development, at 14 months, many of the same neuron-astrocyte and astrocyte-neuron L-R combinations were present, while some new pairs, such as *EFNA5*→*EPHB1* for thalamic astrocytes to *GRIK1+* midbrain-derived GABAergic neurons, emerged (**Fig. 4B**, see **Table S17** for quantification of the fraction of L-R pairs shared across 3 or more ages for all neuronal clusters within a region for both species).

**Figure 4.**
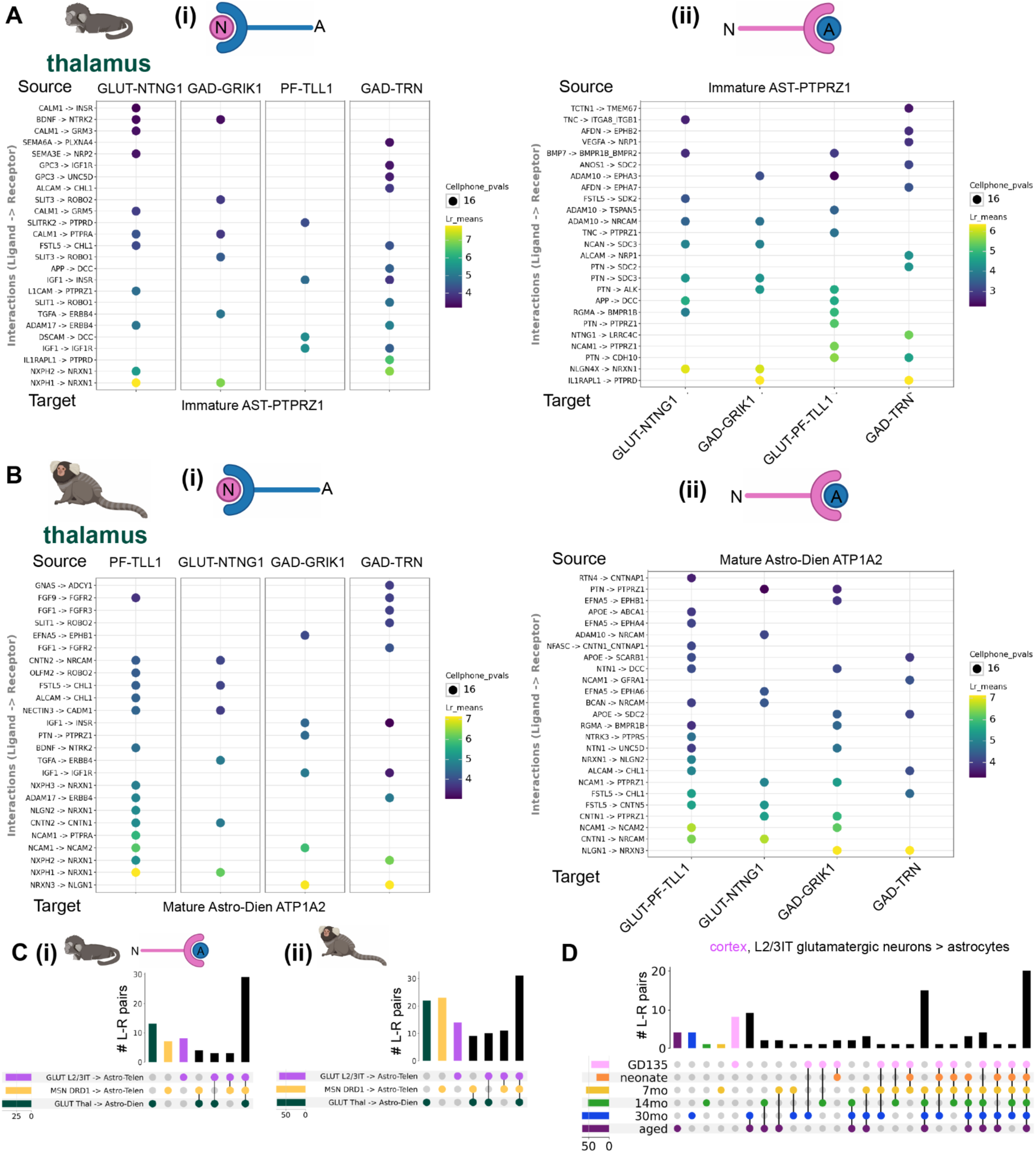
Cell-cell communication analysis for neuron-astrocyte and astrocyte-neuron predicted ligand-receptor pairs across regions and developmental time points in marmoset. **A,** Dot plot showing magnitude and specificity of the top 25 near-unique (shared with at most one other neuronal cluster) CellPhoneDB-predicted **(i)** neuron-astrocyte and **(ii)** astrocyte-neuron ligand receptor pairs for the most abundant astrocyte and neuronal Leiden clusters in the fetal marmoset thalamus. The source cell (top of the plot) expresses the ligand (left side of arrow on the row labels), while the target cell (bottom of the plot) expresses the receptor (right side of arrow on the row labels). The color of the dot indicates ligand-receptor expression magnitude (“Lr_means”, calculated as the average of the mean expression of the ligand in the source group and the mean expression of the receptor in the target group), while the size of the dot is inversely related to the p-value on ligand-receptor expression sensitivity (-log10(p)). **B,** Same as **(A)**, for the 14-month marmoset thalamus. **C,** UpSet plot showing the number of overlapping neuron-astrocyte predicted ligand-receptor pairs between regions, from the most abundant neuronal and astrocyte subtypes in each region for **(i)** fetal and **(ii)** late adolescent marmoset. For cortex (purple), glutamatergic L2/3IT neurons to cortical astrocytes; striatum (yellow), *DRD1+* medium spiny neurons to striatal telencephalic astrocytes; and thalamus (green) thalamic glutamatergic neurons to thalamic astrocytes. The colored dots below each vertical bar indicate which regional neuron-astrocyte (N-A) subtypes share that set of L-R pairs, while the colored horizontal bars indicate the total number of L-R pairs for each N-A subtype. Unlike in panel **A**, all L-R pairs meeting minimum expression criteria, including pairs shared with other neuronal and astrocytic clusters, were included in this analysis and that for panel **(D)**. **D,** UpSet plot (as in **(C)**) showing the number of overlapping cortical glutamatergic L2/3IT neuron to cortical astrocyte predicted ligand-receptor pairs between ages.

To summarize the shared and divergent expression of predicted L-R pairs underlying neuron-astrocyte communication across regions, we examined the overlap of these pairs for the most abundant neuronal cluster and the most abundant astrocyte cluster in cortex, striatum, and thalamus (**Fig. 4C**). We found that many L-R pairs were shared across regions at both GD135 and 14 months (43% and 29% of total L-R pairs, respectively), while the thalamus (at GD135), and later striatum (at 14 months) had the most L-R pairs not shared with other regions for the clusters examined. To exclude the possibility that the region-specificity of neuron-astrocyte L-R pairs is due solely to neuronal heterogeneity, we performed analyses examining the magnitude and specificity of L-R pairs between local (non-projecting) neurons and regional astrocyte populations, the overlap of neuron-astrocyte and neuron-OPC L-R pairs within a region, and the proportion of astrocyte subtype DEGs overlapping with ligand (when source) or receptor (when target) lists compared to neurons. We found that the region-specificity of neuron-astrocyte L-R pairs is not solely explained by neuronal heterogeneity, reflecting a contribution of astrocyte regional heterogeneity. For detailed results, please see the Methods section and associated notebooks in our GitHub repository. In several cases, different members of the same family were used as region-specific neuron-astrocyte/astrocyte-neuron L-R pairs in different regions. For example, in 14-month marmoset, *EFNA5→EPHA5* was unique to cortical astrocyte → cortical L2/3IT glutamatergic neurons, while *EFNA5→EPHA7* was unique to striatal astrocytes *→ DRD1 +* medium spiny neurons, and *EFNA5→EPHA6* was shared across all three regional A→N subtype pairs.

To assess how the expression of L-R pairs underlying neuron-astrocyte communication changes over the course of development, we examined the concordance of L-R pairs of a single neuronal cluster (cortical glutamatergic L2/3IT neuron) and cortical astrocyte at different developmental time points. In contrast to the expression of rDEGs, a larger proportion (20/90, 22%) of L-R pairs (all L-R pairs meeting minimum expression criteria, including pairs shared with other neuronal and astrocyte clusters, were included in this analysis) were shared between all time points (**Fig. 4D**), suggesting that these putative mediators of neuron-astrocyte communication emerge early and are maintained throughout development. However, 15/90 (17%) L-R pairs emerged at 7 months and were maintained throughout adulthood. At later developmental time points, many more L-R pairs were shared between ages than were unique (**Fig. 4D**). This, along with the increased proportion of rDEGs shared across later time points (**Fig. 2C**), suggests that the expression of molecules underlying neuron-astrocyte communication stabilizes postnatally in marmosets at some point between 0 and 7 months.

Overall, patterns of predicted neuron-astrocyte and astrocyte-neuron communication in mouse were similar to marmoset, including implication of neurexin and neuroligin, contactin, fibroblast growth factor and receptor, and neural cell adhesion molecular families (**Fig. S14A-B**, **Table S16**). As in marmoset, mouse thalamus had more unique neuron-astrocyte L-R pairs than cortex or striatum at P4 and P90 (**Fig. S14C**). Unlike in marmoset, mice had more age-specific L-R pairs (from cortical L2/3IT glutamatergic neurons to cortical astrocytes) at earlier time points (particularly at P4, 17/84 or 20% of all L-R pairs unique at this time point) before stabilizing with more shared L-R pairs at later time points. This suggests that mediators of mature neuron-astrocyte interactions emerge relatively later in mouse (**Fig. S14D).** Only 11%of L2/3ITs→astrocyte L-R pairs were shared across the lifespan. Taken together, these result suggest that many L-R pairs potentially underlying neuron-astrocyte communication are shared across developmental time points and regions in both species. However, more neuron-astrocyte predicted L-R pairs emerged later in development.

### Age-dependent refinement of astrocyte identity

In mouse, initiation of gliogenesis in the diencephalon precedes that in the telencephalon by approximately 1 gestational day (E13.5 vs E14.5^66,67^). To determine whether relative immaturity of telencephalic glia compared to diencephalic glia could explain the robust regional expression differences we observed at each sampled time point (**Figs. 2, 3**), we examined the developmental trajectory of astrocytes in pseudotime, a prediction of position along a low-dimensional developmental trajectory based on RNA expression only, using Palantir^68^. Palantir recovered the known developmental trajectory of the oligodendrocyte lineage in both species (**Fig. S15A-B**).^69^ Furthermore, it underscored the precocious myelination in the marmoset brain compared to mouse, as evidenced by a faster rate of pseudotime progression towards maturity and a larger proportion of newly-formed and myelinating oligodendrocytes at earlier time points in marmoset (**Fig. S15C-D**).

In astrocytes from both species, pseudotime analysis with separate terminal states for mature telencephalic (“AST-TE” branch) and diencephalic (“AST-DI” branch) astrocytes revealed a transcriptional developmental trajectory within astrocytes that aligned with actual age and annotation of mature and immature Leiden clusters (**Fig. 5A-B**, **Fig. S15E-F**). Pseudotime values were slightly higher in mature diencephalic versus mature telencephalic astrocytes, and higher in cortical astrocytes than striatal astrocytes, in both species at mature time points, suggesting arrival at distinct terminal states, though these need not be more or less mature than one another (**Fig. 5A-B, Fig. S15E-F**). The rate of maturation (that is, distribution of pseudotime values during development relative to those in adulthood) also differed slightly across regions. To identify genes potentially driving pseudotime transitions, we used Mellon^70^ to calculate gene change scores, a measure of expression change in regions of low cell-state density (see **Methods**), for each pseudotime trajectory branch. Many of the top 25 change-scoring genes in both branches were cortex-thalamus rDEGs (**Fig. S15C-D**, **Table S18**), suggesting that the expression of region-enriched genes is correlated with, and potentially drives, astrocyte maturation.

**Figure 5.**
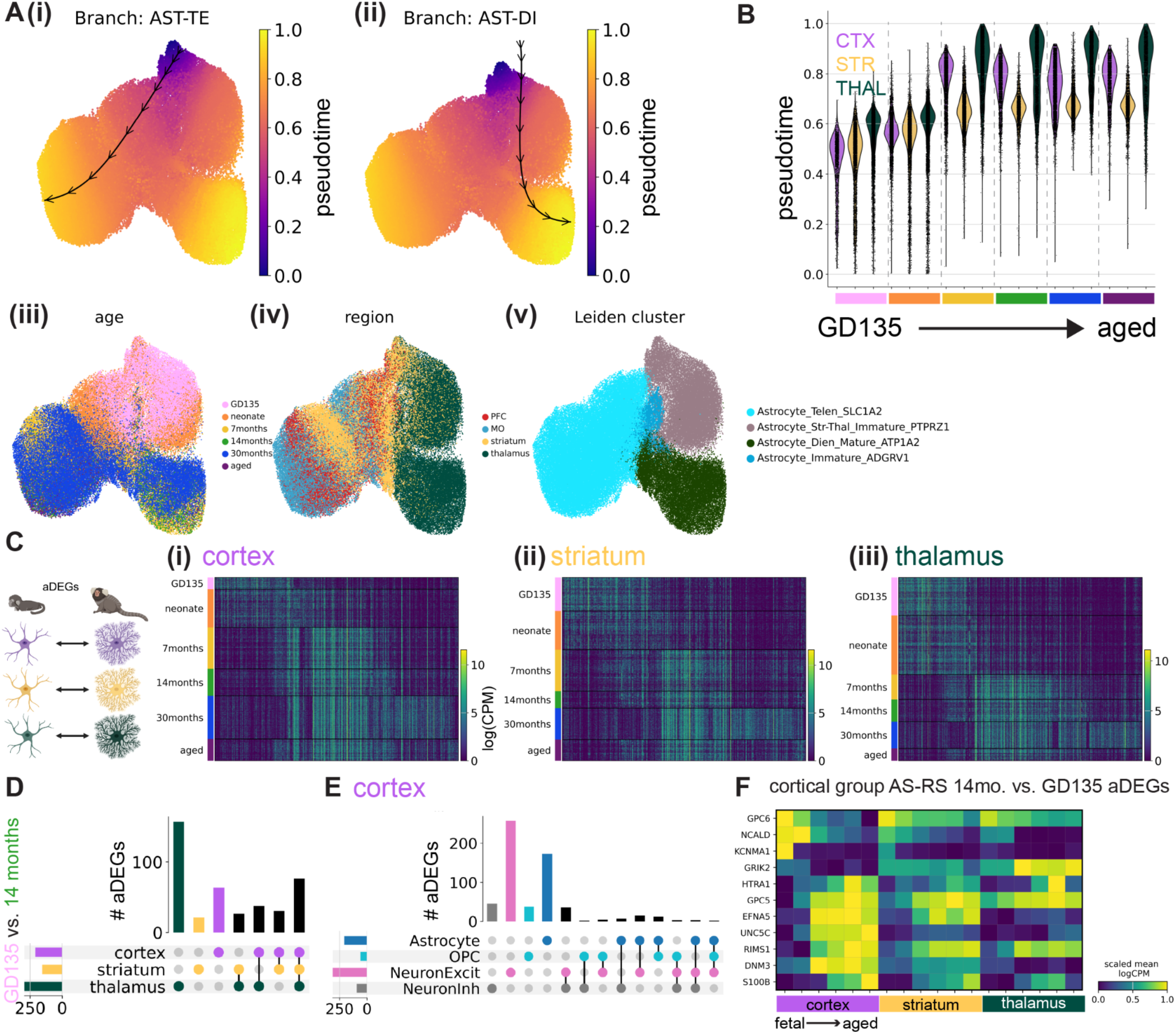
The postnatal developmental specification of marmoset astrocytes within and across brain regions. **A,** Integrated UMAP embeddings of 103,009 marmoset astrocytes colored by **(i-ii)** Palantir-predicted pseudotime, **(iii)** developmental time point, **(iv)** brain region (assigned), and **(v)** Leiden cluster assignment. Trajectory path (black lines and arrows) are overlaid for **(i)** telencephalic astrocyte (AST-TE) and **(ii)** diencephalic astrocyte (AST-DI) branches. **B,** Violin plot (scanpy’s default) showing the estimated distribution of pseudotime values for the astrocytes in **(A)** grouped by region within each developmental time point (color code as in **A(i)**). Vertical dashed lines indicate separation between time points. **C,** Heatmaps (rows corresponding to nuclei and columns to gene) showing expression in logCPM of astrocyte age differentially expressed genes (aDEGs) in astrocytes from **(i)** cortex, **(ii)** striatum, and **(iii)** thalamus, grouped by developmental time point as indicated on the left of the heatmap. The strategy for calculating aDEGs is schematized on the left. **D,** UpSet plot showing the number of overlapping GD135 vs. 14-month astrocyte aDEGs between cortex, striatum, and thalamus. The colored dots below each vertical bar indicate which region(s) share that set of aDEGs, while the colored horizontal bars indicate the total number of cortex-thalamus aDEGs for each region. Overlap categories with 0 aDEGs are not shown. **E,** UpSet plot (as in **(D)**) showing the number of overlapping GD135 vs. 14-month cortical astrocyte aDEGs between OPCs (light blue), astrocytes (dark blue), excitatory neurons (pink), and inhibitory neurons (gray). **F,** Matrix plot showing mean expression of selected cortex group astrocyte-specific, region-specific (AS-RS) aDEGs (rows) in marmoset astrocytes grouped by region and developmental time point (columns, blocked by region first and then by increasing age within each region block). Expression units of mean logCPM are standardized between 0 and 1 by subtracting the minimum and dividing by the maximum for each trait.

Next, for each region we binned marmoset astrocytes by pseudotime quintile in the appropriate trajectory branch rather than by actual age and recomputed rDEGs. As with rDEGs grouped by actual age, the number of rDEGs increased from pseudotime bin 1 (PT1) to pseudotime bin 5 (PT5): 54 rDEGs at PT1, 108 at PT2, 117 at PT3, 177 at PT4, and 181 at PT5. We found that matching by predicted maturational stage largely recapitulated the original rDEGs calculated from actual age: 56% of PT1 cortex-thalamus rDEGs overlapped with GD135 rDEGs, 81% of PT2 rDEGs overlapped with neonate rDEGs, 39% of PT3 rDEGs overlapped with 7-month rDEGs, 85% of PT4 rDEGs overlapped with 14-month rDEGs, 81% of PT5 rDEGs overlapped with adult rDEGs, and 85% of PT5 rDEGs overlapped with aged rDEGs. This suggests that regional imprinting of astrocytes is not simply driven by relative differences in the birth timing of cells across the different brain structures.

We next sought to determine the sequence of molecular changes that unfold in astrocytes within a given region over time. We calculated **a**ge **d**ifferentially **e**xpressed **g**enes (aDEGs) within each brain region from metacells of each age (see **Methods**). In each brain region, there were over 100 unique aDEGs (unique after pooling pairwise age combinations). In marmoset, the largest fraction of aDEGs distinguished GD135/neonate from 7-month and older astrocytes (**Fig. 5C**). aDEGs enriched in the 30-month dataset could conceivably arise from the different sample preparation and reference genome used in our previous study^7^; for this reason we used the 14-month time point to further assess age-related changes across regions.

Examining the overlap of marmoset astrocyte GD135 vs. 14-month aDEGs between brain regions (409 aDEGs in total), we found that ∼19% were shared between cortex, striatum, and thalamus (**Fig. 5D**). The striatum had a modest number of GD135 vs. 14-month aDEGs not shared with other regions (21/409), while the cortex had 3-fold more (63/409), and the thalamus had the most (156/409), as expected given the stark regional heterogeneity between telencephalon and diencephalon (**Fig. 2**). Additionally, we found that few to no GD135 vs. 14-month aDEGs were shared between astrocytes, OPCs, and excitatory neurons or astrocytes, OPCs, and GABAergic neurons in the cortex (**Fig. 5E**).

We found similar results in mice, where we calculated P4 vs. P90 rDEGs, as mouse E18.5 astrocytes were transcriptionally immature relative to marmoset GD135 astrocytes and GD135-P4 timepoints appear to have better correspondence, as discussed in the next section). One notable difference from marmoset was that early adolescent (P14) astrocytes in mice expressed many aDEGs shared with embryonic and neonate timepoints, particularly in striatum (**Fig. S16A**). As in marmoset, thalamic astrocytes had more unique aDEGs than their cortical and striatal counterparts (**Fig. S16B**), and most astrocyte aDEGs were not shared with other radial glia-derived cell types (**Fig. S16C**).

We found very few (3 or less) astrocyte aDEGs that were cell type-agnostic and region-specific (i.e., that were also aDEGs in neurons and OPCs for a given brain region, see **Methods**). In contrast, there were 74 (marmoset GD135 vs. 14-month) and 56 (mouse P4 vs. P90) aDEGs that were astrocyte-specific and region-agnostic, reflecting more universal aspects of astrocyte transcriptional maturation, regardless of brain region. We found a similar number of astrocyte-specific, region-specific aDEGs (20 in striatum, 51 in cortex, and 135 in thalamus for GD135 vs. 14-month marmoset), which reflect the brain region’s influence on the maturation of astrocytes *only* in a given brain region. In both species, the developmental pattern of selected astrocyte-specific, cortex-specific aDEGs was similar but not identical in the striatum, and more dissimilar with the thalamus (**Fig. 5F**, **Fig. S16D**).

### Conservation and divergence of astrocyte patterning in mouse and marmoset

Hundreds of differentially expressed genes distinguish adult human and mouse astrocytes^71^, and engrafting human glial progenitors into mouse brain results in mature astrocytes that retain certain human-specific astrocyte characteristics^72^. This suggests that aspects of an astrocyte’s developmental program are cell intrinsic and are shaped by its species-specific genomic features. We therefore aimed to compare transcriptional signatures of telencephalic and diencephalic regional astrocyte populations between marmoset and mouse. We integrated a randomly downsampled subset (100,000 nuclei each) of mouse and marmoset nuclei (all cell types included). To do so, we used 547 highly variable 1:1 ortholog genes selected from top differentially expressed genes of superclusters (related groups of Leiden clusters) shared across species and employed the semi-supervised variational auto-encoder scANVI^73^ (see **Methods**). The resulting integrated UMAP plot showed broad conservation of superclusters between mouse and marmoset, despite differences in cell type proportions across development (**Fig. S17A-B**). Another method called SATURN^74^, which avoids 1:1 mapping of genes, had largely concordant cross-species integration results (**Fig. S17C-D**).

Species-integrated astrocytes partitioned into three superclusters that segregated by developmental stage and by brain structure (diencephalon vs telencephalon) (**Fig. 6A**). This indicates that at the level of broad cephalic domains, region patterning is conserved between the two species. Mature telencephalic astrocytes showed better species integration than diencephalic or immature astrocytes, and immature mouse astrocytes composed a distinct cluster (**Fig. 6A**). This finding implies that at birth, mouse astrocyte maturity lags behind that of marmoset, in line with our findings about oligodendrocyte maturation (**Fig. S13A-B, G-H**). Interestingly, marmoset astrocyte aDEGs had more discrete expression boundaries across time (**Fig. 5C**), while mouse astrocyte aDEGs had more continuous temporal expression, especially in the striatum (**Fig. S16**). Additionally, marmoset rDEGs shared across ages were largely divided into younger (GD135, neonate) and older (7 months and older) groups (**Fig. 2C**). In contrast, temporally overlapping mouse rDEGs were more evenly distributed across individual ages and smaller groups of ages (**Fig. 3C**), suggesting that developmental changes occur more slowly over the sampled timepoints in mouse.

**Figure 6.**
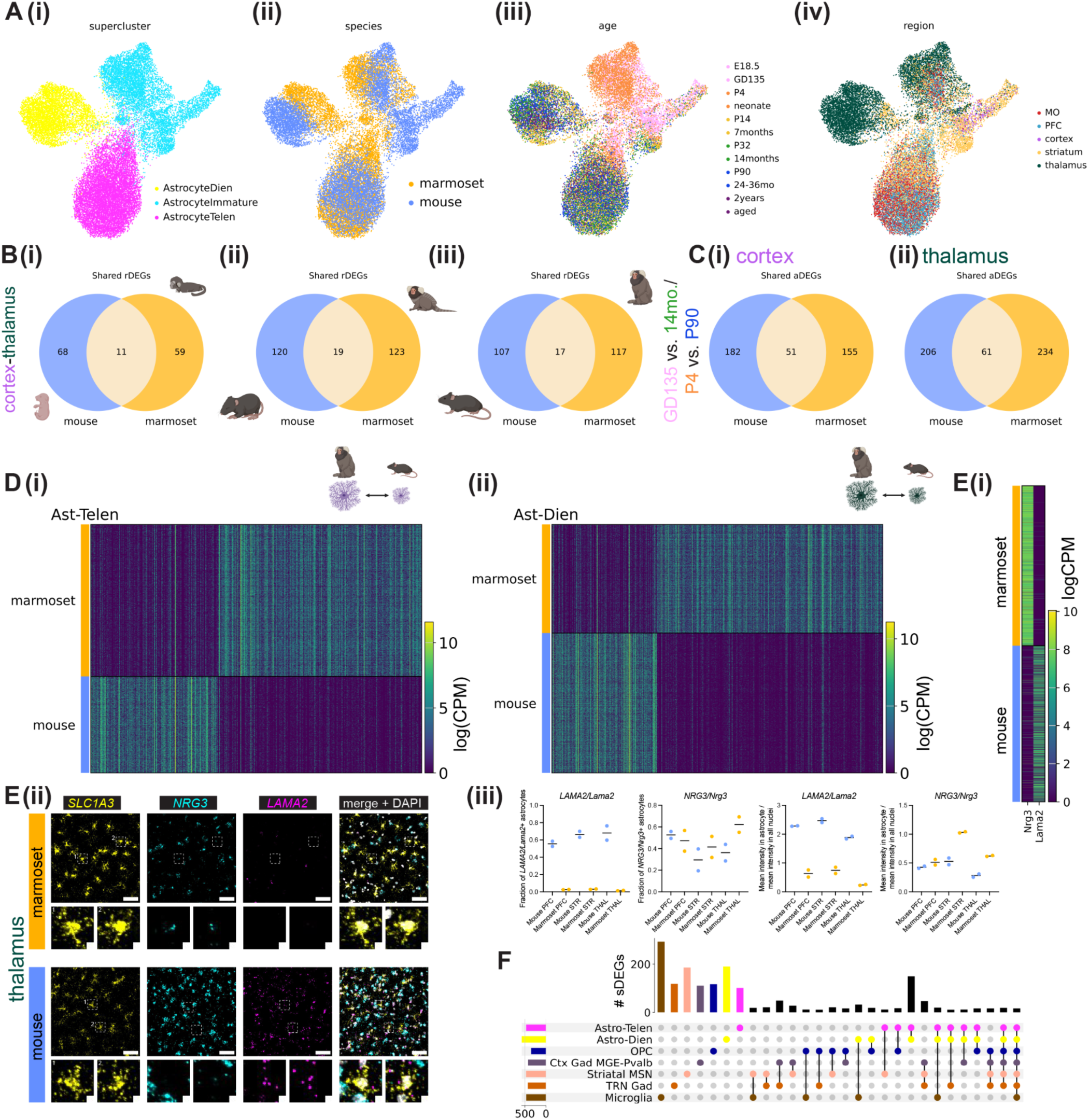
Conservation and divergence of the development of astrocyte heterogeneity in mouse and marmoset. **A,** scANVI-integrated UMAP embeddings of marmoset and mouse astrocytes, colored by **(i)** supercluster, **(ii)** species, **(iii)** age, and **(iv)** region. **B,** Venn diagrams showing regional differentially expressed genes (rDEGs) between cortex and thalamus astrocytes shared across mouse and marmoset at **(i)** fetal, **(ii)** early adolescent, and **(iii)** young adult time points. **C,** Venn diagram showing age differentially expressed genes (aDEGs) shared between mouse and marmoset astrocytes within the **(i)** cortex and **(ii)** thalamus. **D,** Heatmaps showing expression in logCPM of species differentially expressed genes (sDEGs) between marmoset and mouse within **(i)** telencephalic astrocytes and **(ii)** diencephalic astrocytes. E, *In situ* validation of selected sDEGs in marmoset and mouse tissue with the RNAscope v2 assay. **(i)** Heatmap (rows are nuclei, columns are genes) of sDEGs *NRG3*/*Nrg3* (higher in marmoset) and *LAMA2*/*Lama2* (higher in mouse) expression in marmoset and mouse astrocytes in logCPM. **(ii)** Top row: single-channel and composite maximum intensity projections of cropped fields of view in the marmoset (top row) and mouse (bottom row) thalamus stained via RNAscope v2 FISH for astrocyte marker *SLC1A3/Slc1a3*, *NRG3/Nrg3*, and mouse *LAMA2/Lama2*. Scale bar, 50µm. Bottom row: high-magnification images of the boxed astrocytes in the top row. Scale bar, 5µm. **(iii)** CellProfiler quantification of sDEG abundance in both species from RNAscope v2 data (n = 2 donors per species, see **Methods**). From left to right: Fraction of probe positive astrocytes for *LAMA2*/*Lama*2, fraction of probe positive for *NRG3*/*Nrg3*, normalized mean intensity of *LAMA2/Lama2* (mean intensity in expanded astrocyte nuclei divided by mean intensity in all nuclei, including astrocyte nuclei), and normalized mean intensity of *NRG3/Nrg3*. Data points are from individual biological replicates, with 2 slices averaged for mouse, and the horizontal black line denotes the median. **F,** UpSet plot showing shared sDEGs across superclusters, comparing telencephalic astrocytes, diencephalic astrocytes, oligodendrocyte precursor cells (OPCs) medial ganglionic eminence-derived and *PVALB*+ cortical GABAergic neurons (Ctx Gad MGE-PVALB), thalamic reticular nucleus (TRN) GABAergic neurons, striatal medium spiny neurons (MSNs), and microglia. The colored dots below each vertical bar indicate which supercluster(s) share that set of sDEGs, while the colored horizontal bars indicate the total number of mouse-marmoset sDEGs for each supercluster. For simplicity, only supercluster combinations with 10 or more shared sDEGs are shown.

In mouse but not marmoset, we observed immature astrocyte clusters composed of nuclei from all time points, suggesting continued generation of new astrocytes throughout the lifespan. These included the *Top2a+* immature astrocyte population seen in the neurogenic subventricular zone throughout the lifespan (**Fig. S18**), which forms part of the rostral migratory stream.^75^ All marmoset astrocytes and cortical and thalamic mouse astrocytes exhibited separate embryonic and neonatal subclusters from early adolescent and older cells (**Fig. S12-13A, C**). In contrast, in the mouse striatum, there were several more immature clusters (9 total), including some composed of astrocytes from mature timepoints (**Fig. S13B**, “Str_Ast2”, “Str_Ast6”, and “Str_Ast12”). However, the relative immaturity of mouse astrocytes after adolescence is not driven solely by the persistence of *Top2a+* cells in the SVZ, as we found an immature population of *Top2a*-striatal astrocytes in mouse but not marmoset that may reside outside of the SVZ (**Fig. S13B,** portions of “Str_Ast2” and “Str_Ast12”). We note that the presence of this immature cluster in the striatum does not suggest that all mouse astrocytes are less mature than their marmoset counterparts in adolescence and adulthood.

We next tested whether genes that best distinguished astrocytes from a given brain region in one species were more likely than chance to be rDEGs in the other species. Focusing on astrocyte cortex-thalamus rDEGs at each developmental time point, we found that the majority of rDEGs were not shared across species, and that the proportion of shared rDEGs decreased only slightly from fetal to early adolescence time points, from ∼14-16% to ∼13-14% in both species (**Fig. 6B**, full list of species-overlapping and species-unique cortex-thalamus astrocyte rDEGs at each developmental time point in **Table S21**). The proportion of overlapping rDEGs was not greater than chance (see **Methods**, p-value from a Fisher’s exact test > 0.05 at all developmental time points).

Similarly, the majority of astrocyte aDEGs in cortex and thalamus were not shared between species. 51 cortical aDEGs (22% of mouse cortical P4-P90 aDEGs and 25% of marmoset GD135-14-month aDEGs) and 61 thalamic aDEGs (23% of mouse thalamic P4-P90 aDEGs and 21% of marmoset thalamic GD135-14-month aDEGs) were shared between species (**Fig. 6C**, p-value from a Fisher’s exact test on the proportion overlapped = 0.025 for cortex and 0.454 for marmoset). Lists of species-shared and species unique aDEGs by region are provided in Table S22.

Next, we directly tested for differential expression of 1:1 orthologs between species within shared superclusters. We calculated **s**pecies **d**ifferentially **e**xpressed **g**enes (sDEGs) based on 1:1 orthologs between species within each integrated supercluster using our metacell method (see **Methods**). We found hundreds of sDEGs in both telencephalic (464 total) and diencephalic (579 total) astrocytes whose expression could clearly distinguish between marmoset- and mouse-derived populations (**Fig. 6D**). Astrocyte sDEGs encoded both cytosolic and membrane-bound proteins with varied cellular functions (**Table S24**). For example, telencephalic astrocyte sDEGs higher in marmoset included *NALCN/Nalcn*, a non-selected sodium leak channel; the RNA-binding protein *RBFOX2/Rbfox2*; *KCNT2*/*Kcnt2*, a sodium-activated potassium channel subunit; *FABP7/Fabp7*, a fatty acid binding protein with established roles in neurogenesis; and *DNM3/Dnm3*, a multi-domain GTPase involved in membrane remodeling. Even this short list of genes suggests that important cellular functions such as ion buffering, RNA processing, fatty acid binding, and membrane remodeling may differ between astrocytes of different species. Furthermore, 63/286 telencephalic and 71/395 diencephalic sDEGs were SFARI^76^ 3.0 Autism Spectrum Disorder (ASD)-related genes (see **Methods**). These overlaps are significantly higher than chance (assuming 20,000 protein-coding genes in the human genome, p-values from Fisher’s exact tests < 10^-15^ for both telencephalic and diencephalic astrocyte sDEGs). Complete lists of sDEGs for each supercluster analyzed, and GO annotations for Cellular Compartment, Biological Process, and Molecular Function are provided in **Table S23.**

We validated the differential expression of 2 astrocyte sDEGs, *NRG3/Nrg3* (present in neurons in both species but higher in marmoset astrocytes) and *LAMA2/Lama2* (higher in mouse astrocytes) *in situ* using RNAscope (**Fig. 6E**). 50% of the genes that distinguish diencephalic astrocytes between species were shared with telencephalic astrocytes (**Table S24**). These telencephalic-diencephalic astrocyte shared sDEGs made up a larger fraction (62%) of telencephalic astrocyte sDEGs. This suggests that evolution has acted on the astrocyte class as a whole, while also shaping divergent regional astrocyte programs between species. Additionally, as with astrocyte rDEGs and aDEGs, we found that most astrocyte sDEGs were not shared with other cell types (superclusters), including OPCs, cortical MGE-derived *PVALB*+ interneurons, GABAergic TRN neurons, striatal MSNs, and microglia (**Fig. 6F**). This result underscores that evolutionary divergence of a cell type’s transcriptome unfolds at different rates across cell types^62,77,78^. Taken together, these findings support both conservation and divergence of postnatal astrocyte regional specialization in mouse and marmoset.

### Astrocytes have regionally divergent morphology and protein expression

Many of the genes we found to vary in astrocytes by region, age, and species implicate processes involved in morphological specification. Indeed, astrocyte morphology, which is highly ramified and complex, is essential for their specialized functions: end feet contact blood vessels to help form the blood-brain barrier and shuttle water and nutrients, while terminal processes closely appose synapses to uptake ions and neurotransmitters^35^ and regulate synapse development and function^79^. Because many of these morphological features exist at the sub-micron scale, conventional light microscopy is not sufficient to visualize the full morphological complexity of astrocytes^36^. We wondered whether nanoscale astrocyte morphology might also be regionally specialized between gray matter regions in cortex, striatum, and thalamus, as recently demonstrated for several CNS regions using diffraction-limited approaches^25^. Thus, we used expansion revealing (ExR), a new variant of protein decrowding expansion microscopy^37^, to visualize astrocyte morphology with enhanced resolution and compare morphological properties in the PFC, striatum, and thalamus, as these regional populations represented the major molecular subpopulations in our snRNAseq data (**Fig. 2, 3**).

We used a viral approach to label astrocytes for expansion (**Fig. 7A-B**, see **Methods**) and created 3D binary segmentations to quantitatively assess morphological differences across regions (**Fig. 7D**, Movies **S19-36**). We calculated the volume, surface area, equivalent diameter (measures of size), surface area to volume ratio (S:V, a measure of shape, inversely proportional to size), aspect ratio (a measure of elongation), fractal dimension (FD)^80^ (a measure of complexity and self-similarity), and branching complexity via Sholl analysis^81^, most of which have been used to characterize astrocyte morphology in prior studies^36,82^.

**Figure 7.**
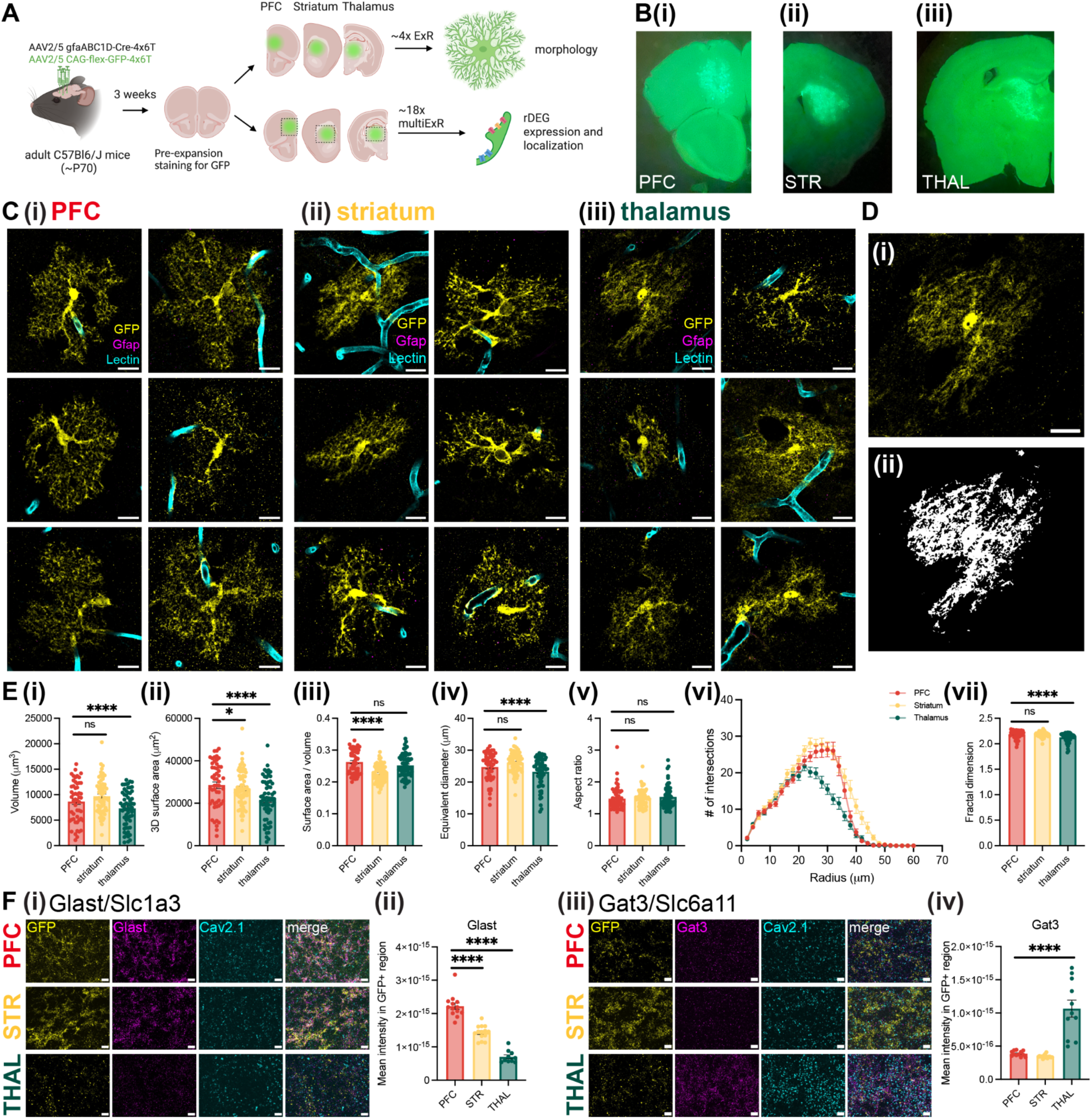
Expansion microscopy of virally-labeled astrocytes in the mouse and marmoset brain. **A,** Viral labeling approach for mouse astrocytes (see **Methods** for details). Created using BioRender.com. **B,** Brain slice hemispheres containing regions of interest in **(i)** PFC **(ii)** striatum and **(iii)** thalamus after pre-expansion staining for GFP, as visible through the eyepiece on a dissecting microscope under blue light illumination. **C-E, (i)** Single z-slices of background-subtracted images of ∼3.5x expanded astrocytes in the **(i)** prefrontal cortex, **(ii)** striatum, and **(iii)** thalamus, co-stained with GFP, GFAP, and the blood vessel marker Lectin. Shown are 6 examples of medium to high GFP-expressing astrocytes from 6 separate mice. Scale bar, 10µm in biological units. Contrast was adjusted to 35% saturation (Fiji’s “auto”) in most cases and was further increased for dimmer astrocytes to aid visibility; therefore, astrocytes are not equally contrast adjusted. See Movies S1-36 for 3D-projected image volumes and their corresponding 3D segmentations. **D, (i)** Single z-slice of one of the thalamic astrocytes in **C(iii)** and **(ii)** its corresponding binary segmentation used as input for morphology analysis. **E,** Bar plots showing quantified morphological properties for mouse astrocytes from PFC, striatum, and thalamus (n = 52-60 astrocytes from 3 female and 5 male mice for each region, with statistical significance determined using a linear mixed effects model with “animal” as the random effect group variable, see **Table S25**): **(i)** volume, **(ii)** surface area, **(iii)** surface area / volume, **(iv)** equivalent diameter, **(v)** aspect ratio, **(vi)** Sholl analysis (number of intersections with concentric shells as a function of radius), and **(vii)** fractal dimension by the box counting method (see **Methods**). ns, non significant, *p≤0.05, **p≤0.01, ***p≤0.001, ****p≤0.0001, **F,** Maximum intensity projection composites of cropped (in x and y and across 51 z-slices) regions of GFP-labeled astrocyte processes co-stained with either **(i)** Glast (telencephalic rDEG) or **(iii)** Gat3 (thalamic rDEG) and the synaptic protein Cav2.1 alongside GFP generated using the ExR protocol (∼18x expansion factor). Shown are processes from astrocytes in the PFC (top row), striatum (middle row), and thalamus (bottom row) of the same mouse (though **(i)** and **(iii)** are from different mice). Scale bar, 0.5µm. Contrast was manually adjusted by setting equal minimum and maximum intensity values for Cav2.1 and rDEG targets and using Fiji’s “Auto” for GFP. **(ii, iv)** Bar plots showing quantified mean intensity of either Glast or Gat3 within masked GFP+ regions (astrocytes) in the whole ExR image volume (n = 10-16 fields of view from 2 mice, with statistical significance determined using a linear mixed effect model as described in the **Methods, see Table S26 for numerical results**). In panels **E** and **F**, error bars indicate standard error of the mean.

After correcting for multiple comparisons across the 6 univariate measures, we found significant differences in size (volume, surface area, and equivalent diameter), shape (surface area to volume ratio), and morphological complexity (FD) between astrocytes from different brain regions, particularly between cortical/striatal and thalamic astrocytes (**Fig. 7E**, n = 52-60 astrocytes from 3 female and 5 male mice for each region, with statistical significance determined using a linear mixed effects model with “animal” as the random effect group variable, see **Table S25**). Specifically, thalamic astrocytes were smaller and less complex compared with cortical and striatal astrocytes, while striatal astrocytes had smaller surface area and surface area to volume ratios compared to cortical astrocytes. To exclude the possibility that proximity to under-digested blood vessels or incomplete capture in the axial dimension impacted these differences, we repeated this analysis on a subset of astrocytes meeting additional criteria and found largely similar results (**Fig. S19A**, see **Methods**). Similarly, Sholl analysis revealed fewer intersections at radii larger than ∼25 µm for thalamic compared to cortical and striatal astrocytes (**Fig. 7E(vi)**). Taken together, these results suggest that thalamic astrocytes are smaller and less morphologically complex than their cortical and striatal counterparts, in line with prior work using conventional microscopy^25^.

We next probed rDEG protein product expression level and localization in astrocytes at the nanoscale using ExR. We processed tissue from 2 of the virally-labeled adult C57 Bl/6J mice used for morphology analysis and proceeded with staining for Glast (encoded by the telencephalic rDEG *Slc1a3*) and Gat3 (encoded by the thalamic rDEG *Slc6a11*) alongside GFP (astrocyte processes) and Cav2.1 (a presynaptic protein) in gels prepared from PFC, striatum, and thalamus. We found that the expression of Glast and Gat3 at the protein level in GFP-labelled astrocytes agreed with snRNAseq predictions: Glast expression was highest in the cortex, lower in the striatum, and lowest in the thalamus, while Gat3 expression was higher in the thalamus than cortex or striatum (**Fig. 7F**, **Fig. S19B**). Both proteins localized to astrocyte processes near synapses. Taken together, these results support the differential expression of rDEG protein products between telencephalic and diencephalic astrocytes, and reveal the localization of rDEG protein products on and near astrocyte processes, in close proximity to neuronal synapses.

## Discussion

Astrocytes are a ubiquitous, versatile brain cell type with increasingly appreciated roles in health and disease. While their regional molecular heterogeneity has been evident for some time^83,84^, the source of this regional heterogeneity, in particular, the relative contributions of embryonic patterning versus response to environmental cues after birth, is not well understood^85^. To help bridge this knowledge gap, we generated a unified, multi-region, postnatal developmental snRNAseq atlas of mouse and marmoset brain cells. Because our dataset contains all brain cell types, we anticipate this atlas will be a valuable resource for the field. As such, we have made both raw and processed data publicly available on NeMO and the Broad Single Cell Portal, respectively (see **Data and Code Availability**). The latter is useful for exploring cell type clusters and querying the expression pattern of genes of interest across ages and regions, and does not require coding expertise.

We found that astrocytes were regionally patterned before birth in both species, a discovery that was not unexpected given the prevalence of homeobox patterning genes among astrocyte regionally differentially expressed genes^8^, evidence from a recent study showing regionally patterned glioblasts in the first-trimester human brain^6^, and older lineage tracing showing regional allocation of astrocytes based on the region of their originating radial glia^27^. Less predictably, we discovered dramatic changes in astrocyte regional identity between birth and early adolescence, in line with their maturation during this period. This period also coincides with peak synaptogenesis, pruning, and myelination^86^, consistent with the notion that astrocyte specialization depends on the activity of neighboring cells^87^.

The functions of embryonically-patterned and postnatally-acquired astrocyte rDEGs were varied, but implicate known astrocyte processes, including supporting synaptic transmission, ion transport, neurotransmitter uptake, cell-cell adhesion, and morphological specification. The function of some rDEGs, including *SLC6A11* (GAT-3) and *SPARC*, has been studied in astrocytes, and shown to be important in modulating the effects of brain injury^88^ and controlling synaptogenesis^53^, respectively. We anticipate future mechanistic studies of other astrocyte rDEGs will reveal yet more essential functions.

We found that neuron-astrocyte and astrocyte-neuron predicted ligand-receptor pairs, many of which were specialized for distinct neuronal subtypes, were upregulated during postnatal development into adulthood, again supporting the hypothesis that astrocytes specialize in postnatal development to meet the needs of local neurons. Despite the striking regional heterogeneity of astrocytes, many predicted neuron-astrocyte ligand-receptor pairs were shared across regions. Even those not shared across regions were functionally similar, suggesting neurons and astrocytes have developed a common language of molecular communication across the forebrain. Indeed, some of our rDEGs were members of the same protein family or functional class, pointing to variations on a common theme of neuron-astrocyte crosstalk across brain regions. Many of the top predicted neuron-astrocyte ligand-receptor pairs, such as neurexins and neuroligins, are more traditionally associated with neuron-neuron contact at the synapse^89^. However, adhesion molecules such as ephrins, neurexins/neuroligins, and NrCAMs have been shown to play important roles in neuron-astrocyte communication^49^.

In both species, we found hundreds of age differentially expressed genes (aDEGs), many of which astrocyte-specific but region-agnostic, some of which were astrocyte-specific and region-specific, and very few of which were cell type-agnostic but region-specific. The thalamus had the most unique astrocyte developmental gene expression signature of the three brain regions, suggesting that thalamic astrocytes undergo distinct developmental changes from their telencephalic counterparts. Our astrocyte-specific, region-agnostic aDEGs can be interpreted as a core forebrain astrocyte developmental program, and were more likely to be shared across the species. For example, in marmoset, this included *NTRK2*, which encodes the BDNF receptor TrkB, the short isoform of which has been shown to be essential for astrocyte morphogenesis^90^. Perhaps unsurprisingly, several of our region-specific aDEGs were also rDEGs, and/or had a high degree of functional overlap with rDEGs. While pseudotime approaches have limitations^91^ and may not fully capture how maturational states differ across brain regions, they can provide information about the progression of change along a trajectory that is correlated with actual age. For this reason, we used pseudotime to compare relative maturation differences across regions. This analysis suggested that intrinsic maturation rates are relatively low drivers of regional differences in gene expression.

In both species, most astrocyte rDEGs, aDEGs, and sDEGs were not shared with OPCs or neurons, suggesting that astrocyte region- and age-specializations are unique, rather than general to all radial-glia derived cell types in the same region, developmental time point, or species. This suggests either that regional gene expression signatures change throughout neuro- and glio-genesis, or that the downstream transcriptional effects of this early regional patterning depend on the daughter cell’s fate. Evidence for both exists in the cortex^92^. Why neurons and astrocytes exhibit stark regional patterning in adulthood, albeit in different ways, while the oligodendrocyte lineage does not, is an outstanding question for future study.

The present study characterized two mammalian neuroscience model species, mouse and marmoset. While mice and humans have a high degree of genetic conservation^93^, mice have certain limitations as a model for studying the human brain including lack of a well-developed prefrontal cortex and complex social behaviors, and poor visual acuity. In light of these limitations, non-human primates, with whom we share much closer genetic ancestors, are considered as more translationally-relevant models of brain function and dysfunction. The common marmoset has become an increasingly popular non-human primate model in neuroscience studies due to its faster generation time for genetic engineering, shorter lifespan than other larger primates for developmental and late-onset disease studies, and complex social behaviors^94^.

Our data suggest that the development of astrocyte regional heterogeneity, marked by embryonic regional patterning along cephalic boundaries followed by dramatic postnatal specialization, is broadly conserved between mouse and marmoset. However, the expression of many rDEGs and aDEGs differs across species, and we identified hundreds of species differentially expressed genes within both telencephalic and diencephalic astrocytes (**Fig. 5**). These sDEGs encode proteins involved in key cellular functions which may have undergone evolutionary selection, and a significant portion have been associated with ASD. Future studies exploring the function of these sDEGs within astrocytes may reveal how primate astrocytes have evolved to suit the unique anatomy and physiology of the primate brain. Taken together, these findings suggest that each species may have evolved by recruiting different sets of genes that facilitate postnatal regional specialization of astrocytes. We found that many cell types in the marmoset brain are transcriptionally more mature at time of birth than their mouse counterparts, in line with previously documented precocious development in early postnatal marmosets^95^. This species divergence in transcriptional maturity at time of birth suggests that researchers should use caution when comparing early postnatal time points between rodents and NHPs, especially in light of differences in developmental tempo between species^96^.

We used expansion revealing (ExR) combined with a viral astrocyte labeling approach to circumvent the diffraction limit of light microscopy and the limitations of immunostaining, respectively, to visualize astrocyte processes with enhanced resolution. Our quantitative profiling of astrocyte morphology in the mouse brain shows that astrocyte size, shape and complexity do vary across brain regions, and are most distinct between thalamus and cortex. Prior studies have also found appreciable morphological differences in mouse astrocytes across brain regions^19,25,97^. Although other higher resolution approaches such as electron microscopy may reveal additional differences in aspects of astrocyte morphology, we demonstrate that expansion microscopy, particularly ExR, offers an inexpensive and accessible alternative to other super-resolution approaches for characterizing astrocyte morphology with enhanced resolution. We anticipate that this approach could also be used to study morphological changes in astrocytes after manipulation and/or in disease contexts.

There are several notable limitations to the current study, only some of which we discuss here. The first is the reliance on 10x Chromium snRNAseq, which is subject to dropout and 3’ bias, and produces short reads that cannot be used to map splice variants or many single-nucleotide polymorphisms that may differ between cell types and species. Additionally, the use of nuclei instead of whole cells prevents our detection of RNAs in the cytoplasm, including those locally translated in distant processes of both neurons^98^ and astrocytes^99^, which are likely relevant in establishing cell type and state identity. However, others have found similar cell type discrimination capabilities for single-cell and single-nucleus RNAseq in the mouse cortex, despite lower RNA content (20-50% of total cellular mRNA) in single nuclei.^100^ The second significant limitation is the relatively small sample size, especially for marmosets due to practical limitations including cost, which limits our ability to compare between sexes.

Third, we relied in part on pathway analysis to summarize patterns and deduce functional implications arising from sets of rDEGs. Our use of WebGestalt did not incorporate any fold-change or p-value information for genes, treating each DEG equally regardless of its differential expression level, which may skew results. Furthermore, pathway analysis is only as accurate as the underlying annotations, which can be lacking for glial biology. Finally, many genes are involved in several pathways. For these reasons, we encourage interested readers to directly examine our DEG lists provided in the **Supplementary Tables**.

Fourth, there are significant challenges in integrating snRNAseq data across species. Even before data analysis, read alignment will differ across species, varying with the quality and content of reference genome annotation (for example, we used an optimized version of the mm10 reference genome^101^, for which no analogous version exists for the marmoset). During data analysis, approaches requiring direct merging of the cell x gene count matrices (as in **Fig. 6**) results in the loss of biological information, because only roughly 50-60% of total genes detected in either species were mapped as one-to-one orthologs. For this reason, we also integrated the data across species with an orthogonal approach that does not rely on one-to-one ortholog mapping (**Fig. S17C**). Both approaches are further limited by the need for *a priori* cell type annotation, which may bias towards or against integration of shared and unshared superclusters, respectively. Therefore, we are most confident in sDEGs, which are all one-to-one orthologs and calculated within shared superclusters, as a measure of species divergence.

Fifth, our approach for labeling astrocytes for morphological analysis relies on viral infection and manual identification of astrocytes meeting a minimum brightness level for imaging and segmentation, which may be biased towards a certain astrocyte subtype. Finally, we relied on RNAscope HiPlex to assess rDEG mRNA levels *in situ*. Any multiplexing technique that involves repeated stripping and restaining suffers from some level of reduced fluorescence intensity in later rounds, as well as some amount of registration error. Therefore, any researcher interested in following up on a gene or protein of interest that was imaged in a later round should perform additional confirmatory studies with a single round of imaging.

Taken together, our data support a model of astrocyte regional specialization that includes both embryonic patterning and postnatal specialization in response to local environmental cues, including synapse formation and neuronal activity, as has been previously suggested^25,84^. To determine whether or not early transcriptional patterning is required for proper postnatal astrocyte specialization for such a role, a cross-region astrocyte heterotopic transplant would be illuminating. That is, would a thalamic-born astrocyte be able to acquire the transcriptional and morphological profile of a cortical astrocyte if transplanted in early postnatal life? Evidence from such an experiment in septal astrocyte populations suggests the answer is yes^31^. Alternatively, but not mutually exclusively, early developmental regional patterning may “prime” astrocytes to receive and react appropriately to the signals they receive in their local niches later in development, as a recent study has shown in the context of GABA-induced morphogenesis^102^. We anticipate the current study will be a useful starting point for hypotheses such as these.

## Data and Code Availability

Raw data (sequencing reads in fastq format) for all 10x Chromium snRNAseq samples and CellBender-cleaned aligned counts matrices (in .h5 format) are publicly available for download on the Neuroscience Multi-omic Data Archive (NeMO) at: https://data.nemoarchive.org/biccn/grant/u01_feng/feng/transcriptome/sncell/10x_v3.1/. Pre-processed, clustered, and annotated data (in .h5ad format) is available for download, exploration, and gene search on the Broad Single Cell Portal at: https://singlecell.broadinstitute.org/single_cell/study/SCP2719/a-multi-region-transcriptomic-atlas-of-developmental-cell-type-diversity-in-mouse-brain (mouse) and https://singlecell.broadinstitute.org/single_cell/study/SCP2706/a-multi-region-transcriptomic-atlas-of-developmental-cell-type-diversity-in-marmoset-brain#study-summary (marmoset). Registered RNAscope HiPlex/v2 FISH image stacks, raw and background-subtracted expansion microscopy image volumes and binarized 3D tracings (both in .tif format) are available for download on BossDB at https://bossdb.org/project/schroeder2025. Custom scripts used to analyze data are available at https://github.com/Feng-Lab-MIT/AstrocyteHeterogeneity.

## Supporting information

Schroeder_2025_SI

Schroeder_2025_SITables

Schroeder_2025_SIMovie_Links

## Author Contributions

M.E.S., K.X.L, G.F., and F.M.K. conceptualized the project. M.E.S. harvested and microdissected mouse brain tissue and microdissected marmoset brain tissue with assistance from K.X.L. and D.M.M. Q.Z. oversaw marmoset breeding and donor selection and performed brain tissue harvest. M.E.S., L.M., and K.X.L. conducted single-nucleus RNA sequencing experiments. M.E.S., D.M.M., and K.X.L. analyzed the single-nucleus RNA sequencing data with guidance from F.M.K. M.E.S. and L.M. performed mouse colony management. L.M. performed stereotaxic surgery for AAV injection in adult mice and performed transcardial perfusions alongside D.M.M. E.Y. and J.K. optimized collagenase treatment for blood vessel preservation and made ExR gels. M.E.S. dissected, pre-stained, stained and imaged low-expansion ExR gels and analyzed images. J.K. stained and imaged high-expansion ExR gels. H.Z. and K.M.L. imaged marmoset tissue on the TissueFAXS microscope and provided guidance with marmoset tissue sectioning and staining. F.M.K., G.F., and E.S.B. supervised the project. M.E.S., F.M.K., E.S.B., and G.F. acquired funding. M.E.S. wrote and revised the manuscript with D.M.M., F.M.K., G.F., and E.S.B. An author contributions matrix, inspired by a recent *Nature* news article (https://www.nature.com/nature-index/news/researchers-embracing-visual-tools-contribution-matrix-give-fair-credit-authors-scientific-papers) is included below.

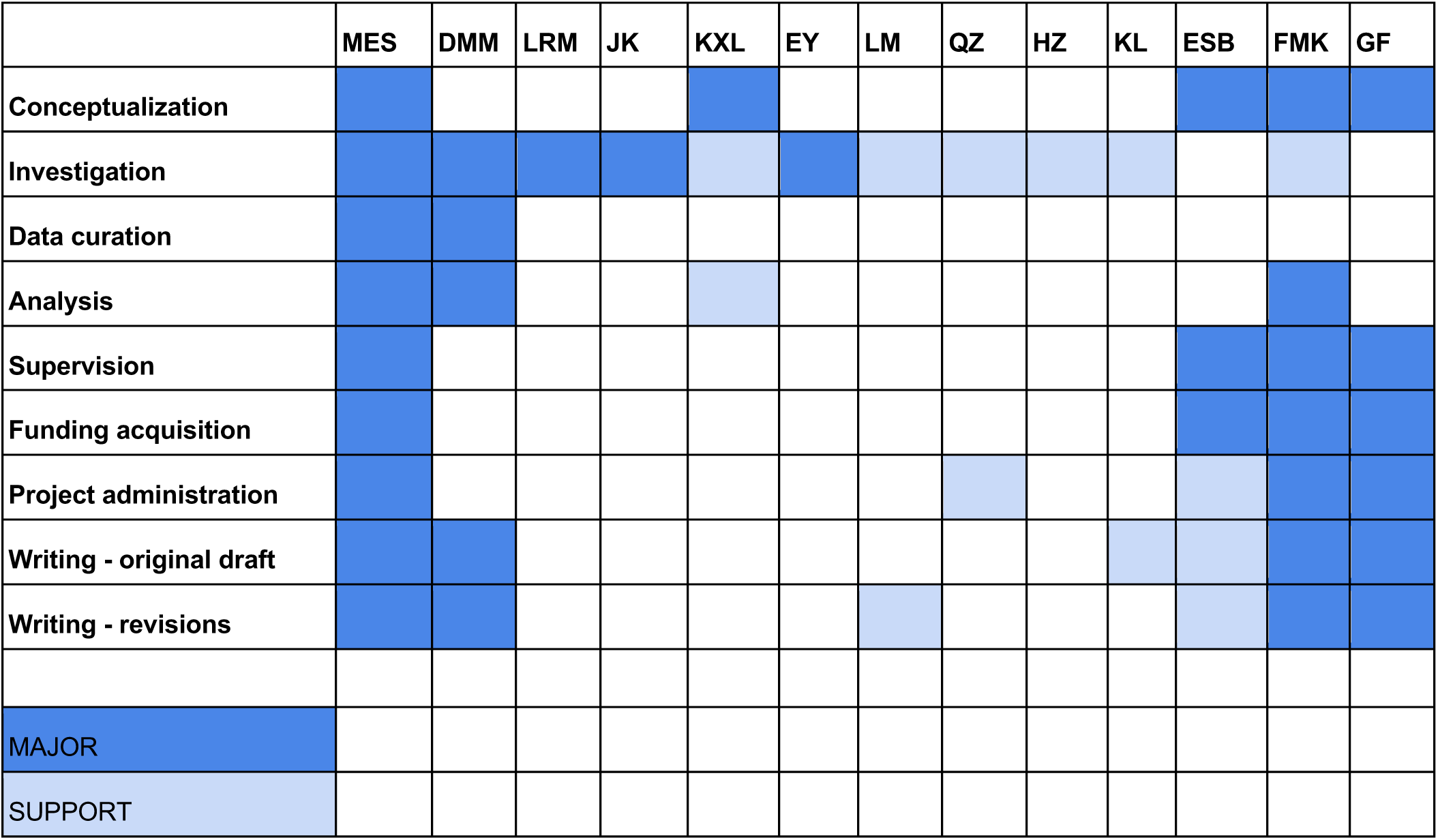

## Acknowledgements

Pictograms were generated using BioRender.com. MES was supported by the MathWorks Science Fellowship at MIT, the Collamore-Rogers Fellowship at MIT, the NSF Graduate Research Fellowship Program #1745302, and NIH 1F31MH133329-01. GF was supported by the National Institute of Mental Health and BRAIN Initiative (U01MH114819), Hock E. Tan and K. Lisa Yang Center for Autism Research of the Yang Tan Collective at MIT, the Poitras Center for Psychiatric Disorders Research at MIT, Stanley Center for Psychiatric Research at Broad Institute of MIT and Harvard. This work was supported by Brain Initiative grant UM1MH130981 to GF and FMK. ESB was supported by HHMI, Lisa Yang, NIH R01AG087374, NIH 1R01EB024261, Good Ventures, Tom Stocky and Avni Shah, Kathleen Octavio, NIH 1R01AG070831, European Research Council (ERC) SYNERGY Grant No 835102. We thank Eric Nyase, Andrew Harrahill, and In-Hye Kang for AAV packaging and preparation, Ken Chan and Ben Deverman for the BI103 plasmid and tissue from an injected animal co-stained with Sox9 to help assess expression in astrocytes, Michael Debarbardine for generating mitochondrial gene annotations for mCalJa1.2.Pat.X, Yanay Rosen for generating the marmoset protein embedding for SATURN, Menglong Zeng for guidance particularly on molecular cloning and for sharing many reagents, Morgan Fleishman for animal colony maintenance and support, Victoria Beja-Glasser for guidance and the gift of several wild-type mice, Ryan Kast for guidance particularly on RNA FISH and sharing many reagents, Nik Jorstad for guidance on analysis, Samia Silva de Castro, Jitendra Sharma, and Yefei Chen for performing necropsies and harvesting marmoset tissue, Alyssa Lutservitz and Xian Adikonis for tips on the 10x snRNAseq protocol, 10x Genomics Technical Support for answering many questions and providing replacement reagents when required, Jennifer Shih and Mriganka Sur for the gift of Aldh1l1-Cre mice, Lizzy Joo for the Cre genotyping primer sequences (originally from the Brueckner Lab at Yale), and Stephen Yu for assistance with marmoset PCR-based sex genotyping. We thank all members of the Feng and Boyden labs for insight and suggestions throughout the project. We gratefully acknowledge the MIT BioMicroCenter, the Broad Institute Genomics Platform, and MIT Department of Comparative Medicine for their support and assistance of this work.

## Supplementary Figures and Movies

**Supplementary Figure 1. Summary of sequencing coverage for the cross-region developmental snRNAseq atlas.**

**Supplementary Figure 2. Distribution of nuclei across cell types, regions, ages, sexes, and study for marmoset single-nucleus RNAseq data.**

**Supplementary Figure 3. Distribution of nuclei across cell types, regions, ages, sexes, and study for mouse single-nucleus RNAseq data.**

**Supplementary Figure 4. Leiden cluster annotations and proportions for the cross-region developmental snRNAseq atlas.**

**Supplementary Figure 5. Dissection strategies for marmoset brains and region reassignment to mitigate cross-region contamination for developing samples**

**Supplementary Figure 6. Region reassignment for cross-contaminant nuclei in mouse samples.**

**Supplementary Figure 7. Correlation of pairwise astrocyte rDEG log-fold change between region pairs across development in mouse and marmoset.**

**Supplementary Figure 8. Validation of marmoset astrocyte rDEG expression in situ using multiplexed FISH.**

**Supplementary Figure 9. Quantification of selected rDEG and astrocyte subtype marker expression in situ in adult and neonate marmoset.**

**Supplementary Figure 10. Validation of mouse astrocyte rDEG expression in situ using multiplexed FISH.**

**Supplementary Figure 11. Quantification of selected rDEG and astrocyte subtype marker expression in situ in adult and neonate mouse.**

**Supplementary Figure 12. Astrocyte sub-clustering captures intra-regional heterogeneity in marmoset.**

**Supplementary Figure 13. Astrocyte sub-clustering captures intra-regional heterogeneity in mouse.**

**Supplementary Figure 14. Cell-cell communication analysis for neuron-astrocyte and astrocyte-neuron predicted ligand-receptor pairs across regions and developmental time points in mouse.**

**Supplementary Figure 15. Pseudotime inference in mouse and marmoset oligodendrocyte lineage and mouse astrocytes.**

**Supplementary Figure 16. Gene expression signatures underlying the postnatal developmental specification of mouse astrocytes within and across brain regions.**

**Supplementary Figure 17. Cell type composition across development in both species and cross-species integration with SATURN.**

**Supplementary Figure 18. Expression of Top2a in astrocytes of the mouse subventricular zone via multiplexed FISH.**

**Supplementary Figure 19. Additional detail on astrocyte morphology and rDEG protein expression differences across brain regions in mouse.**

**Supplementary Figure 20. High-concentration collagenase treatment preserves blood vessel morphology in ExR samples.**

**Supplementary Movies 1-36. 3D-projected image volumes of ∼4x expanded mouse astrocytes.**

## Supplementary Tables

**Supplementary Table 1. Biological donor information.**

**Supplementary Table 2. Abundance of cell types and Leiden clusters and descriptions of Leiden clusters in mouse and marmoset.**

**Supplementary Table 3. Proportional breakdown of each Leiden cluster by age, assigned region, and sex for mouse and marmoset.**

**Supplementary Table 4. Proportional breakdown of MapMyCells-derived Allen Brain Cell Atlas subclass assignments by Leiden cluster for mouse and marmoset.**

**Supplementary Table 5. Summary of significant compositional differences in cell type for marmoset as found by scCODA.**

**Supplementary Table 6. Summary of significant compositional differences in cell type for mouse as found by scCODA.**

**Supplementary Table 7. Summary of significant compositional differences in Leiden cluster for marmoset as found by scCODA.**

**Supplementary Table 8. Summary of significant compositional differences in Leiden cluster for mouse as found by scCODA.**

**Supplementary Table 9. Marmoset rDEGs shared across both replicates for each age.**

**Supplementary Table 10. WebGestalt enrichment results for marmoset rDEGs by age.**

**Supplementary Table 11. Mouse rDEGs for each age.**

**Supplementary Table 12. WebGestalt enrichment results for mouse rDEGs by age.**

**Supplementary Table 13. Image-level quantification of each rDEG probe for adult and neonate marmoset and mouse and results of statistical tests for mouse.**

**Supplementary Table 14. Proportion astrocytes positive for one or more fibrous marker(s) in each time point-region combination for both species.**

**Supplementary Table 15. Top ABCA MapMyCells cluster assignments for astrocyte subclusters.**

**Supplementary Table 16. CellPhoneDB-generated cell-cell communication results for each region-age combination in mouse and marmoset.**

**Supplementary Table 17. Fraction of CellPhoneDB-generated ligand-receptor pairs shared across 3 or more ages for all neuronal clusters within a region for both species.**

**Supplementary Table 18. Genes with the highest Mellon change scores over pseudotime for mouse and marmoset telencephalic and diencephalic astrocytes.**

**Supplementary Table 19. Marmoset astrocyte, OPC, GABAergic, and glutamatergic neuron aDEGs for each brain region.**

**Supplementary Table 20. Mouse astrocyte, OPC, GABAergic, and glutamatergic neuron aDEGs for each brain region.**

**Supplementary Table 21. Shared and species-unique cortex-thalamus astrocyte rDEGs for each developmental timepoint in mouse and marmoset.**

**Supplementary Table 22. Shared and species-unique fetal-late adolescent astrocyte aDEGs for each region in mouse and marmoset.**

**Supplementary Table 23. sDEGs for each supercluster.**

**Supplementary Table 24. Telencephalic-specific, diencephalic-specific, and shared astrocyte sDEGs.**

**Supplementary Table 25. Summary statistics for quantification of mouse astrocyte morphology across brain regions.**

**Supplementary Table 26. Summary statistics for quantification of Glast and Gat3 expression in ∼18x ExR images from mouse PFC, striatum, and thalamus.**

**Supplementary Table 27. Blocking strategy for marmoset region dissection based on custom brain matrix.**

**Supplementary Table 289. Sequencing coverage statistics for all 10x Chromium reactions.**

**Supplementary Table 30. Table used to convert between mouse, marmoset, and human gene IDs for 1:1 orthologs.**

**Supplementary Table 31. List of reagents, including vendor and product information, used for experiments.**

**Supplementary Table 32. Expansion factor measurements for mouse ExR samples.**

**Supplementary Table 33. Notes on each ∼4x expanded mouse astrocyte imaged (Fig. 7 and Fig. S19).**

