## Supplementary material for "Astrocyte regional specialization is shaped by postnatal development": Schroeder_2025_SI

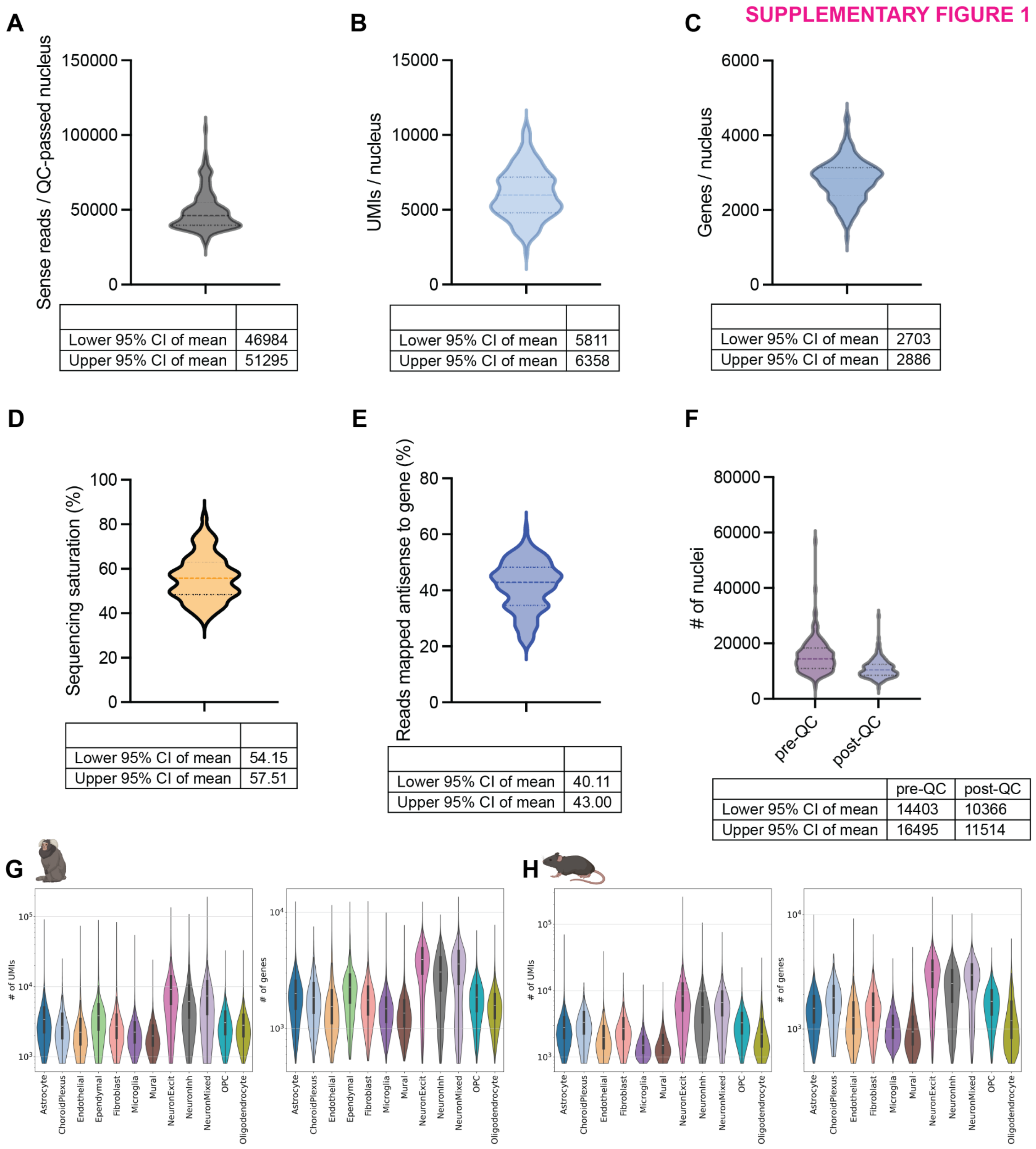


**Supplementary Figure 1. Summary of sequencing coverage for the cross-region developmental snRNAseq atlas. A,** Violin plot showing the estimated population distribution of sense reads per rough quality control (QC)-passed (low-UMI and low-gene but not doublet removal, see **Methods**) for all nuclei across all 144 10x Chromium reactions sequenced in the current study (not including adult marmoset data from our previous study[7]). Dashed lines indicate the median and quartiles. **B**, Violin plot (as described for **(A)**) of the median number of unique molecular identifiers (UMIs) per nucleus as described in **(A)** (n = 144 10x Chromium reactions). **C,** Same as **(B)**, for the median number of genes per nucleus. **D**, Same as **(C)**, for sequencing saturation as calculated by 10x Cell Ranger software (v7.1). **E**, Same as **(D)**, for the fraction of reads mapped antisense to genes, as calculated by 10x Cell Ranger software (v7.1). **F**, Number of nuclei called by 10x Cell Ranger software before (pre-QC) and after (post-QC) rough QC (low-UMI and low-gene but not doublet removal, see **Methods**). **G,** Violin plot (scanpy's default) showing the estimated distribution of the number of UMIs (left) and genes (right) for full QC-passed marmoset nuclei, as presented in the main figures, grouped by cell type. **H**, Same as **(G)**, for mouse nuclei.

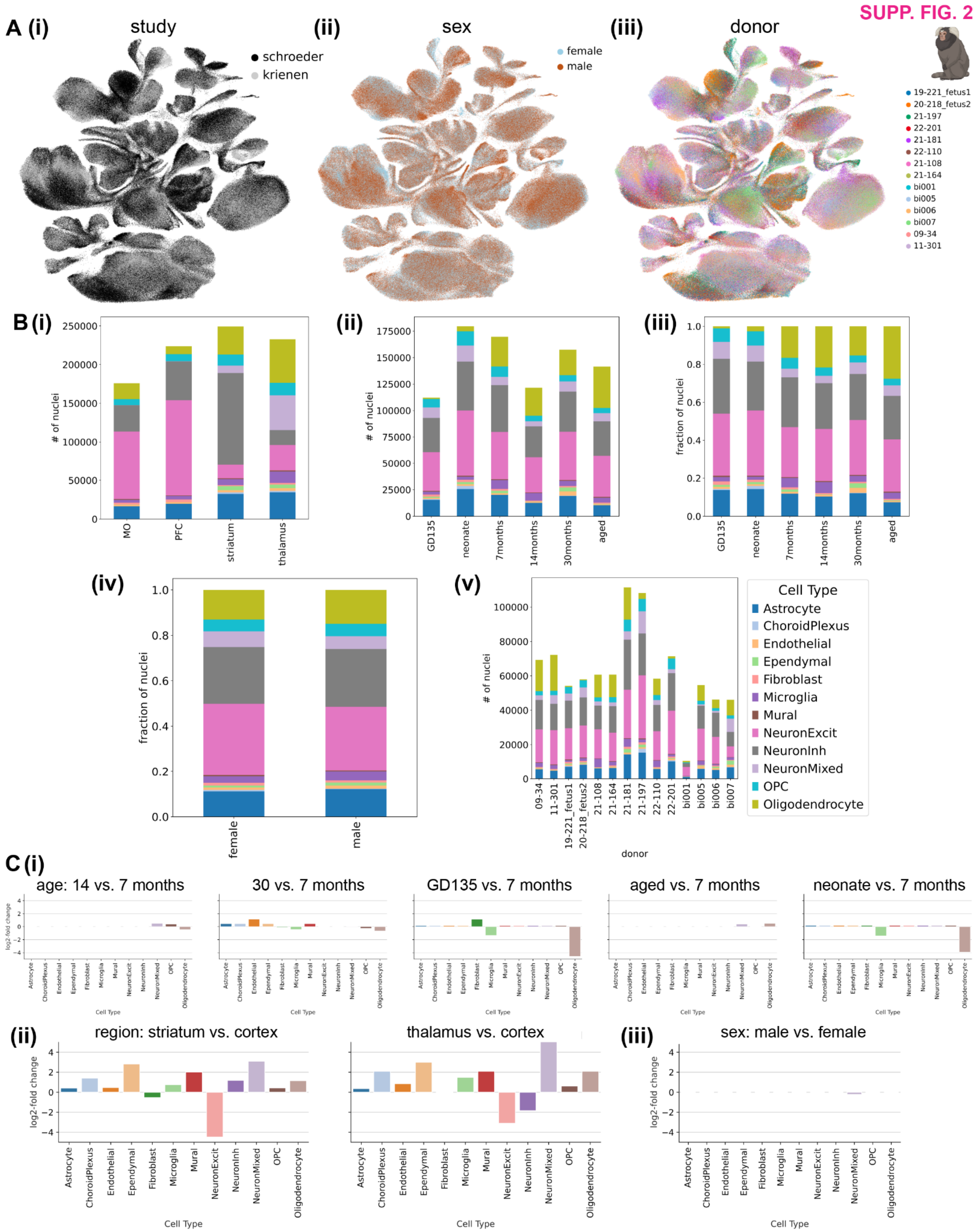


**Supplementary Figure 2. Distribution of nuclei across cell types, dissected regions, ages, sexes, and study for marmoset single-nucleus RNAseq data. A,** Integrated UMAP embedding of 881,832 marmoset nuclei from PFC, motor cortex, striatum, and thalamus (as dissected, before reassignment) across all developmental time points assayed and a randomly downsampled portion of adult nuclei from our previous study[7] colored by **(i)** study (present, “schroeder” or previous, “krienen”), (**ii)** sex, and **(iii)** biological replicate (for marmoset equivalent to single donor). **B**, Stacked bar plots showing the number or proportion of full QC-passed nuclei for each **(i)** dissected brain region, **(ii)** age (number of nuclei), **(iii)** age (proportion of nuclei) **(iv)** sex (proportion of nuclei) and **(v)** biological replicate, broken down by assigned cell type. **C**, Log2-fold change in cell type composition as estimated by single-cell compositional data analysis (scCODA)[42] for the coefficients on **(i)** age (reference = 14 months), **(ii)** region (reference = cortex), and **(iii)** male sex (reference = female).Only fold-changes for credible effects (i.e., significantly nonzero coefficient) are shown. The 881,832 marmoset nuclei were composed of 12 broad cell classes: glutamatergic excitatory neurons (29.7%, GABAergic inhibitory neurons (25.2%), neurons belonging to mixed glutamatergic and GABAergic clusters (6.24%), astrocytes (11.7%), oligodendrocytes (14.0%), oligodendrocyte progenitor cells (OPCs, 5.35%), microglia and macrophages (3.45%), mural cells (0.51%), endothelial cells (1.22%), vascular-associated fibroblasts (1.05%), ependymal cells (1.11% in), and choroid plexus cells (0.45%).

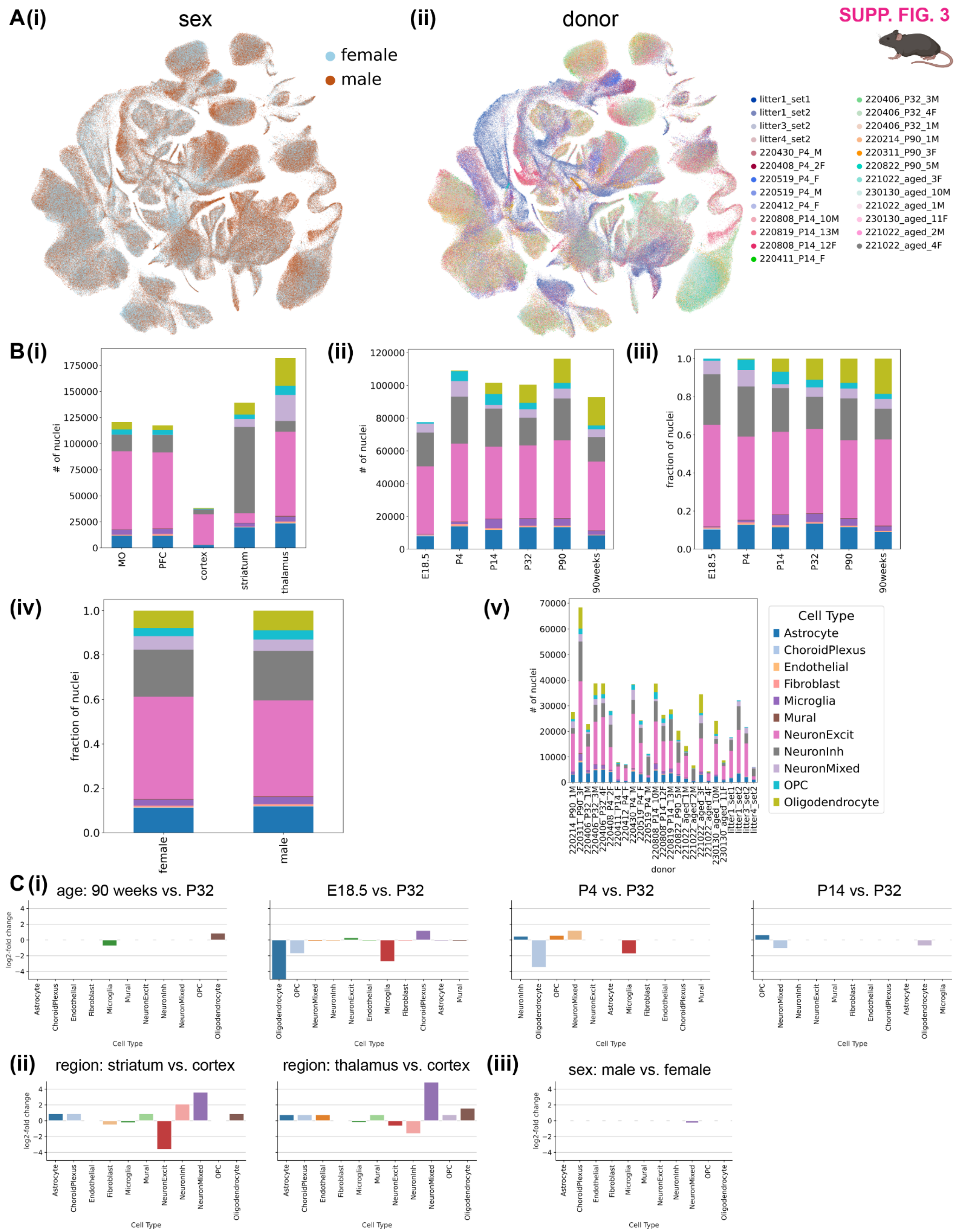


**Supplementary Figure 3. Distribution of nuclei across cell types, regions, ages, sexes, and study for mouse single-nucleus RNAseq data. A,** Integrated UMAP embedding of mouse (597,668 nuclei) from PFC, motor cortex, striatum, and thalamus (as dissected, before reassignment) across all developmental time points assayed and a randomly downsampled portion of adult nuclei from our previous study[7] colored by **(i)** sex and **(ii)** biological replicate (for most mouse time points, is the same as individual donor). **B**, Stacked bar plots showing the number or proportion of full QC-passed nuclei for each **(i)** dissected brain region, **(ii)** age (number of nuclei), **(iii)** age (proportion of nuclei) **(iv)** sex (proportion of nuclei) and **(v)** biological replicate, broken down by assigned cell type. **C**, Log2-fold change in cell type composition as estimated by scCODA for the coefficients on **(i)** age (reference = P32), **(ii)** region (reference = cortex), and **(iii)** male sex (reference = female) for. Only fold-changes for credible effects (i.e., significantly nonzero coefficient) are shown. The 597,668 mouse nuclei were composed of 12 broad cell classes: glutamatergic excitatory neurons (44.7%), GABAergic inhibitory neurons (21.8%), neurons belonging to mixed glutamatergic and GABAergic clusters (5.54%), astrocytes (11.5%), oligodendrocytes (8.41%), oligodendrocyte progenitor cells (OPCs, 3.91%), microglia and macrophages (2.98%), mural cells (0.28%), endothelial cells (0.26%), vascular-associated fibroblasts (0.62%), ependymal cells (0%; these instead co-clustered with astrocytes), and choroid plexus cells (0.12%).

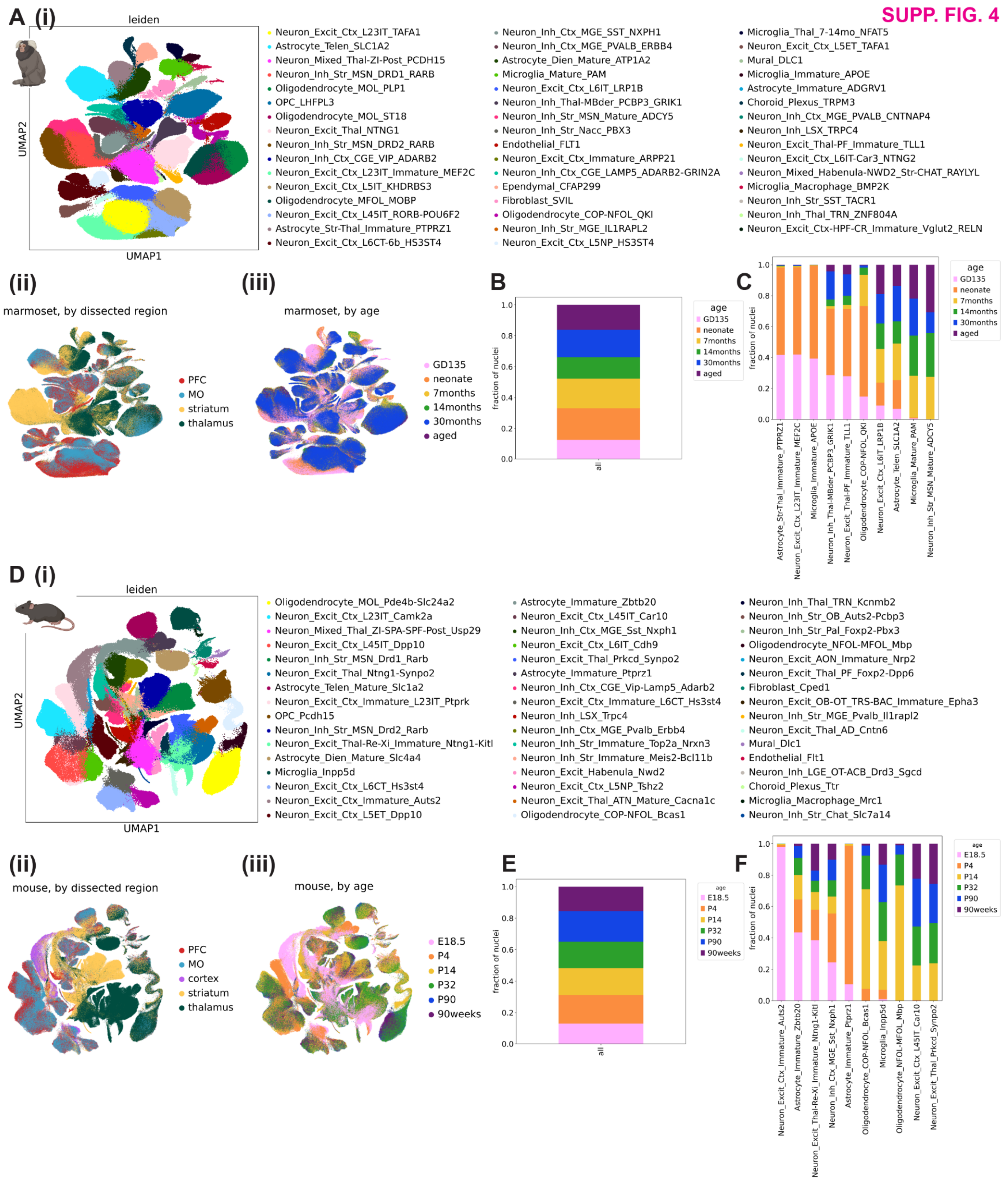


**Supplementary Figure 4. Leiden cluster annotations and proportions for the cross-region developmental snRNAseq atlas. A,** Integrated UMAP embedding of marmoset (**A**, 881,832) or mouse (**B**, 597,668) nuclei from PFC, motor cortex, striatum, and thalamus (as dissected, before reassignment) across all developmental time points assayed and a randomly downsampled portion of adult nuclei from our previous study[7] colored by **(i)** Leiden-algorithm determined cluster annotation (hereafter, Leiden cluster), **(ii)** dissected region and **(iii)** developmental time point. **B**, Stacked bar plot showing the proportion of all marmoset nuclei from each developmental time point (see legend in **A(iii)**). **C,** Stacked bar plot showing the proportion of each Leiden cluster from each developmental time point for a hand-selected set of 10 clusters that are developmentally enriched or depleted. **D,** Stacked barplots showing the number of nuclei assigned to each of the top 5 most abundant Allen Brain Cell Atlas mouse whole-brain atlas class or subclass using MapMyCells[41] for **(i)** excitatory neurons **(ii)** inhibitory neurons and **(iii)** astrocytes, broken down by dissected brain region. **E-H,** Same as **A-D,** for mouse.

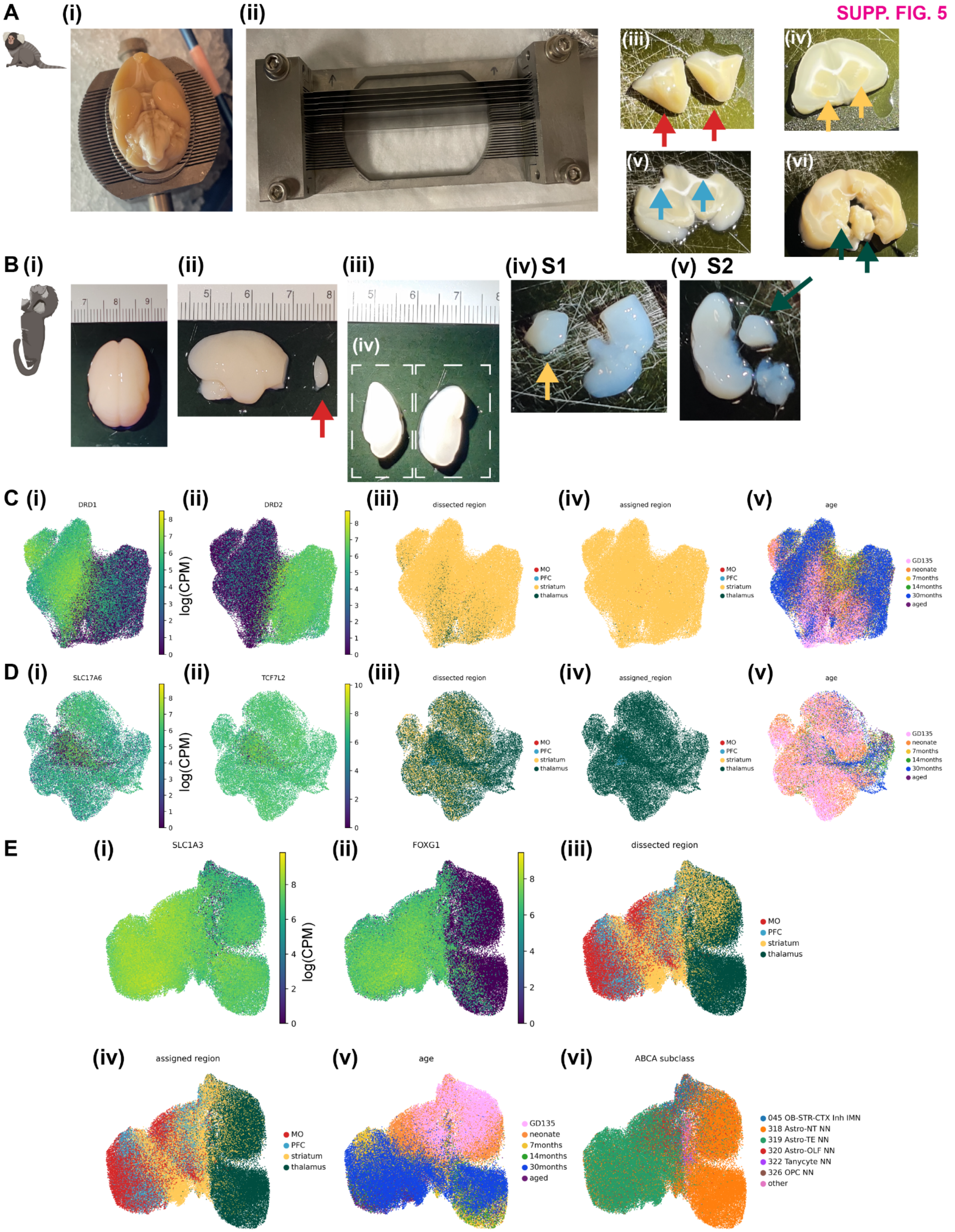


**Supplementary Figure 5. Dissection strategies for marmoset brains and region reassignment to mitigate cross-region contamination for developing samples. A**, Dissection strategy for 7-month, 14-month, and aged marmosets used in this study. **(i)** Ventral view of brain from female 14-month marmoset donor 21-164 in brain matrix. **(ii)** Blades positioned in blade holder to create desired tissue slabs for dissection. Arrows on the blade holder indicate anterior direction. **(iii)** Tissue slab for prefrontal cortex (PFC) dissection, with red arrows indicating PFC. Some white matter was removed, and then the entire slab was frozen for nuclei isolation. **(iv)** One out of two tissue slabs used for striatum dissection, with yellow arrows indicating striatum (not yet dissected). **(v)** Tissue slab used for motor cortex dissection, with blue arrows indicating where motor cortex was removed (already removed in this photo). **(vi)** Tissue slab used for thalamus dissection, with dark green arrows indicating the thalamus (already dissected from the left hemisphere, but not yet removed). **B,** Dissection strategy for GD135 marmoset (similar for neonate). **(i)** Dorsal view of brain from female GD135 donor 19-221_fetus1. **(ii)** Prefrontal cortex as grossly dissected from lateral surface of the brain. **(iii)** Two slabs cut from the bissected region spanning approximately from the anterior beginning of the temporal lobe to its posterior end, used to dissect the striatum (slab 1, S1) and thalamus (slab 2, S2). **(iv)** Striatum as dissected from slab 1 (S1). **(v)** Thalamus as dissected from slab 2 (S2). **C**, Analysis of dissection cross-contamination in marmoset (all ages, but only nuclei from GD135 and neonate were reassigned) using medium spiny neurons (MSNs). **(i-v)** UMAP projection of marmoset MSNs colored by **(i)** *DRD1* expression in logCPM units, **(ii)** *DRD2* expression in logCPM units, **(iii)** dissected brain region, **(iv)** reassigned brain region, and **(v)** age. **(iii)** Shows contamination of thalamus in striatal dissections for GD135 and neonate samples. **D,** Same as **(C)**, for marmoset thalamic excitatory neurons, with **(i)** showing *SLC17A6* expression and **(ii)** showing *TCF7L2* expression. **(iii)** Shows contamination of striatum in thalamic dissections for GD135 and neonate samples. **E,** Analysis of dissection cross-contamination in marmoset (all ages, but only nuclei from GD135 and neonate were reassigned) astrocytes of all clusters. UMAP projection of marmoset astrocytes colored by **(i)** *SLC1A3* expression, **(ii)**, *FOXG1* expression, **(iii)** dissected brain region, **(iv)** reassigned brain region, **(v)** age, and **(vi)** ABCA subclass, as determined using MapMyCells.

SUPP. FIG. 6

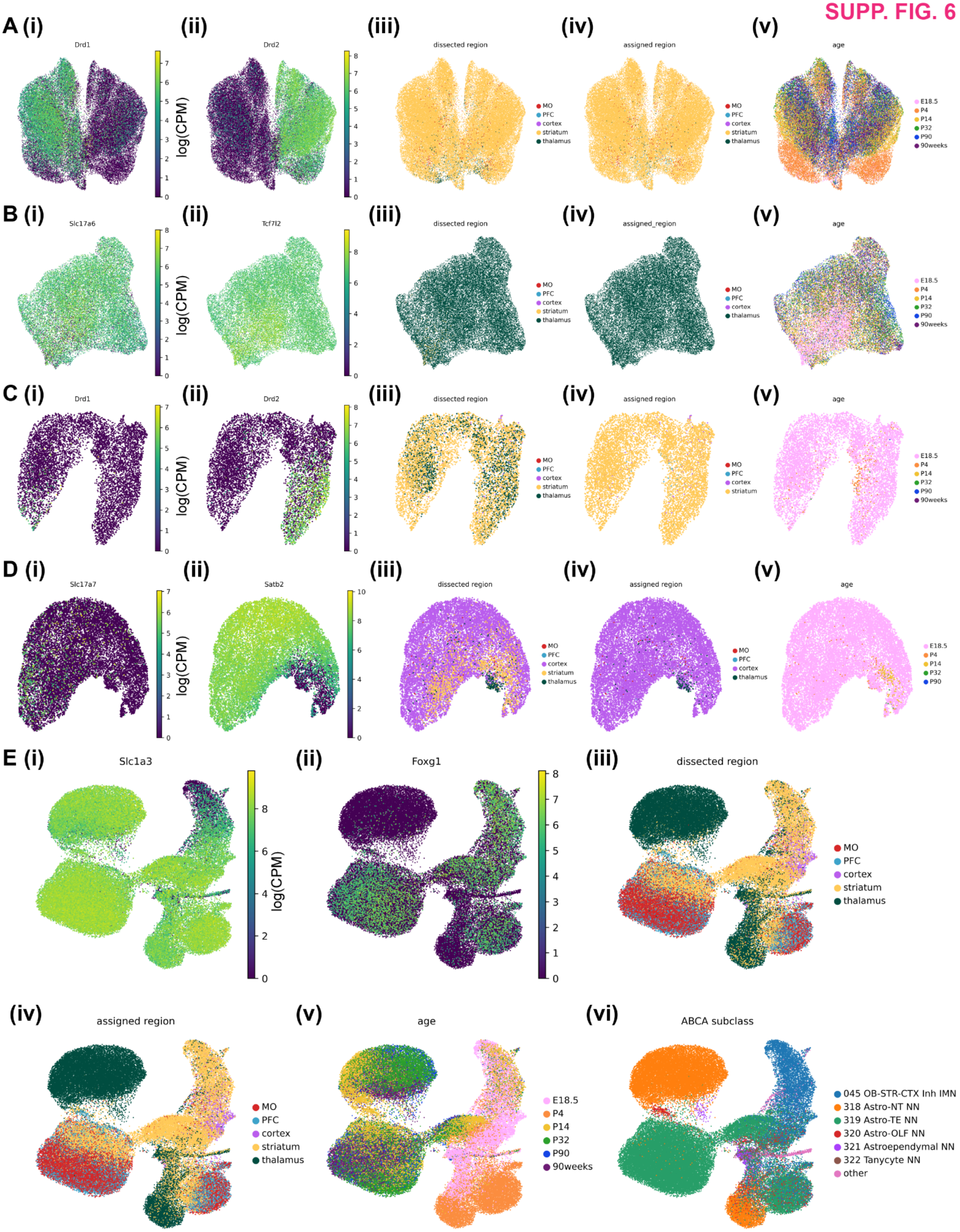


**Supplementary Figure 6. Region reassignment for cross-contaminant nuclei in mouse samples. A,** Analysis of dissection cross-contamination in mouse (all ages) using medium spiny neurons (MSNs). **(i-v)** UMAP projection of mouse MSNs colored by **(i)** *Drd1* expression in logCPM units, **(ii)** *Drd2* expression in logCPM units, **(iii)** dissected brain region, **(iv)** reassigned brain region, and **(v)** age. **B,** Same as **(A)**, for mouse thalamic excitatory neurons, with **(i)** showing *Slc17a6* expression and **(ii)** showing *Tcf7l2* expression. **C,** Same as **(A),** for immature mouse MSNs, showing a higher degree of striatum-thalamus cross-contamination in E18.5 mouse samples. **D,** Same as **(A-C)**, for immature mouse cortical neurons, with **(i)** showing *Slc17a7* expression and **(ii)** showing *Satb2* expression. **(iii)** Illustrates cortical contamination of striatal and thalamic dissections in the mouse E18.5 data. **E,** Analysis of dissection cross-contamination in mouse (all ages) astrocytes of all clusters. UMAP projection of mouse astrocytes colored by **(i)** *Slc1a3* expression, **(ii)**, *Foxg1* expression, **(iii)** dissected brain region, **(iv)** reassigned brain region, **(v)** age, and **(vi)** ABCA subclass, as determined using MapMyCells.

**SUPP. FIG. 7**

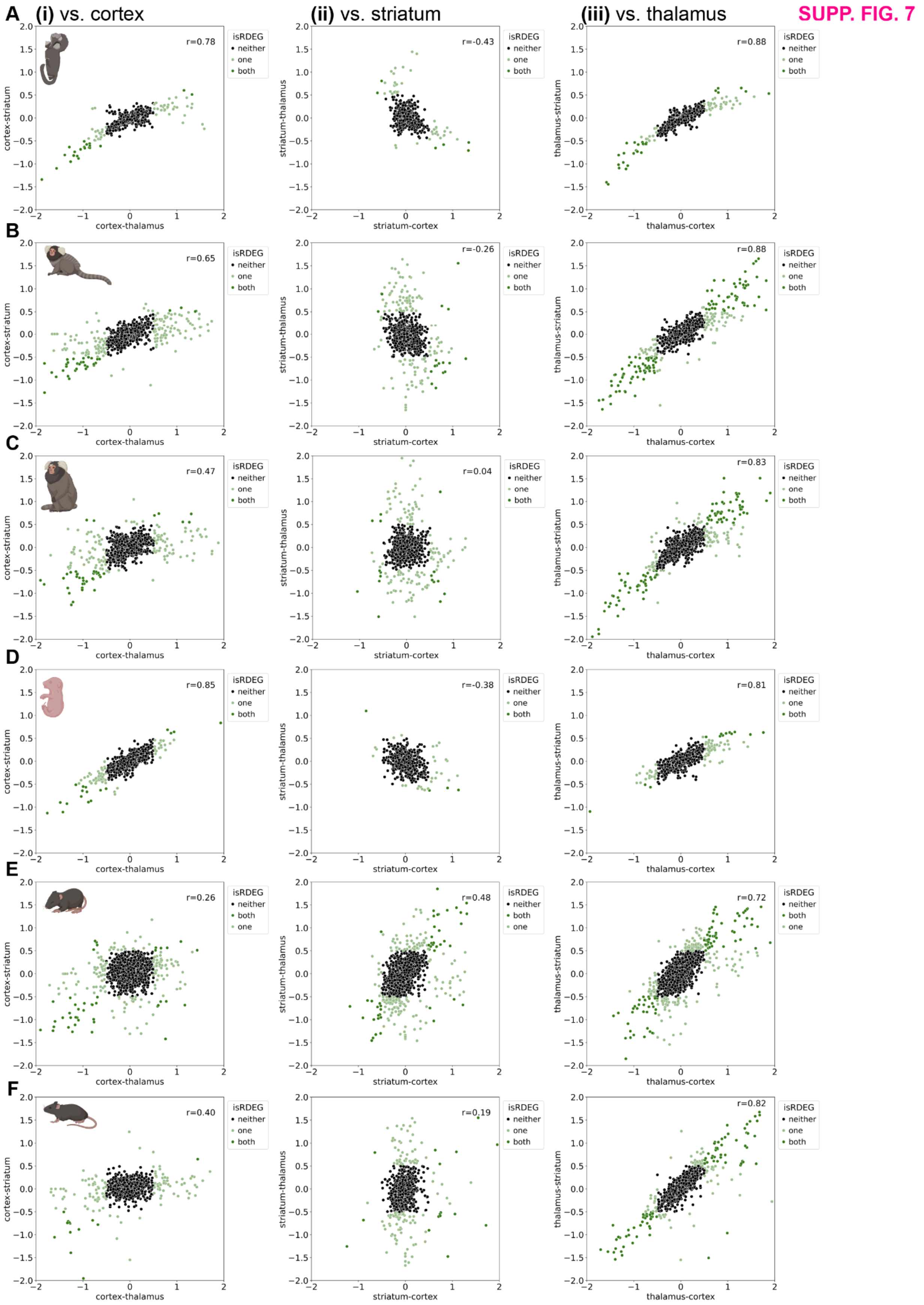


**Supplementary Figure 7. Correlation of pairwise astrocyte rDEG log-fold change between region pairs across development in mouse and marmoset. A**, Scatterplot of log fold-change (logFC) difference in expression in marmoset GD135 astrocytes for genes meeting minimum metacell expression criteria (relaxed from the criteria for rDEGs shown in **Figs. 2-3**, see **Methods**) for 2 different region pairs: **(i)** cortex-striatum vs. cortex-thalamus (a negative value indicates downregulation in cortex), **(ii)** striatum-thalamus vs. striatum-cortex (a negative value indicates downregulation in striatum), and **(iii)** thalamus-striatum vs. thalamus-cortex (a negative value indicates downregulation in thalamus). Light green dots indicate genes who meet rDEG (magnitude of logFC > 0.5) criteria for one region pair, and dark green dots indicate genes who meet rDEG criteria for both region pairs. In marmoset, logFC is averaged across replicates. r, Pearson's correlation coefficient calculated using all the points shown. **B,** Same as **(A)** for 7-month marmoset. **C,** Same as **(A)** for 30-month marmoset. **D**, Same as **(A)** for E18.5 mouse. **E**, Same as **(A)** for P14 mouse. **F**, Same as **(A)** for P90 mouse.

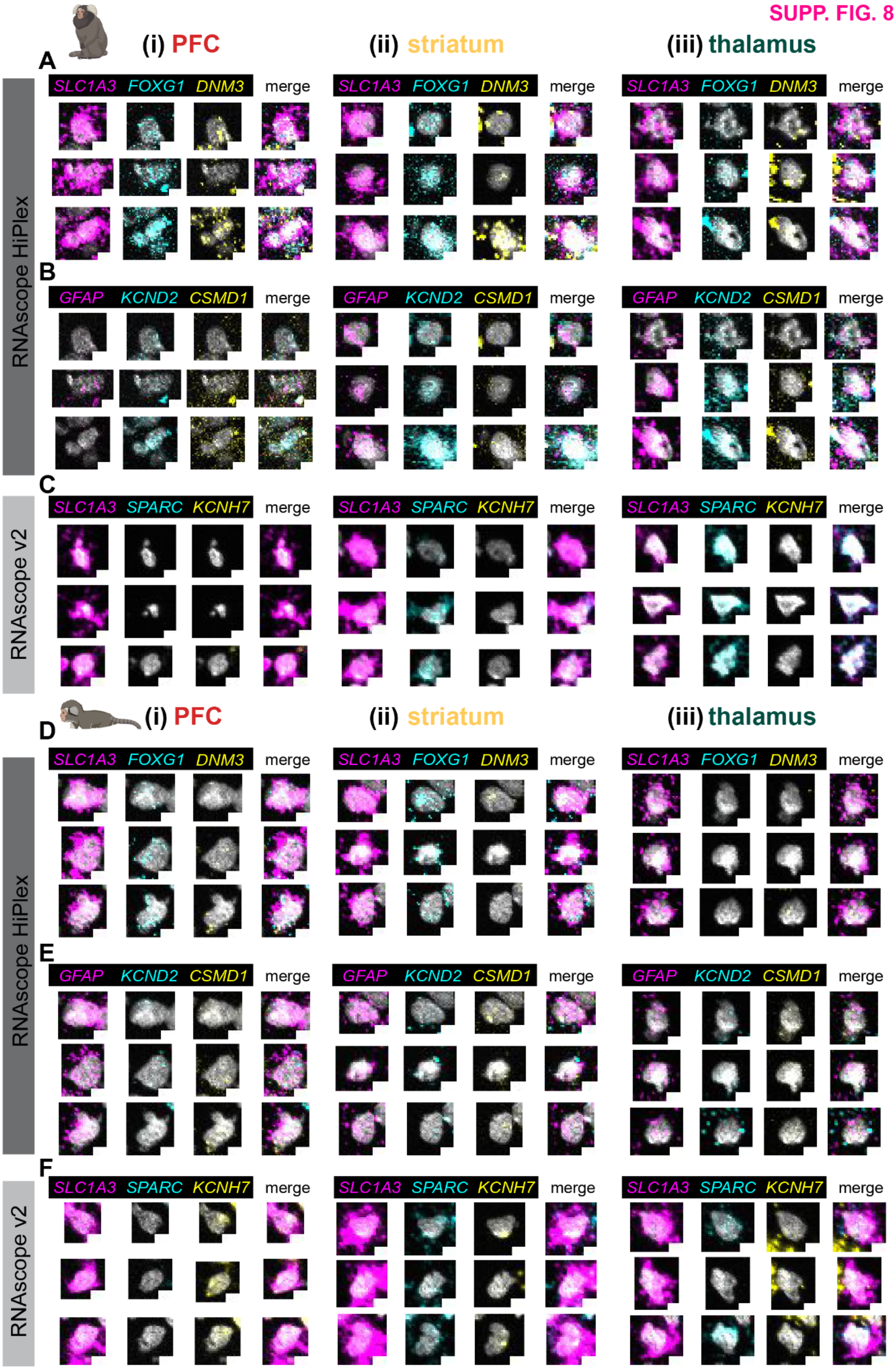
(i) PFC
(ii) striatum
(iii) thalamus
A
SLC1A3 FOXG1 DNM3 merge
B
GFAP KCND2 CSMD1 merge
C
SLC1A3 SPARC KCNH7 merge
RNAscope HiPlex
RNAscope v2
D
SLC1A3 FOXG1 DNM3 merge
E
GFAP KCND2 CSMD1 merge
F
SLC1A3 SPARC KCNH7 merge

**Supplementary Figure 8. Validation of marmoset astrocyte rDEG expression in situ using multiplexed and single-round FISH. A**, Single-channel and composite registered (across rounds) maximum-projected images of telencephalic astrocyte rDEGs for 3 exemplary astrocytes (low autofluorescence and high signal-to-noise) from one adult female marmoset in **(i)** PFC, **(ii)** striatum, and **(iii)** thalamus, generated using the RNAscope HiPlex protocol. *SLC1A3* was used to identify astrocytes. Throughout the images, DAPI nuclear counterstain is shown in gray. **B,** Same as **(A)**, for diencephalic astrocyte rDEGs. Scale bar, 5μm. **C,** Single-channel and composite images of rDEGs for 3 representative astrocytes from one adult female marmoset generated using the RNAscope v2 protocol. Scale bar, 5μm. **D-F,** Same as **A-C**, for one male **(D-E)** and one female **(F)** neonate marmoset. Contrast was manually adjusted by setting minimum and maximum intensity values, using the same values across regions within each age, except for *SLC1A3* in adult **(A)**, for which we reduced the maximum intensity value in the thalamus (vs. cortex and striatum), to account for lower expression in order to visualize its expression in astrocytes.

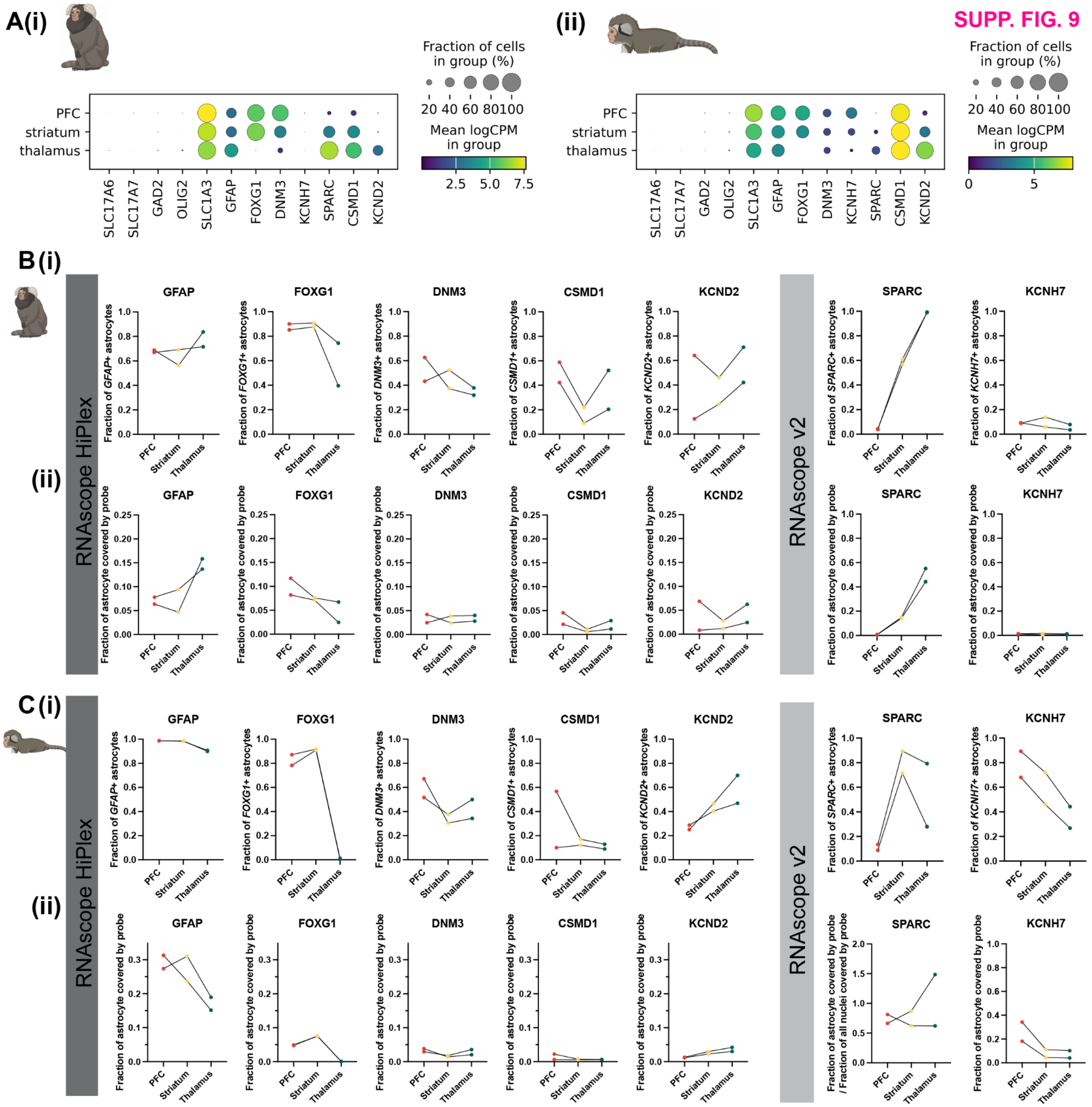


**Supplementary Figure 9. Quantification of selected rDEG and astrocyte subtype marker expression *in situ* in adult and neonate marmoset. A,** Mean expression (in logCPM from the snRNAseq data) of selected rDEG (*DNM3, KNC7, SPARC, CSMD1, KCND2*) and astrocyte subtype (*GFAP*) genes by region in the **(i)** adult (30 month) marmoset and **(ii)** neonate marmoset. **B, (i)** Line plots showing the fraction of astrocytes positive for these genes in PFC (red), striatum (yellow), and thalamus (green) from the RNAscope HiPlex or v2 FISH data (**Fig. S8**) for 2 adult female marmosets. **(ii)** Line plots showing the mean fraction of astrocytes covered by probes as in **(i)**. See **Methods** for details on quantification, which was performed semi-automatically using CellProfiler. Colored dots connected by lines are data originating from the same marmoset donor. **C,** Same as **(B)**, for two neonate marmoset donors, one male and one female.

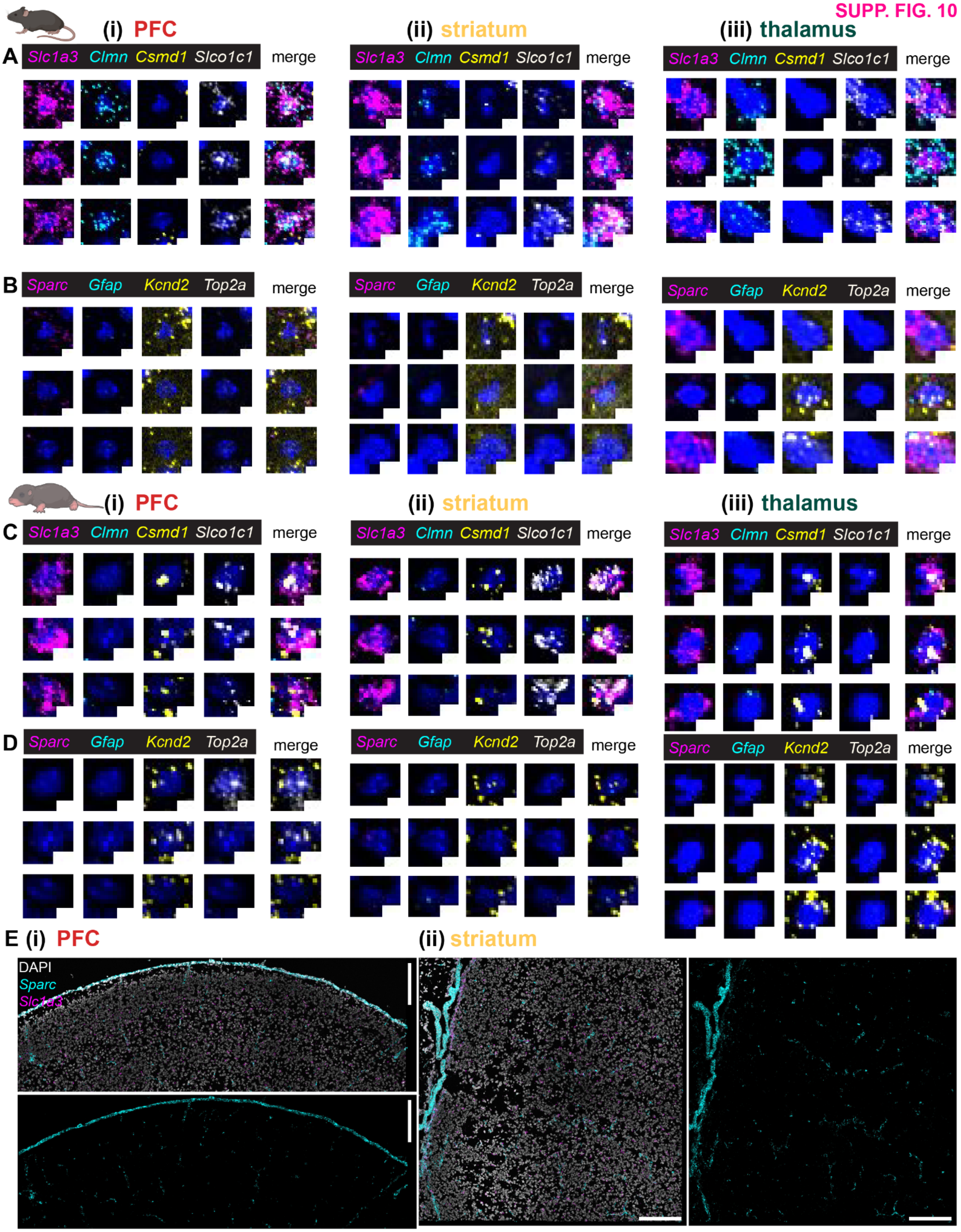


**Supplementary Figure 10. Validation of mouse astrocyte rDEG expression in situ using multiplexed FISH.**
**A,** Single-channel and composite registered (across rounds), maximum-projected images of telencephalic astrocyte

rDEGs for 3 exemplary astrocytes in the **(i)** PFC **(ii)** striatum and **(iii)** thalamus in one female P90 mouse, generated using RNAscope HiPlex FISH. **B,** Same as **(A)**, for diencephalic rDEGs (*Sparc, Kcnd2*) and other astrocyte subtype markers (*Gfap* and *Top2a*). Scale bar, 5μm. **C-D,** Same as **A-B**, for one male P4 mouse. Contrast was manually adjusted by setting minimum and maximum intensity values, using the same values across regions within each age. **E,** *Sparc* expression pattern in early postnatal mouse. Composite and single-channel registered, max-projected, stitched images of the entire field of view imaged for **(i)** PFC and **(ii)** striatum in one P4 female mouse, generated using RNAscope HiPlex FISH. *Sparc* staining is shown alone to aid visualization of its high signal intensity along brain borders (pial surface and ventricles) and along the vasculature (based on morphology) at this time point. *Sparc* contrast was increased to aid visualization in the larger, lower-zoom images. The PFC image has been rotated 90°C from its original orientation. Scale bar, 200μm.

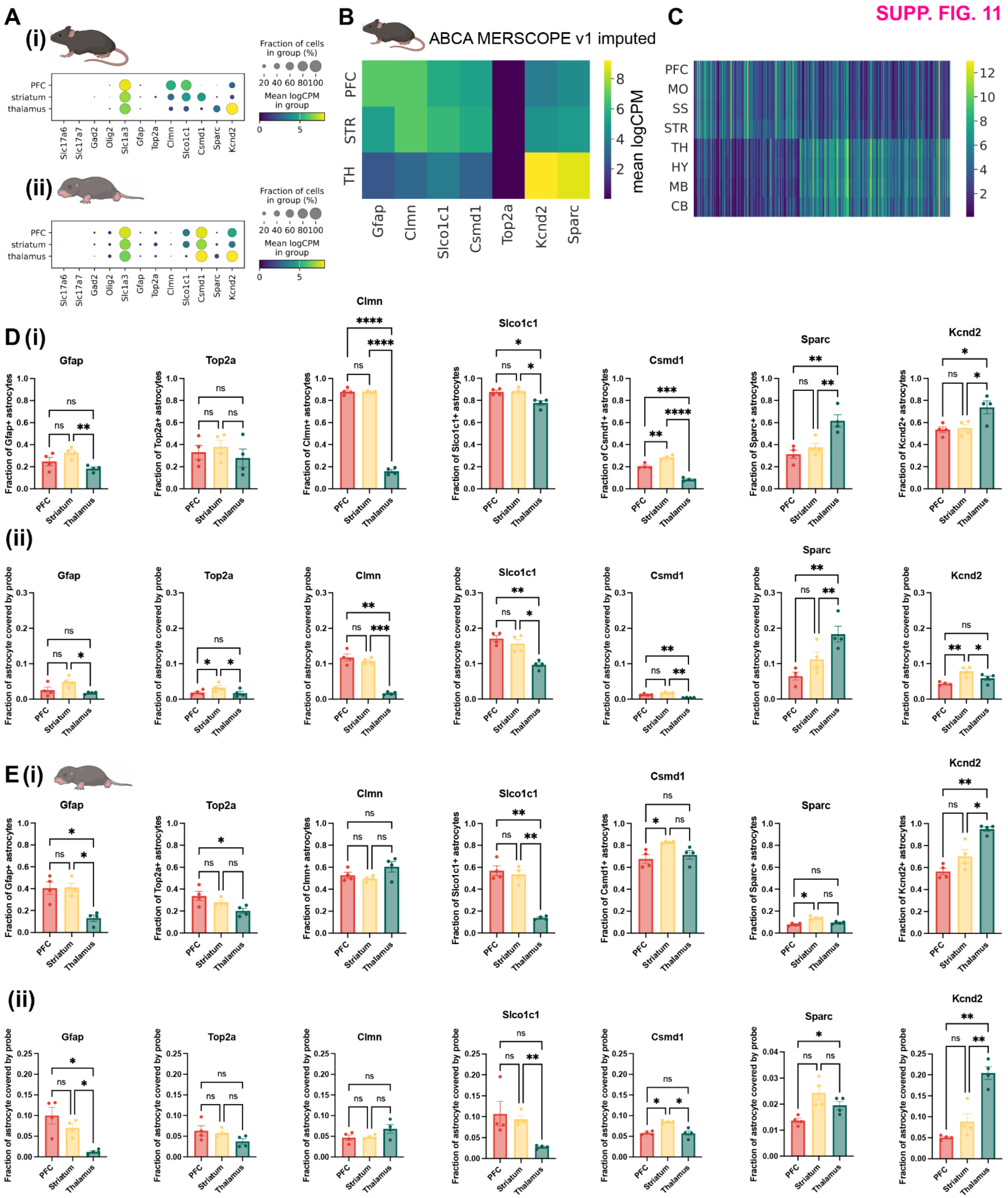


**Supplementary Figure 11. Quantification of selected rDEG and astrocyte subtype marker expression *in situ* in adult and neonate mouse. A,** Mean expression (in logCPM from the snRNAseq data) of selected rDEG (*Clmn, Slco1c1, Csmd1, Sparc,* and *Kcnd2*) and astrocyte subtype (*Gfap, Top2a*) genes by region in the **(i)** adult (P90) and **(ii)** neonate (P4) mouse. **B,** Mean expression (across astrocyte nuclei in each brain region, in logCPM) of the genes in **(A)** in astrocytes from the Allen Brain Cell Atlas (ABCA) MERSCOPE v1 whole mouse brain dataset with imputed genes from 10x Genomics scRNAseq (https://knowledge.brain-map.org/abcatlas).[1] PFC, prefrontal cortex. STR, striatum. TH, thalamus. **C,** Mean expression (across astrocyte nuclei in each brain region, in logCPM) of all mouse P90 astrocyte rDEGs from the ABCA MERSCOPE v1 whole mouse brain dataset as in **(B)**, with additional brain regions added. SS, somatosensory cortex. HY, hypothalamus. MB, midbrain. CB, cerebellum. **(i)** Bar plots showing the mean fraction of astrocytes positive for these genes in PFC (red), striatum (yellow), and thalamus (green) from FISH data in P90 mice (**Fig. S10**, n = 4 mice, 2 of each sex, data point shows average of 2 tissue sections each). **(ii)** Bar plots showing the mean fraction of astrocytes covered by probes for the genes in **(i)** in PFC (red), striatum (yellow), and thalamus (green) from FISH data (**Fig. S10**, n = 4 mice, 2 of each sex, data point shows average of all astrocytes from 2 tissue sections each). Error bars indicate standard error of the mean. See **Methods** for details on quantification, which was performed semi-automatically using CellProfiler. Statistical significance was determined in GraphPad Prism using a repeated-measures one-way ANOVA, followed by Tukey's multiple comparisons test (see **Table S13** for full statistics for each gene). ns, non significant, *$p \leq 0.05$, **$p \leq 0.01$, ***$p \leq 0.001$, ****$p \leq 0.0001$. **D,** Same as **(C)**, for P4 mice (n = 4, 2 of each sex, data point shows average across 2 tissue sections each).

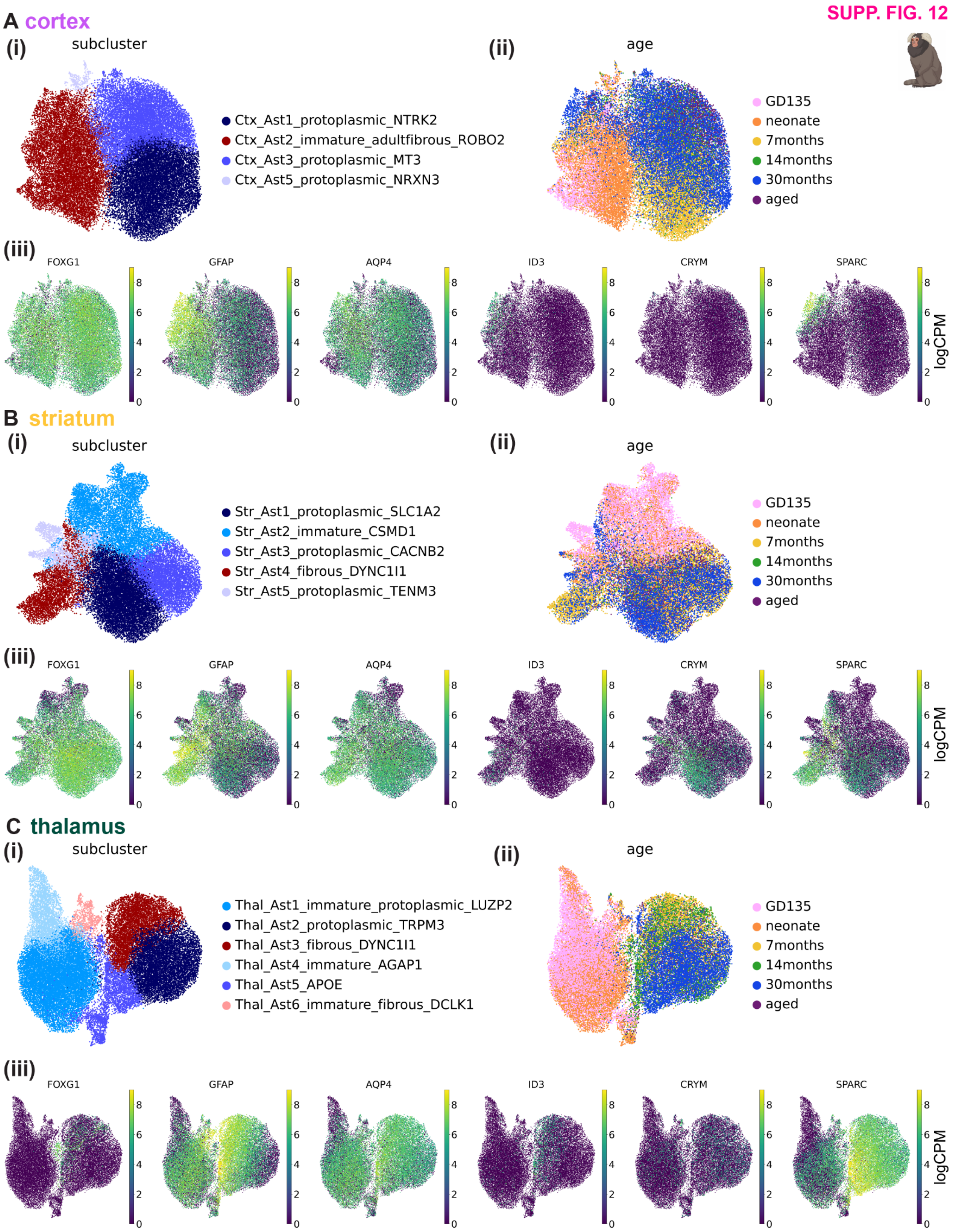


**Supplementary Figure 12. Astrocyte sub-clustering captures intra-regional heterogeneity in marmoset. A,** UMAP embedding of 36,136 marmoset cortical astrocytes from all developmental time points colored by **(i)**

subcluster (see **Methods**), **(ii)** age, or **(iii)** *FOXG1, GFAP, AQP4, ID3, CRYM,* or *SPARC* expression in logCPM units. *FOXG1* marks telencephalic astrocytes, *GFAP, AQP4*, and *ID3* mark fibrous and interlaminar astrocytes[63], *CRYM* marks striatal astrocytes[19], and *SPARC* marks thalamic astrocytes[18]. **B,** Same as **(A)** for 29,931 marmoset striatal astrocytes. **C,** Same as **(A)** for 35,493 marmoset thalamic astrocytes. There are 1,380 *FOXG1+* astrocytes in the thalamus clusters, ~81% of which are assigned to the non-telencephalic astrocyte ABCA subclass, most but not all of which come from the donors from the current study, and may reflect contamination from neighboring regions such as globus pallidus or septum. **D-F**, same as **(A-C)** for mouse astrocytes: **(D)** 25,558 from cortex **(E)** 19,687 from striatum, and **(F)** 23,240 in thalamus. *Top2a* marks immature and SVZ rostral migratory stream astrocytes.

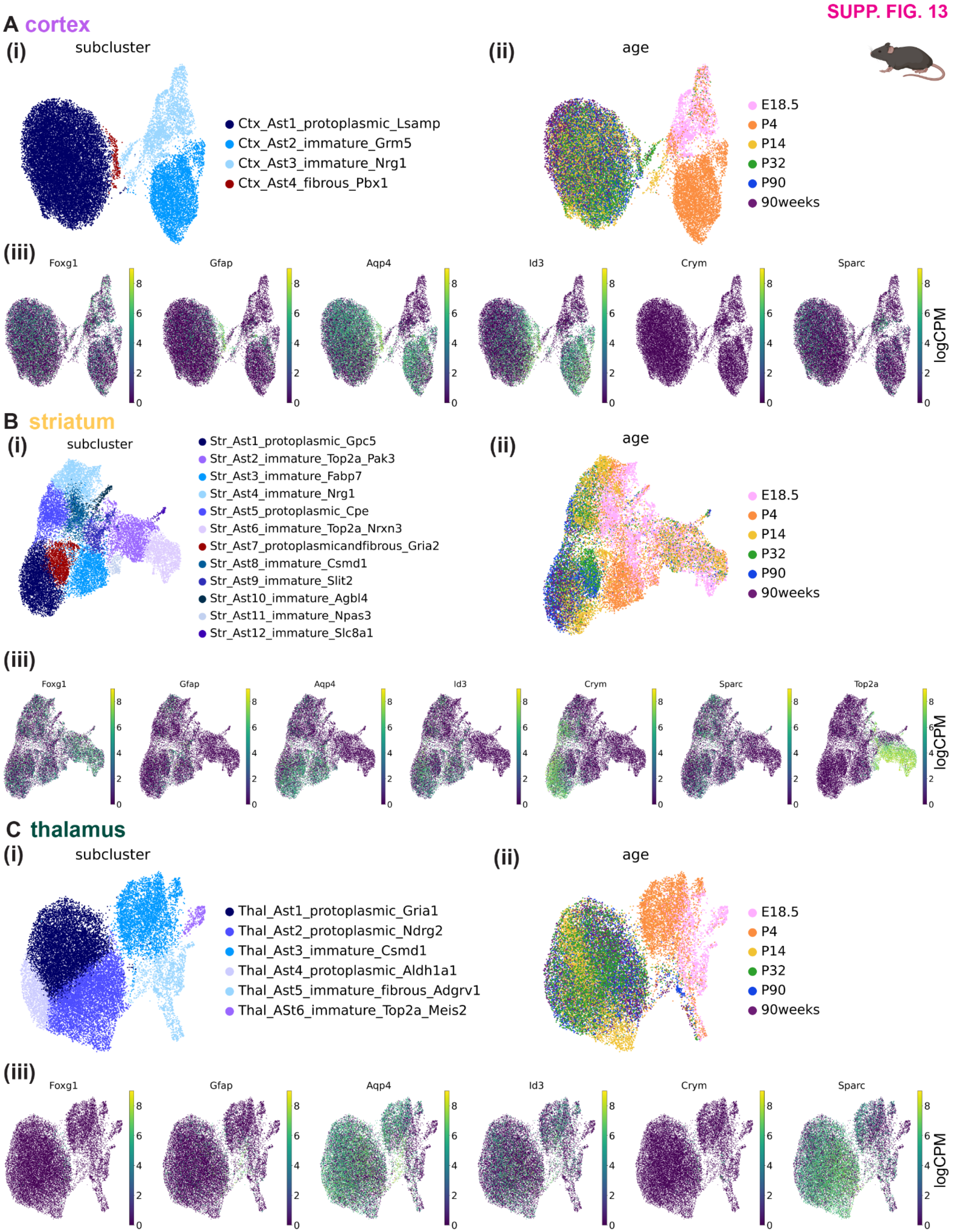


**Supplementary Figure 13. Astrocyte sub-clustering captures intra-regional heterogeneity in mouse. A,** UMAP embedding of 36,136 mouse cortical astrocytes from all developmental time points colored by **(i)** subcluster (see **Methods**), **(ii)** age, or **(iii)** *Foxg1, Gfap, Aqp4, Id3, Crym, Sparc* expression in logCPM units. *Foxg1* marks telencephalic astrocytes, *Gfap, Aqp4*, and *Id3* mark fibrous and interlaminar astrocytes[63], *CRYM* marks striatal astrocytes[19], and *SPARC* marks thalamic astrocytes[18]. **B,** Same as **(A)** for 19,687 striatal astrocytes, with the addition of *Top2a* plotted in **B(iii)**. **C,** Same as **(A)** for 23,240 thalamic astrocytes.

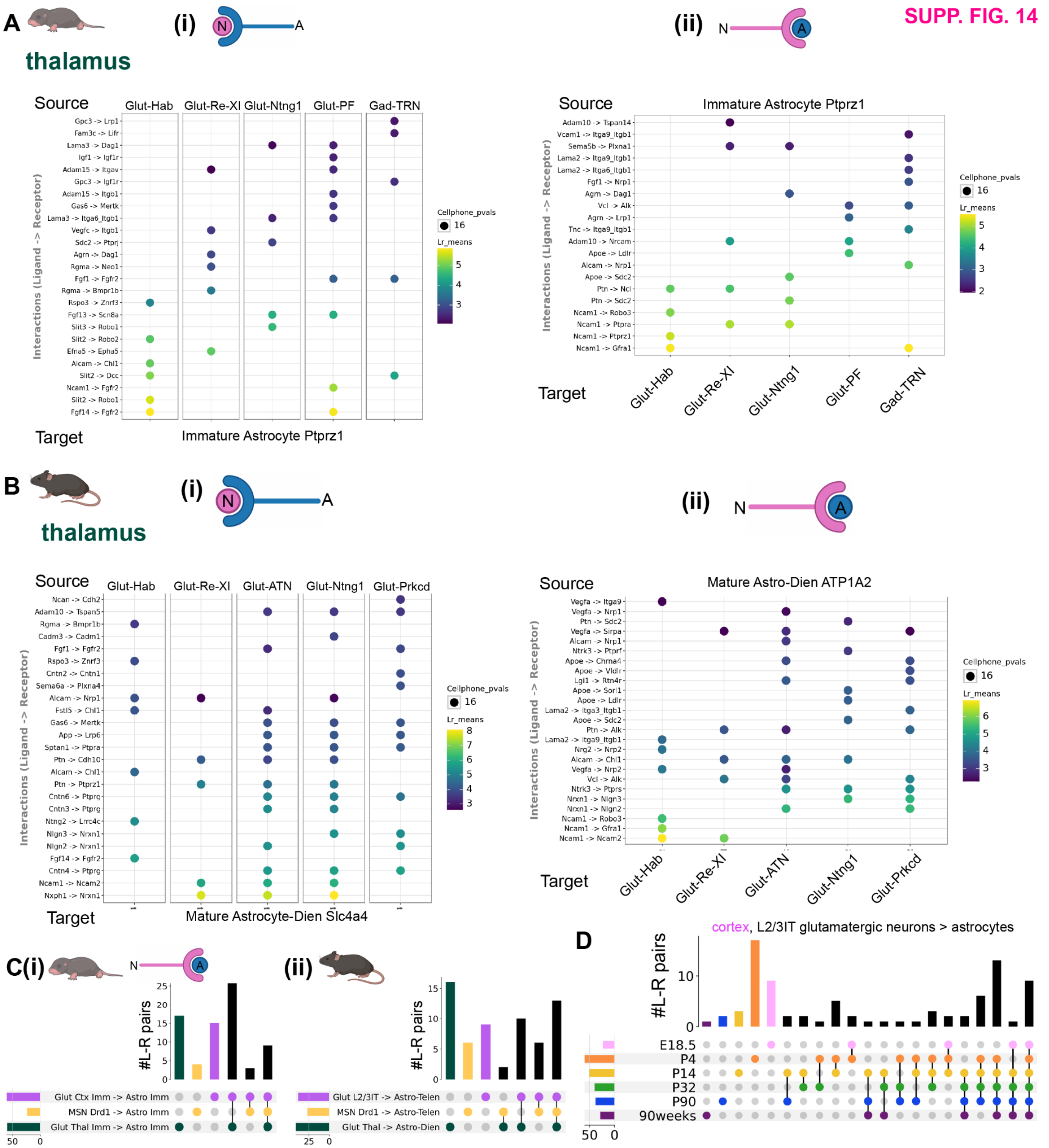


**Supplementary Figure 14. Cell-cell communication analysis for neuron-astrocyte and astrocyte-neuron predicted ligand-receptor pairs across regions and developmental time points in mouse. A,** Dot plot showing magnitude and specificity of the top 25 near-unique (shared with at most one other neuronal cluster) CellPhoneDB-predicted **(i)** neuron-astrocyte and **(ii)** astrocyte-neuron ligand receptor pairs for the most abundant astrocyte and neuronal Leiden clusters in the neonate mouse thalamus. The source cell (top of the plot) expresses the ligand (left side of arrow on the row labels), while the target cell (bottom of the plot) expresses the receptor (right side of arrow on the row labels). The color of the dot indicates ligand-receptor expression magnitude ("Lr_means", calculated as the average of the mean expression of the ligand in the source group and the mean expression of the receptor in the target group), while the size of the dot is inversely related to the p-value on ligand-receptor expression sensitivity (see **Methods**). Glut-Hab, glutamatergic habenula; Glut-Re-Xl, glutamatergic reunions/xiphoid nucleus; Glut-PF, glutamatergic parafascicular nucleus; Gad-TRN, GABAergic thalamic reticular nucleus. **B,** Same as **(A)**, for the P90 mouse thalamus, with the uniqueness criteria relaxed to up to 3 neuronal clusters. Glut-ATN, glutamatergic anterior thalamic nuclei. **C,** UpSet plot showing the number of overlapping neuron-astrocyte predicted ligand-receptor pairs between regions, from the most abundant neuronal and astrocyte subtypes in each region for **(i)** fetal and **(ii)** late adolescent mouse. For cortex (purple), glutamatergic L2/3IT neurons to cortical astrocytes; striatum (yellow), *Drd1+* medium spiny neurons to striatal telencephalic astrocytes; and thalamus (green) thalamic glutamatergic neurons to thalamic astrocytes. Unlike in panel **A**, all L-R pairs meeting minimum expression criteria, including pairs shared with other neuronal and astrocytic clusters, were included in this analysis and that for panel **(D)**. **D**, UpSet plot (as in **(C)**) showing the number of overlapping cortical glutamatergic L2/3IT neuron to cortical astrocyte predicted ligand-receptor pairs between ages.

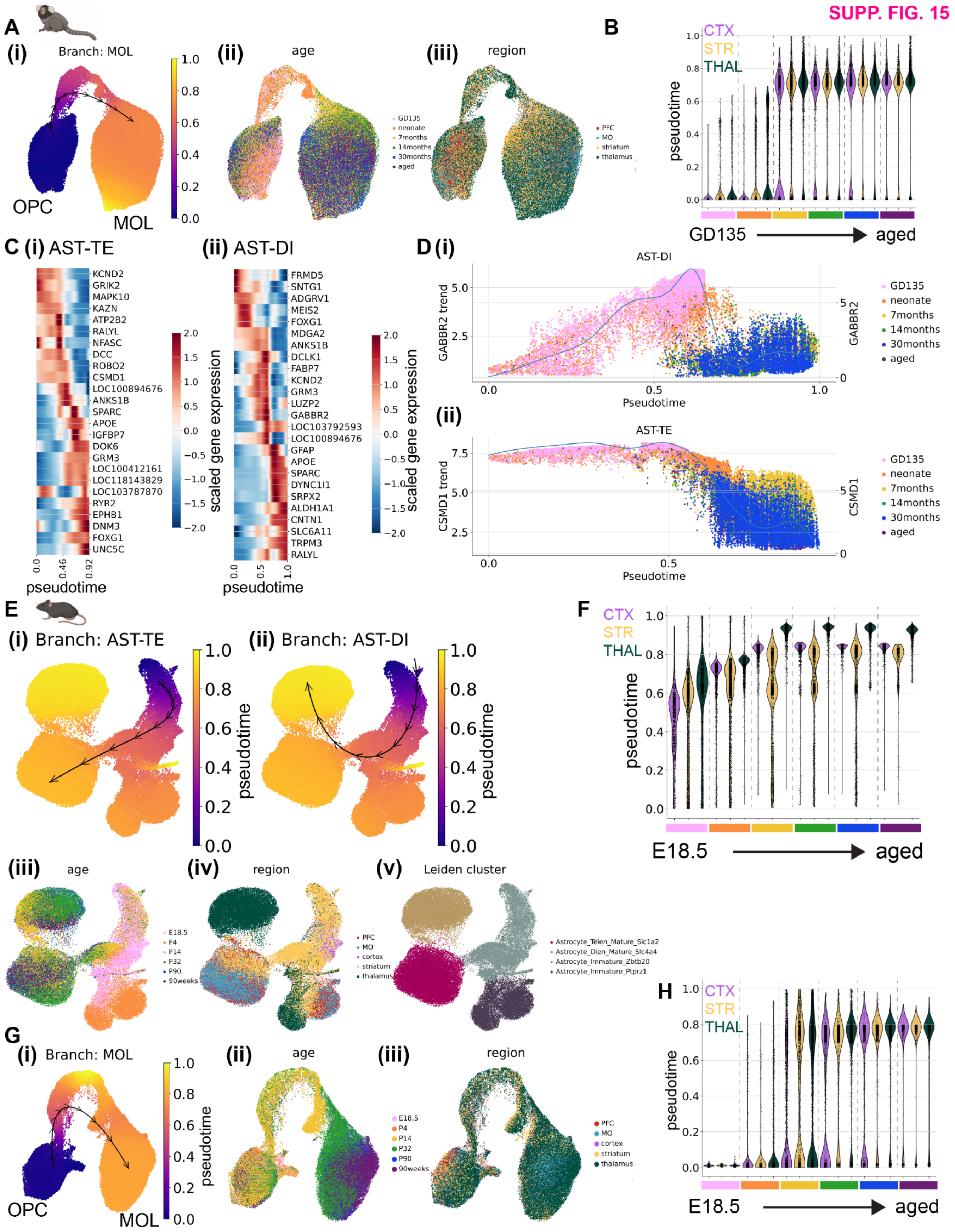


**Supplementary Figure 15. Pseudotime inference in mouse and marmoset oligodendrocyte lineage and mouse astrocytes. A**, Integrated UMAP embeddings of 100,000 (out of 170,786, randomly downsampled)

marmoset oligodendrocyte lineage cells colored by **(i)** Palantir-predicted pseudotime, with trajectory path (black lines and arrows) overlaid, **(ii)** developmental time point, and **(iii)** brain region. **B,** Violin plot (scanpy's default) showing the estimated distribution of pseudotime values for the oligodendrocyte lineage cells in **(A)** grouped brain region and developmental time point. Color scheme for ages as in **(ii)**, with vertical dashed lines indicating divisions between time points. **C,** Heatmap of scaled gene expression over pseudotime for the 25 genes with the highest Mellon change scores for the **(i)** AST-TE and **(ii)** AST-DI branches of marmoset astrocytes. **D,** Individual (single dots) and average (blue line) gene trends (MAGIC-imputed expression over pseudotime) for 2 of the top change-scoring genes for the **(i)** AST-DI (*GABBR2*) and **(ii)** AST-TE (*CSMD1*) trajectory branches. Dots represent single astrocytes colored by developmental timepoint. **E-F,** Same as **(A-B)**, for 73,638 mouse oligodendrocyte lineage cells. **G,** Integrated UMAP embeddings of 68,485 mouse oligodendrocyte lineage cells colored by **(i)** Palantir-predicted pseudotime, **(ii)** developmental time point, and **(iii)** brain region. **F,** Violin plot (scanpy's default) showing the estimated distribution of pseudotime values for the oligodendrocyte lineage cells in **(E)** grouped by brain region and developmental time point. Color scheme for ages as in **(ii)**.

SUPP. FIG. 16

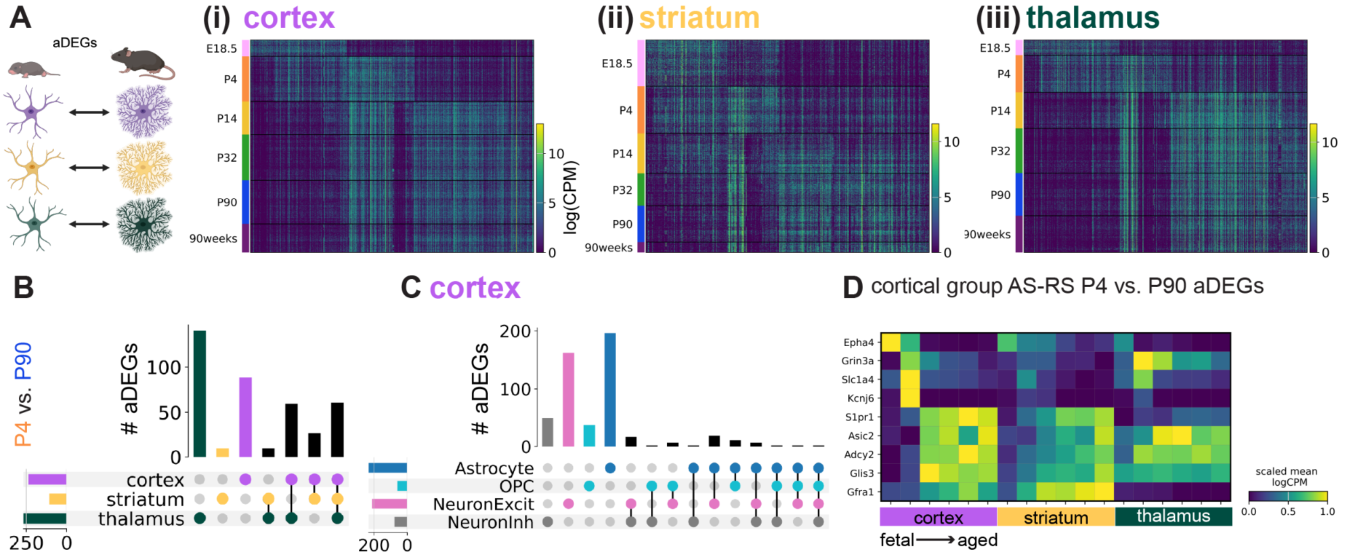


**Supplementary Figure 16. Gene expression signatures underlying the postnatal developmental specification of mouse astrocytes within and across brain regions. A,** Heatmaps (rows corresponding to nuclei and columns to gene) showing expression in logCPM of astrocyte age differentially expressed genes (aDEGs) in astrocytes from **(i)** cortex, **(ii)** striatum, and **(iii)** thalamus, grouped by developmental time point as indicated on the left of the heatmap. The strategy for calculating aDEGs is schematized on the left. **B**, UpSet plot showing the number of overlapping P4 vs. P90 astrocyte aDEGs between cortex, striatum, and thalamus. The colored dots below each vertical bar indicate which region(s) share that set of aDEGs, while the colored horizontal bars indicate the total number of cortex-thalamus aDEGs for each region. Overlap categories with 0 aDEGs are not shown. **C**, UpSet plot (as in **(B)**) showing the number of overlapping P4 vs. P90 cortical astrocyte aDEGs between OPCs (light blue), astrocytes (dark blue), excitatory neurons (pink), and inhibitory neurons (gray). **D,** Matrix plot showing mean expression of selected cortex group astrocyte-specific, region-specific (AS-RS) aDEGs (rows) in mouse astrocytes grouped by region and developmental time point (columns, blocked by region first and then by increasing age within each region block). Expression units of mean logCPM are standardized between 0 and 1 by subtracting the minimum and dividing by the maximum for each trait.

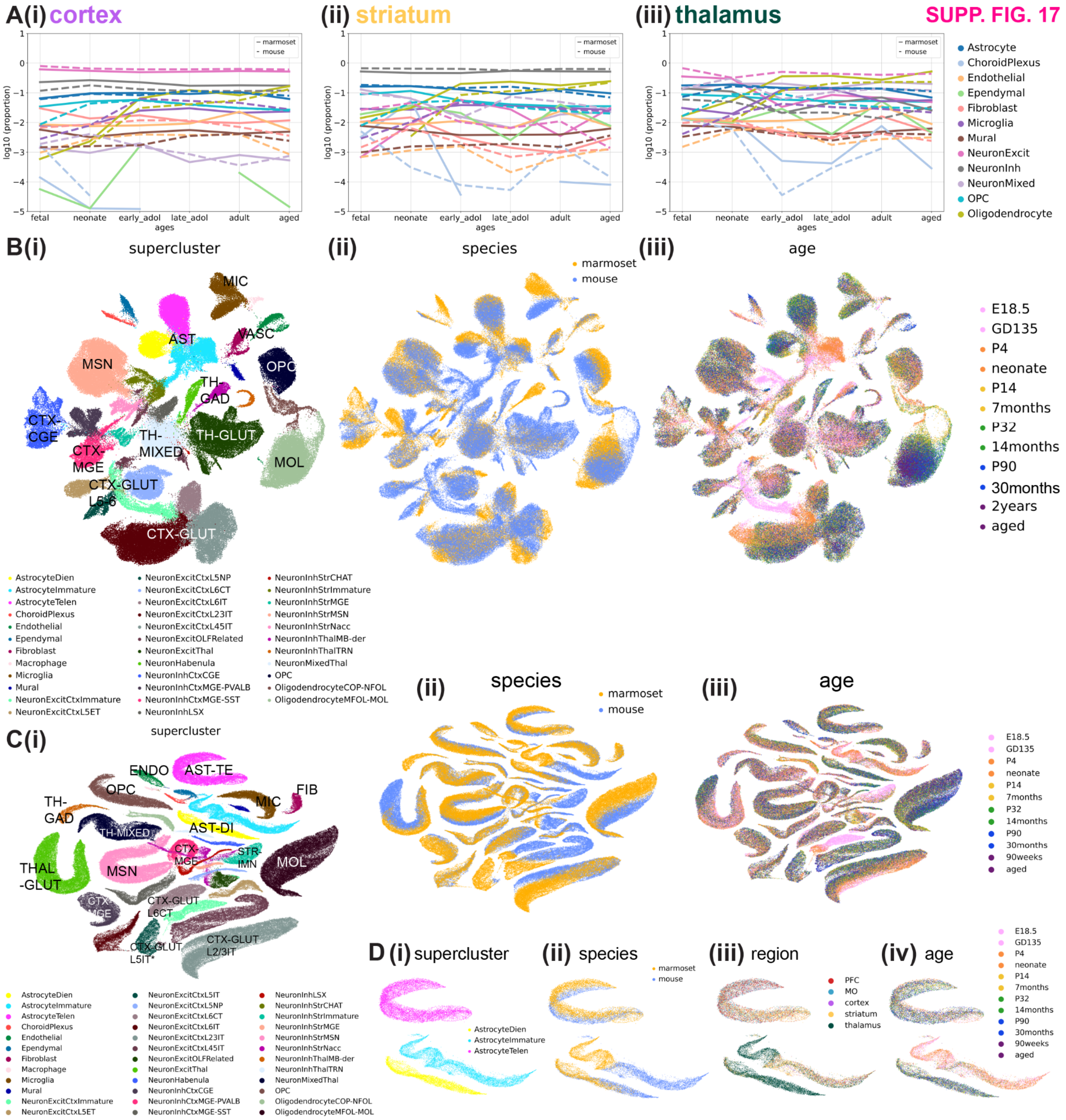


**Supplementary Figure 17. Cell type composition across development in both species and cross-species integration with SATURN. A,** Line graphs showing the proportions of major cell classes in the marmoset (solid lines) and mouse (dashed lines) in log10 scale across developmental time points in the **(i)** cortex, **(ii)** striatum, and **(iii)** thalamus. **B,** scANVI-integrated UMAP embeddings of downsampled marmoset and mouse nuclei, colored by **(i)** supercluster, **(ii)** species, and **(iii)** age. **C,** SATURN-integrated UMAP embeddings of downsampled marmoset and mouse nuclei, colored by **(i)** supercluster, **(ii)** species, and **(iii)** age. **D,** SATURN-integrated UMAP embeddings of marmoset and mouse astrocytes, colored by **(i)** supercluster, **(ii)** species, **(iii)** age, and **(iv)** region.

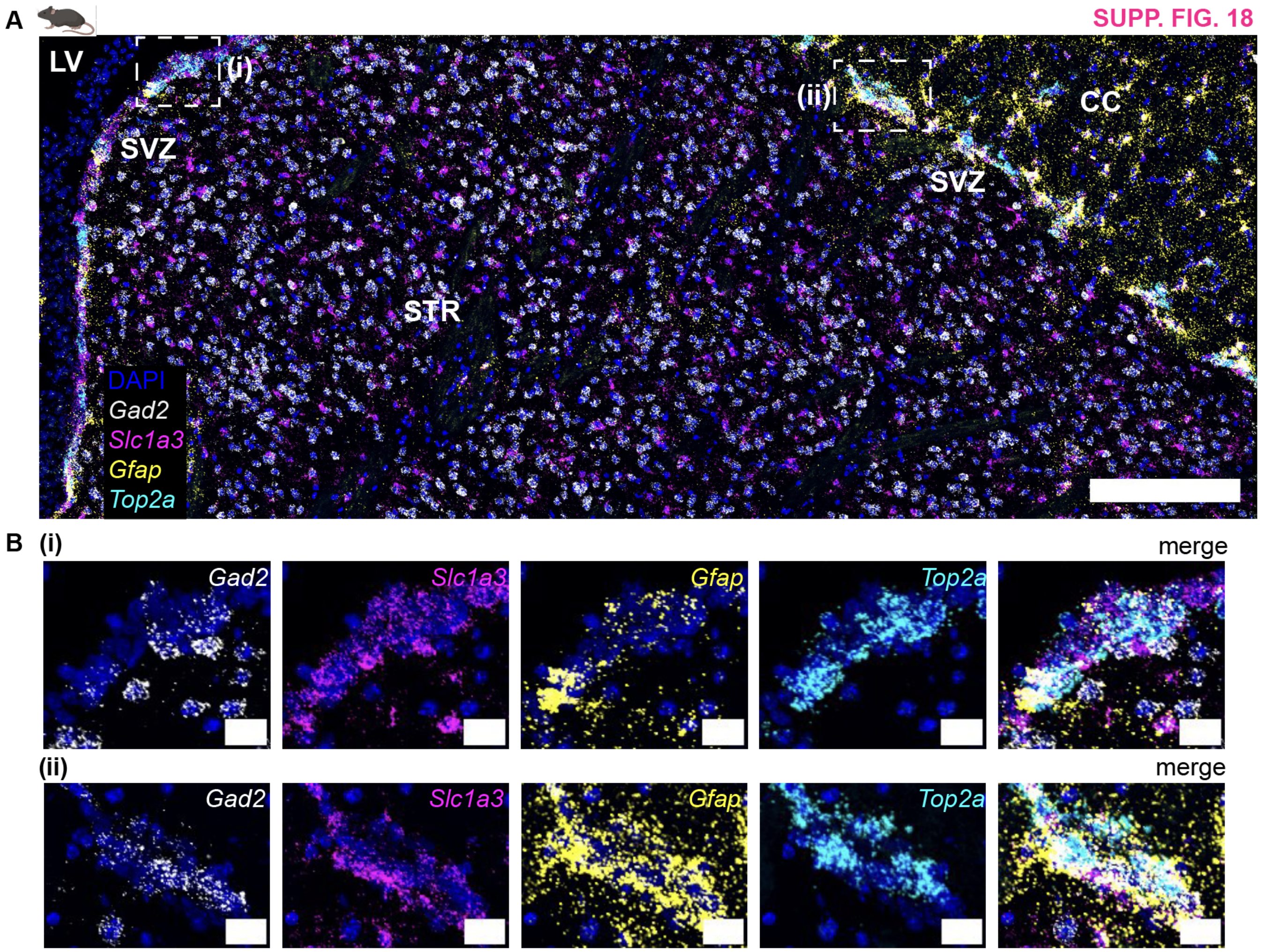


**Supplementary Figure 18. Expression of *Top2a* in astrocytes of the mouse subventricular zone via multiplexed FISH. A,** Composite image of a registered (across imaging rounds), cropped maximum intensity projection of imaged fields of view in the **(i)** PFC **(ii)** striatum and **(iii)** thalamus in one female P90 mouse. Scale bar, 200µm. **B,** Single-channel and composite images of the boxed regions of the subventricular zone in **(A)**. Scale bar, 20µm. LV, lateral ventricle. SVZ, subventricular zone. CC, corpus callosum. Contrast was manually adjusted by setting minimum and maximum intensity values.

**SUPP. FIG. 19**

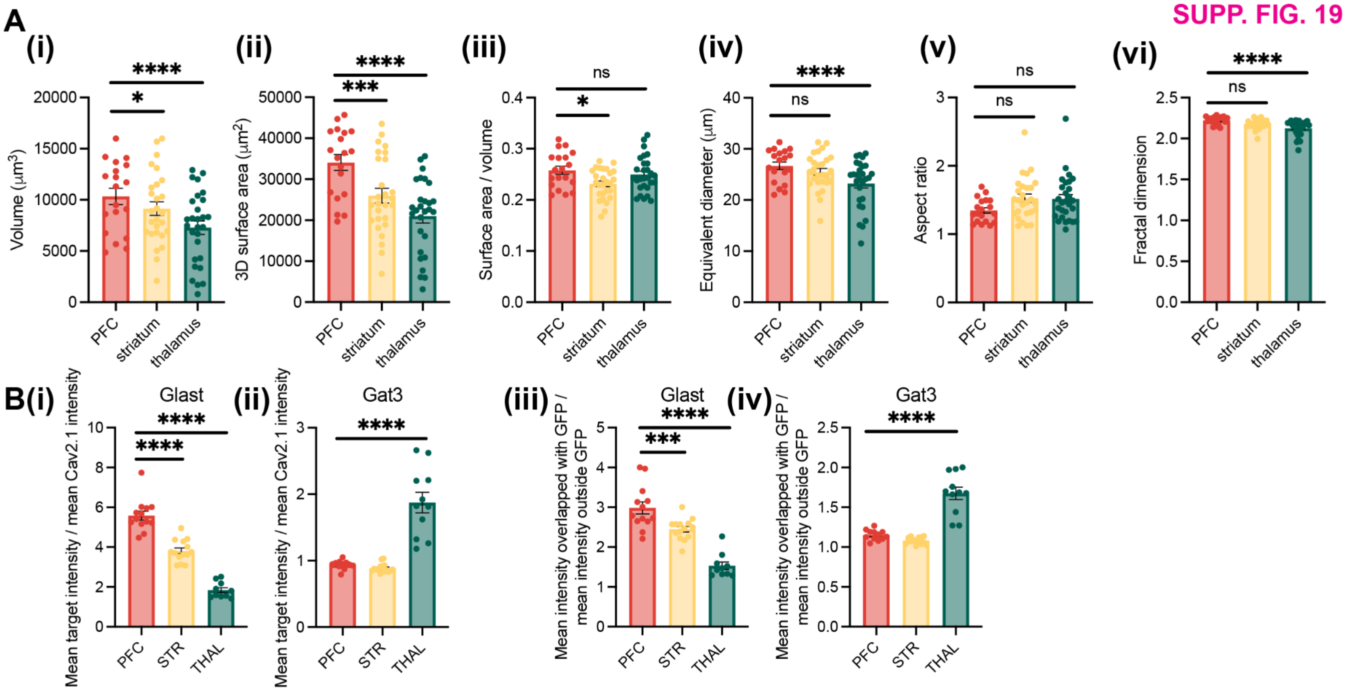


**Supplementary Figure 19. Additional detail on astrocyte morphology and rDEG protein expression differences across brain regions in mouse. A,** Quantitative measures of mouse astrocyte morphology across regions (as in **Fig. 7E**) on a subset of astrocytes with no fractured blood vessels in the field of view and complete capture in the axial dimension (n = 19-28 astrocytes from 3 female and 5 male mice for each region, with statistical significance determined using a linear mixed effects model with “animal” as the random effect group variable). **B,** Bar plots showing **(i, ii)** quantified normalized mean intensity (to Cav2.1 synaptic reference channel) within GFP+ regions and **(iii, iv)** enrichment ratio (mean intensity in GFP+ regions divided by mean intensity outside GFP+ regions) of either **(i, iii)** Glast or **(ii, iv)** Gat3 (from ~18x ExR data as in **Fig. 7F**, n = 12-19 fields of view from 2 mice, with statistical significance determined using a linear mixed effect model as described in the **Methods**). In **A** and **B**, data points are individual fields of view, and error bars are standard error of the mean.

SUPP. FIG. 20

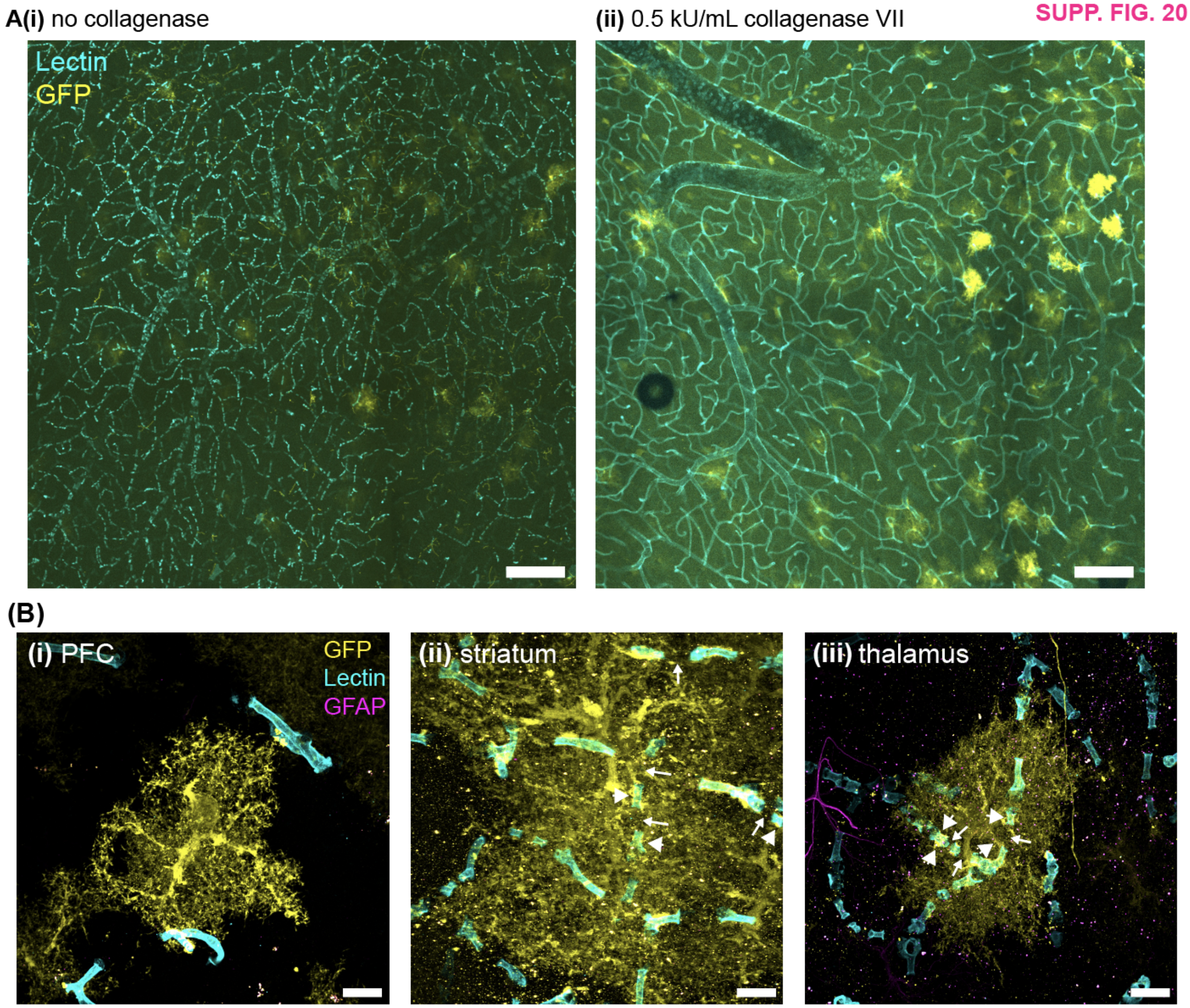


**Supplementary Figure 20. High-concentration collagenase treatment preserves blood vessel morphology in ExR samples. A, (i)** Maximum intensity projection cropped field of view obtained at 4x magnification of ~4x expanded tissue from an Aldh1l1-Cre (Jackson Laboratories # 023748) mouse injected at P1-P2 with PhP.eB CAG-FLEX-GFP and perfused with ExR fixative after 5 weeks. Tissue was sectioned at 80µm, pre-stained for GFP and processed according to the original ExR protocol without collagenase treatment[37]. Blood vessels (cyan, Lectin stain) appear fragmented. This animal is from a litter of mice from a separate experiment whose data was not included in the current paper. **(ii),** Same as **(i)**, for a mouse from a separate experiment whose data was not included in the current paper, sectioned at 150µm, pre-stained for GFP, and treated overnight with collagenase VII at 0.5kU/mL (see **Methods**). Blood vessels appear continuous. Both fields of view are from the thalamus. Scale bar, 420µm in physical units (~100-120µm in biological units, assuming a 3.5-4.2x expansion factor; the expansion factor was lower in the experiment in **(i)**). **B,** Maximum intensity projections of 40x magnification astrocyte fields of view in the **(i)** PFC, **(ii)** striatum, and **(iii)** thalamus from the experiment shown in **(A)(i)**. Fields of view in **(i)** and **(ii)** are from one male mouse, while **(iii)** is from a female mouse. Contrast was manually adjusted in Fiji at or above 35% saturation to show similar GFP brightness across regions. Despite fragmented blood vessels, astrocyte morphology appears almost entirely continuous (compare to **Fig. 7**), except at end feet directly touching blood vessels. Smaller white arrows indicate breaks within blood vessels where astrocyte end feet may exhibit possible local distortion, while larger white arrowheads indicate unbroken blood vessels, where astrocyte processes appear continuous and in good registration with the vessel. Scale bar, 35µm in physical units (~10µm in biological units, assuming a 3.5x expansion factor).

### Methods

#### Marmoset tissue harvest for snRNAseq.

Common marmosets were housed in AAALAC-accredited facilities at MIT, in spacious holding rooms with a 12 hour light/dark cycle, temperature 74.0 ± 2.0°F (23.3 ± 1.1°C), relative humidity of 50 ± 20 %, and unrestricted access to food and water. Cages contained a variety of perches and enrichment devices. Procedures were conducted with prior approval by the MIT Committee for Animal Care (CAC) and following veterinary guidelines. A list of marmosets used in this study and their ages is provided in **Table S1**. Marmosets (GD135 - 13+ years old, 10 individuals for snRNAseq and 4 individuals for FISH), were euthanized and brains harvested as previously described[7]. Marmosets were generated from a total of 14 breeding pairs. Briefly, animals were deeply sedated by intramuscular injection of ketamine (20–40 mg/kg) or alfaxalone (5–10 mg/kg), followed by intravenous injection of sodium pentobarbital (10–30 mg/kg). When the pedal with-drawal reflex was eliminated and/or the respiratory rate was diminished, animals were trans-cardially perfused with ice-cold sterile PBS. Whole brains were rapidly extracted into fresh PBS on ice.

After transporting the brain to the lab on wet ice, the brain was sectioned into coronal blocking cuts (slabs, 2-8mm in thickness) using a chilled custom-designed marmoset brain matrix[7]. Surgical tools were autoclaved and allowed to cool before use. All tools, the matrix, and the dissecting block were cleaned with RNase Zap™ wipes (ThermoFisher) prior to each dissection. Slabs were transferred to a pre-chilled dissecting block and regions were dissected using a marmoset atlas as reference[103] (**Table S27**). The areas targeted for prefrontal cortex includes areas 8, 9, 10, 11, 47L, 14R, 46, 47, 13, 32, and 45; areas 6M, 6DC, 4c and 4ab for motor cortex; caudate and putamen for striatum, and all thalamic nuclei except posterior regions of the pulvinar and lateral geniculate nucleus for thalamus. Fetal and neonate brains were not dissected using the brain matrix due to their small size. Instead, the brains were hemi-sectioned, placed on a cooled dissecting block, and the prefrontal cortex and motor cortex (only for neonate, not dissected at GD135) were scooped from the surface of either hemisphere using anatomical landmarks. Two large (several mm) coronal slabs approximately spanning from the anterior beginning of the temporal lobe to its posterior end were cut using a razor blade, and the striatum and thalamus were dissected from the anterior and posterior slabs respectively (**Fig. S5A-B**). For one neonate replicate (21-197), the brain was frozen and stored at −80°C for several months, placed at -20°C overnight prior to the day of dissection, and thawed on ice prior to dissection. Dissected tissue was transferred to chilled 1.5mL microcentrifuge tubes, snap-frozen in liquid nitrogen, and stored at −80°C until nuclei isolation. Dissections began within 90 minutes of euthanasia and were performed in a median time of ~40 minutes (range 30-80 minutes).

#### Mouse tissue harvest for snRNAseq.

Animal work was performed in accordance with protocols approved by MIT's Committee on Animal Care and NIH guidelines. All postnatal mice were wild-type C57BL/6J originally obtained from Jackson Laboratories and bred in-house. Timed pregnant C57BL/6J females were either obtained from Jackson Laboratories to arrive between gestation day 11 and 15 or were impregnated in house by setting up overnight mating pairs with females in proestrus or estrus phase. Embryos were harvested at E18.5 (18 days after the plug date). Mice were housed in a facility with a light cycle running from 07:00 to 19:00, temperature 20-22.2°C, humidity 30-70%, and food and water available *ad libitum*. Postnatal mice were not derived from timed pregnant females. Instead, age was determined during regular pup checks by experienced researchers based on the Jax Mice Pup Appearance Chart. Thus, ages are approximate within +/- 0.5 days for P4 neonates, within +/-1 day for P14 early adolescents, within +/- 3 days for P32 juvenile mice and P90 young adult mice, and within +/-1 week for aged mice (90 weeks). Except for the P32 and aged time points, mice were obtained from different litters, and minimal replicate effects were observed in the snRNAseq data, suggesting adequate matching of developmental time points across replicates. A list of mice used in this study is provided in **Table S1.**

Non-neonate animals were acclimated to the lab space for at least 30 minutes prior to beginning euthanasia. Euthanasia took place between 9am-12pm to control for circadian rhythm effects, with a maximum of four animals processed per batch. Non-neonate animals were deeply anesthetized with isoflurane and decapitated. Heads were briefly submerged in liquid nitrogen for 3 seconds. Neonates were anesthetized via hypothermia and decapitated. Surgical tools were autoclaved and allowed to cool before use. All tools, brain mold, and the dissecting block were cleaned with RNase Zap™ wipes prior to each dissection. Brains were harvested and sagittally sectioned at 1mm thickness for a total of 2 mm from the midline for either hemisphere (total of four ~1mm slices), on a brain mold using chilled razor blades. For P4 animals, two ~2mm sections from the midline were used. Tissue was dissected from slices on a chilled dissecting block exposed to room air, using a dissecting microscope at 1.6X magnification. Regions of interest were identified using the Allen Institute reference brain atlas at the appropriate time point. Dissected tissue was placed in cooled 1.5mL microcentrifuge tubes and spun down in a tabletop mini centrifuge prior to snap freezing in liquid nitrogen before storage at -80°C until nuclei isolation. 2-3 neonates were pooled in each tube. Time from decapitation to snap freezing ranged from 7-13 minutes per animal. Samples from at least 3 mice (at least 1 female) are represented at each developmental time point (except 90 weeks, which is missing a male donor for PFC, see **Table S1**) and for each brain region. However, due to failures during microdissection, nuclei isolation, and 10x Genomics chip running, not all biological replicates are balanced across brain regions (e.g., some replicates have only one or two brain regions present).

For embryonic brain microdissection, the pregnant dam was deeply anesthetized with an overdose of isoflurane, decapitated, and placed on a cooled dissecting block. The abdomen was opened and placentas were removed from the abdominal cavity. Embryos were harvested from the placenta and rapidly decapitated one-by-one. Heads were frozen on metal disks over dry ice for 5-10 minutes until frozen solid, stored at -20°C for 1.5 hours prior to microdissection. Heads were cut approximately in half using a small mouse brain mold and placed on a dry-ice cooled metal platform. Regions of interest were dissected using a tissue punch (1.27mm Ted Pella MilTex Biopsy Punch with Plunger, 15110-10) to extract tissue from most medial surface on either hemisphere. 2-3 embryos were pooled in each tube. The other dissection procedures were the same as described above. Dissections were performed in less than 20 minutes per set of tubes from decapitation to snap freezing.

**Nuclei isolation and single-nucleus RNA sequencing.**

Nuclei were extracted from frozen tissue using the 10x Genomics Chromium Nuclei Isolation Kit (Protocol CG000505, Rev A). Manufacturer instructions were followed with the following notable exceptions: 1) Most samples were dissected and frozen directly in Sample Dissociation Tubes (omitting step e), 2) Total lysis time was decreased to 10-14 total minutes of incubation in the lysis buffer (longer for marmoset and larger tissue chunks) from when lysis buffer was first added to the first sample (effectively shortening protocol step h) before proceeding to step i; 3) Tissue mass was larger than the 45mg upper limit recommendation for some marmoset samples; 4) Lysis buffer was supplemented with Roche Protector RNase inhibitor at 0.2U/uL, and 5) if no pellet was visible following any centrifugation steps, ~200uL or less of supernatant was retained for samples with a visible pellet or debris. For the final resuspension step (step s), if no pellet was visible, less than ~40-100uL of supernatant was retained at the bottom of the tube and no additional volume was added prior to nuclei counting. To avoid large clogs, large chunks of marmoset tissue were split in half and processed in parallel for nuclei isolation, and re-pooled prior to 10x Chromium chip loading. For nuclei isolation, a maximum of 4 samples were processed in series by a single researcher in a given preparation (usually 8 samples total, with 2 researchers in parallel). For tissue dissociation (step f), samples with a small amount of tissue (~20mg or less) were processed (transferred to wet ice, coated with 200uL lysis buffer, and dissociated with pestle) one at a time. For dissociating larger amounts of tissue (larger than ~20mg) that required some thawing before pestle dissociation, samples were transferred to wet ice and coated with 200-300uL of lysis buffer in parallel, and then homogenized with the pestle one at a time.

Nuclei concentration was quantified by staining suspensions with DAPI, loading on a C-Chip hemocytometer, imaging on a fluorescence microscope, and using the Fiji 3D Object Counter plugin to automatically quantify the number of nuclei within 4 large grid squares (0.8uL of volume). Debris was assessed by comparing the signal in bright field to the signal in DAPI and nuclei quality was assessed by examining DAPI-stained nuclei at 40-60x magnification prior to starting the 10x Genomics Chromium snRNAseq protocol. Nuclei suspensions with an unacceptable amount of debris (i.e., large clumps in bright field that were not DAPI+) and/or blebbing (i.e., with most nuclear membranes appearing substantially disrupted) were discarded. Because marmoset tissue is precious, a higher level of debris and/or blebbing was tolerated for marmoset nuclei suspensions. Nuclei suspensions were diluted to a target concentration of 1,000 nuclei/uL for 10x Genomics Chromium chip loading.

snRNAseq libraries were prepared using 10x Genomics Chromium Next GEM Single Cell 3' Reagent Kits v3.1 (Protocol CG000315 Rev C or Rev D) following manufacturer instructions. Time from tissue lysis to 10x Chromium chip loading averaged ~2 hours. Whenever possible, channels were loaded with enough nuclei suspension to recover a target of 10,000 nuclei. Initial cDNA amplification was performed using 13 PCR cycles and Sample Index PCR was performed using 11-12 PCR cycles. Amplified cDNA (product of protocol step 2) and libraries (product of protocol step 3) were quantified and quality-checked using both Qubit (HS dsDNA Assay) and a Fragment Analyzer. Libraries were pooled and sequenced on an Illumina NovaSeq at the Broad Genomics Platform. Data are available for download at: https://data.nemoarchive.org/biccn/grant/u01_feng/feng/transcriptome/sncell/10x_v3.1/.

**Read alignment.**
Reads were aligned to an optimized mouse reference genome based on the mouse GRCm38 primary sequence assembly (version 2)[101] or or a modified marmoset mCalja1.2.pat.X assembly (https://www.ncbi.nlm.nih.gov/datasets/genome/GCF_011100555.1/) provided courtesy of Michael DeBerardine and Fenna Krienen (Princeton Neuroscience Institute) in which mitochondrial genes from CM021961.1 (https://www.ncbi.nlm.nih.gov/nuccore/1820101357/) were annotated using MITOS2 (https://doi.org/10.1093/nar/gkz833). The marmoset reference genome was generated from the .fasta and .gtf files using Cell Ranger "mkref" v7.1.0 using default parameters without filtering for any gene/transcript biotype. 10x Genomics Cell Ranger software version 7.1 was used for alignment and counting via the 10x Genomics Cloud Analysis platform. For samples that were sequenced across several library pools, fastq files were grouped prior to alignment to create one cell-by-gene (cell x gene) counts matrix per sample. CellBender (v0.2.0) remove-background[104] was used to remove ambient RNA and call nuclei with default parameters and expected_cells = 10,000, total-droplets-included = 40,000 and the –cuda flag. CellBender-cleaned cell x gene matrix .h5 files were read into Python in the anndata[105] format using a custom function written by Stephen Fleming (https://github.com/broadinstitute/CellBender/issues/57).

**Calculation of sequencing coverage statistics.**
To determine whether we achieved our target of 40,000 sense reads per nucleus and calculate the sequencing coverage statistics shown in **Fig. S1** and **Table S28**, we ran a light quality control on CellBender-cleaned cell x gene matrices. Briefly, nuclei with fewer than 1,000 unique molecular identifiers (UMIs) and fewer than 800 genes expressed were removed, as were genes with nonzero expression in fewer than 10 cells. We note these cutoffs are more stringent than what we used for preprocessing (see "snRNAseq data preprocessing and quality control" section), to account for the lack of doublet removal and additional manual curation that is much more time consuming. Nuclei with greater than 4% of reads aligning to the mitochondrial genome (prefix "mt-" for mouse or "MT-" for marmoset) were removed.

**Sex determination in mouse and marmoset fetal and neonate samples.**
Because 2-3 mouse E18.5 brain regions were pooled into a single tube for generating the nuclei suspension without sex determination, we do not have metadata about sex for these samples. Instead, we performed sex assignment on a per-nucleus basis after snRNAseq based on the expression of Y-chromosome genes. Specifically, if a nucleus had *Zfy1*, *Zfy2, Usp9y*, *Uty*, *Eif2s3y*, *Kdm5d*, or *Ddx3y* expression above 2 log counts per million, it was assigned male sex. This resulted in 41.37% male nuclei for the E18.5 samples, likely an underestimate due to dropout.

Marmoset GD135 donors and one neonate donor (21-197) did not have their sex determined anatomically prior to euthanasia. To determine their sex, we performed PCR-based sex genotyping on either skin or brain tissue. Briefly, DNA was extracted from tissue using the NucleoSpin Tissue kit (Macherey-Nagel) and eluted in nuclease-free water. We genotyped for *ZfX/Y*[106] and *SRY*[107] using the following PCR primers (from 5' to 3'):

- ZfX/Y Forward (modified from the original publication[106]): CTGTGCATAACTTTGTTCCTG

- ZfX/Y Reverse (modified from the original publication[106]): CAGTTGCCTTTGTCATCATC
- SRY Forward: TACAGGCCATGCACAGAGAG
- SRY Reverse: CTAGCGGGTGTTCCATTGTT

And ran the following protocol on a thermocycler:

1. 94°C - 2 min
2. 98°C - 10 sec
3. 58°C - 30 sec
4. 68°C - 40 sec
5. 35 or 40 rxns (34x or 39x)
6. 68°C - 3 min
7. 4°C - Infinite

We then digested 10uL of the ZFX/Y product using the DdeI and MseI enzymes in separate reactions (New England Biolabs). We were able to see clear separation of bands on a 2% agarose gel run at 135V for 40-45 min. Digesting the ZfX/Y PCR product with DdeI will yield double bands if the animal is XX and triple bands if the animal is XY. Digesting the ZfX/Y PCR product with MseI, which can only cut the ZfY band, will show smaller bands below the ~500bp PCR product for XY animals and a single band for XX animals. An *SRY* band of around 218bp indicates male sex. Our predicted sexes also matched *SRY* gene expression after snRNAseq.

**snRNAseq data preprocessing and quality control.**

All scripts used for preprocessing and downstream analyses are available on GitHub at https://github.com/Feng-Lab-MIT/AstrocyteHeterogeneity. Preprocessing was conducted on a species-wide, cross-age, cross-region basis. Filtered counts matrices were pre-processed using scanpy[108]. Briefly, nuclei with fewer than 800 unique molecular identifiers (UMIs) and fewer than 500 genes expressed were removed, as were genes with nonzero expression in fewer than 10 cells. Nuclei with more than 4% of counts annotated as mitochondrial genes (prefix "MT-" in marmoset or "mt-" in mouse) were removed. Counts were normalized to 1 million per cell and log-transformed using scanpy's "log1p" function. Highly variable genes (HVGs) were identified from raw counts on a per-batch basis using scanpy's "seurat_v3" method with 4,000 top genes. HVGs present in less than 10% of batches or of mitochondrial origin (prefix "MT-" in marmoset or "mt-" in mouse) were removed. We used the scanpy-based single cell variational inference (scVI[109]) package to create and train a variational autoencoder on a subset of the cell x gene matrix with highly variable genes only with the following parameters: batch_key corresponding to 10x genomics reaction, raw counts layer, "gene-batch" dispersion, and training with GPU. The resulting nonlinear embedding was used to create a neighborhood graph for clustering and calculate UMAP[110] coordinates with the scanpy "sc.pp.neighbors" and "sc.tl.umap" functions. scVI and the scvi-tools package were the basis for many downstream analyses. We used Solo, an automated doublet removal package[111] based on the scVI model, to calculate doublet scores for each nucleus on a per-batch basis. For the mouse data, one nucleus was removed from the "Exp074_mmP35_2D" 10x reaction to circumvent a known bug in the Solo package (https://discourse.scverse.org/t/solo-scvi-train-error-related-to-batch-size/1591/2). Doublet/singlet thresholds were determined manually by examining the doublet- vs. singlet-score scatter plot and a predicted doublet rate of 13-15%, a conservative estimate based on 10x Genomics' predicted ~8% for a target recovery of 10,000 nuclei. We decided to use this manual threshold to prevent removal of developing cells that are more likely to be flagged doublets automatically.

**Global (species-wide, cross-age, cross-region) integration and annotation.**

After automated doublet removal, highly variable genes were re-calculated and the scVI model was re-trained. Leiden clustering was performed with a resolution of 1. Top differentially expressed genes in each cluster (i.e., putative marker genes) were identified using scanpy's rank_genes_groups function with log-normalized counts and the Wilcoxon rank-sum method. Top marker genes, expression of known marker genes, and dendrograms were used to annotate clusters per the following convention: [Cell class]_[Excit or Inh*]_[region*]_[cortical layer*]_[age*]_[known marker*]_[first rank_genes_group marker]-[second rank_genes_group marker*], where the asterisked attributes were used variably, as applicable. Low-quality clusters were manually identified and removed, highly variable genes re-calculated, the scVI model re-trained, and clustering and annotation processes were repeated. Remaining doublet clusters were manually removed, and the neighborhood space and UMAP coordinates were recalculated. Neuronal annotations were subsequently refined based on predicted mapping to the Allen Brain Cell Atlas using MapMyCells[41] (see **Alignment to Allen Brain Cell Atlas with MapMyCells** section below).

Adult (29-32 month, referred to as 30 month in most of the paper) marmoset data used in this study was generated previously[7], and includes data from 4 donors. CellBender background-removed cell x gene matrices for the four regions of interest (annotated as “pfc”, “m1”, “striatum”, and “thal”) were preprocessed as described above, except that mitochondrial genes were not annotated and therefore not used for quality control. Adult marmoset cell x gene matrices were randomly downsampled to 40,000 nuclei per region to match the approximate number of nuclei in each age-region combination in the developmental dataset. Of note, the adult marmoset snRNAseq reads were aligned to the cj1700 transcriptome, which lacks mitochondrial genome annotation (https://www.ncbi.nlm.nih.gov/datasets/genome/GCF_009663435.1/). 22,582 genes overlapped between the mCalJa1.2.pat.X (developmental and aged data, 31,308 genes) and cj1700 (adult data, 27,304 genes) reference-aligned datasets. Data from GD135, neonate, 7-month, 14-month, and aged timepoints were integrated with the downsampled adult (29-32 month) data using scVI using only highly variable genes, clustered, and annotated and described above. Clusters with the vast majority or all nuclei derived from adult (29-32 month) marmoset data were removed, as they likely derive from differences in dissection strategies between the two studies. We also removed a small subcluster of 233 adult astrocytes that clustered with immature astrocytes, because they derived primarily from a single adult replicate and were not found in other donors 14 months and older. Neuronal annotations were subsequently refined based on predicted mapping to the Allen Brain Cell Atlas using MapMyCells[41] (see **Alignment to Allen Brain Cell Atlas with MapMyCells** section below). In designing and implementing downstream analyses (e.g. compositional, pseudotime, and cell-cell interaction analyses), we relied heavily on the Single Cell Best Practices e-book (https://www.sc-best-practices.org/)[112].

**Integration quality analysis.**
We utilized code from ref[38] to quantify integration quality separately for each species using neighbor consistency, average silhouette width, and donor mixing. Neighbor consistency relied on coordinates in a pre-integration embedding and post-integration embedding, for which we used scanpy’s PCA function (50 components) to recalculate the pre-integration embedding and scVI for the post-integration embedding. We calculated the average silhouette width with a wrapper function of the sklearn package with the scVI embedding, and used “supercluster” as the group labels. Finally, donor mixing was determined using the “seurat-alignment_score” function with the scVI embedding and donor ID.

**Alignment to Allen Brain Cell Atlas (ABCA) with MapMyCells[41].**
Both mouse and marmoset mature neurons (GABAergic, glutamatergic, and mixed from mouse P14 and above or marmoset neonate and above) or astrocytes were subsetted from the larger cell x gene matrix of each species by age and region, with some metadata removed to shrink the file size. To use the maximum number of genes available for each marmoset time point, pre-integrated adult or developmental gene counts (that is, with ~27,000 genes for 29-32 month marmoset data and ~31,000 genes for developmental and aged data) were used as input. Per the MapMyCell input requirements (https://portal.brain-map.org/explore/file-requirements-and-limits), cell x gene matrix entries were set to raw counts and NCBI gene symbols were converted to mouse Ensembl gene IDs using g:Profiler’s g:Convert[113]. Marmoset NCBI gene symbols were converted to mouse NCBI gene symbols using a table downloaded from Ensembl BioMart (https://useast.ensembl.org/info/data/index.html)[114], available in **Table S29**. If no match was found in the table, the marmoset gene name was converted to sentence case. Due to non-uniqueness after converting marmoset IDs to mouse, *Bex2* was removed from the adult marmoset neuron cell x gene matrix, while *Cstl* was removed from the developmental and aged marmoset neuron cell x gene matrices, before mapping to Ensembl IDs. ~25,300 mouse and ~14,800 marmoset genes were successfully mapped to Ensembl IDs, and the others were excluded from MapMyCells analysis. These smaller (<2Gb) cell x gene matrices were uploaded to the Allen Brain Maps’s MapMyCells web portal (https://knowledge.brain-map.org/mapmycells/process/) and aligned to the 10x Whole Mouse Brain (CCN20230722) reference taxonomy[1] with the hierarchical mapping algorithm. Using the output of MapMyCells, which is a “class”, “subclass”, “supertype”, and “cluster” assignment for each barcode, we examined the most abundant (as the percentage of our cells in each of our clusters mapping to that subclass)

ABCA subclass (about the same taxonomic rank) for each of our cross-region, cross-age, within-species embedding Leiden clusters. Broadly, our annotations were identical or highly similar to the most abundant ABCA mappings. When our annotation disagreed with or was not as specific as the most abundant ABCA subclass mapping, and we were not confident in our annotation, we updated the annotation based on the most abundant ABCA subclass (e.g., comprising over 60% of the cells in the cluster for mice, or over 20% for marmoset). For clusters that had a more uniformly distributed mapping onto ABCA subclasses (e.g., less than 10% mapping onto each subclass), we examined several of the top mapping clusters and their anatomical locations (using the web resource from ref[1] available at https://knowledge.brain-map.org/data/5C0201JSVE04WY6DMVC/summary), and if they were in the same taxonomic or anatomical neighborhood, annotated our cluster accordingly. In a few cases, such as for marmoset thalamic neurons mapping to a midbrain ABCA population and immature astrocytes mapping to the Allen olfactory bulb/immature neuron subtype, we did not adopt the ABCA subclass label. The class, subclass, supertype, and cluster mappings for each of our leiden clusters (as proportion of cells in that cluster mapping to each ABCA taxonomic rank) for both species are provided in **Table S4.**

**Compositional analysis with scCODA[42].**

The proportional breakdown of each leiden cluster by developmental time point (age), brain region, and sex are provided in **Table S3** (assigned brain region) and displayed in **Fig. S4** (dissected brain region). To more quantitatively assess cell type composition changes across these variables, we used the scanpy-based single-cell compositional data analysis (scCODA)[42] (https://github.com/theislab/scCODA), which implements a Bayesian model of cell type counts to address the issue of low sample sizes in snRNAseq data. For this analysis, we merged the PFC and MO into one “cortex” assignment, due to their high degree of similarity in cell type composition. Per the tutorial in the Single Cell Best Practices e-book (https://www.sc-best-practices.org/)[112] Section 17.4, we generated an scCODA model of type “cell_level” with the cell type identifier as either “leiden” (cluster) or “cell_type”, sample identifier as “10x_batch” and “sex” and age, region, sex, and replicate as covariate observation. We ran the model with the formula “region + age + sex”, automatic selection of reference cell type for leiden-level analysis and Mural (marmoset) or Astrocyte (mouse) as the reference cell type for cell type-level analysis, and default false discovery rate of 0.05. The “final parameter”, which is a boolean value that indicates whether or not there is a significant effect of age, region, or sex on the composition of each cell type and cluster for both mouse and marmoset are provided in **Tables S5-8.**

**Region reassignment of cross-contaminant nuclei in fetal and neonate marmoset.**

We observed a modest amount of cross-region contamination, particularly between striatum and thalamus in our late embryonic and neonate samples, which is not unexpected given our coarser dissection strategy for these regions (**Fig. S5-6**). For example, in marmoset, 4% of medium spiny neurons came from the thalamic dissection and 23% of thalamic excitatory neurons, which were mostly from GD135 and neonate time points, came from the striatal dissection. To reduce the effect of this cross-contamination on downstream analyses, we reassigned the region annotation for neurons, astrocytes, and OPCs (the most strongly region-segregated cell types on which we focused our analysis) at these ages. We subsetted the cell type of interest from the cross-age, cross-region integrated cell x gene matrix, re-computed neighbors using the scVI latent space, and recomputed the UMAP space for each cell type. We then assigned each nucleus as either *FOXG1+* or *FOXG1-* based on an expression threshold of 4 logCPM, and performed Leiden clustering at low resolution (0.3). Any GD135/neonate astrocyte nucleus from the thalamic dissection that co-clustered with primarily telencephalic astrocytes was assigned striatum if it mapped to the telencephalic astrocyte ABCA subclass. Any GD135/neonate astrocyte from the striatum dissection in the same Leiden cluster was the thalamic nuclei was assigned thalamus if it was *FOXG1-* and it mapped to the non-telencephalic ABCA subclass. Any GD135/neonate astrocyte from the thalamic dissection that did not meet the first condition but was *FOXG1+* or mapped to the telencephalic ABCA subclass was assigned striatum. Any GD135/neonate GABAergic neurons from the striatum clustering with thalamic GABAergic neurons (either TRN or midbrain-derived) was assigned thalamus, any thalamic neuron clustering with medium spiny neurons was assigned striatum, and any *FOXG1+* nucleus from the thalamic dissection was assigned striatum. Any GD135/neonate neurons clustering with thalamic glutamatergic neurons were assigned thalamus, and any GD135 glutamatergic neurons not clustering with thalamic glutamatergic neurons were assigned PFC. Since OPCs did not cluster by region, we did not reassign region based on clustering, but instead, used only *FOXG1*: any OPCs from the thalamic dissection that were *FOXG1+* were assigned striatum. The resulting assigned region annotation is saved in the "region" .obs variable of the annotated data files, while the original region is saved in "region_dissected".

**Region reassignment of cross-contaminant nuclei in mouse.**

Cross-region contamination was higher in the E18.5 and P4 mouse timepoints, but present in all ages, primarily between striatum and thalamus (**Fig. S6**). Similarly to marmoset, we subsetted the cell type of interest from the cross-age, cross-region integrated cell x gene matrix, re-computed neighbors using the scVI latent space, and recomputed the UMAP space for each cell type. We then assigned each nucleus as either *Foxg1+* or *Foxg1-* based on an expression threshold of 4 logCPM, and performed Leiden clustering at low resolution (0.3). Nuclei in immature astrocyte clusters composed of mixed regions were reassigned based on *Foxg1* expression or assignment in ABCA subclass by MapMyCells. Astrocytes from the thalamic dissection in these mixed region clusters were reassigned as striatum if they were *Foxg1+* or had the subclass of "Astro-TE-NN". Immature clusters exhibiting more clear telencephalic and diencephalic divisions (mostly from P4) were manually reassigned, either from thalamus to striatum, or striatum to thalamus, to match the predominant region of origin for that cluster. All thalamic dissected astrocytes in the predominantly telencephalic and P4 cluster were manually reassigned to striatum. For the primarily thalamic and P4 cluster, we reassigned all cells to match their ABCA subclass (all "Astro-TE NN" being labeled striatum and all "Astro-NT NN" being labeled thalamus) and any cells that were *Foxg1+* were also labeled striatum. Because this cluster contained *Foxg1+* cells, we did not reassign all nuclei in it to thalamus. For the mature astrocytes, thalamic nuclei were reassigned to striatum if they clustered with telencephalic astrocytes and were assigned the "Astro-TE NN" ABCA subclass, and vice-versa for striatal astrocyte nuclei clustering with thalamic astrocyte nuclei. For OPCs, thalamic dissected cells were reassigned striatum if they were *Foxg1+*. For excitatory neurons, any striatum dissected cells were reassigned to cortex or thalamus based on the predominant region of the nuclei in their cluster . Any thalamic dissected excitatory neurons were reassigned as cortex if they clustered with cortical excitatory neurons and were *Foxg1+*. Finally, for inhibitory neurons, any thalamic dissected cells outside of the TRN cluster in E18.5 or P4 were reassigned either cortex or striatum based on the prominent region of the cluster they grouped with, and any cell at E18.5 or P4 in the excitatory TRN cluster were reassigned as thalamus. At all other time points, any thalamic excitatory neuron that was *Foxg1+* was reassigned as cortex. The resulting assigned region annotation is saved in the "region" .obs variable of the annotated data files, while the original region is saved in "region_dissected". 4.4% of astrocytes, 0.2% of OPCs, 2.6% of excitatory neurons, and 3.8% of inhibitory neurons were reassigned from their dissected region. There was also a batch of 90 weeks nuclei that was mislabeled prior to sequencing as striatum, but reassigned to cortex due to almost all cells aligning to cortical clusters within our data and to the ABCA clusters (see "Notes" column in **Table S1**).

**Calculation of rDEGs, aDEGs, and sDEGs using a pseudobulk method.**

Regional differentially expressed genes (rDEGs, **Fig. 2**) were calculated as previously described[7] with some modifications. rDEGs were calculated for each individual developmental time point, cell type, and species separately. To use the maximum number of genes available for each marmoset time point, pre-integrated adult or developmental gene counts (that is, with ~27,000 genes for adult marmoset data and ~31,000 genes for developmental and aged data) were used for rDEG calculation. We first created per-region metacells for each cell type (astrocyte, OPC, GABAergic neuron, and glutamatergic neuron) by averaging the raw counts of all cells of each cell type per region and per replicate. If a metacell of one region, cell-type, and replicate combination had fewer than 50 cells, it was omitted. Because they had few rDEGs between them, motor and prefrontal cortices were grouped for rDEG analysis. Metacell counts were normalized to 100,000 counts total and log10-transformed. We required rDEG candidates have at least 10 transcripts per 100,000 in at least one metacell per region and be expressed in at least 33% of nuclei in the metacell. 33% was chosen to require a significant portion of cells in a metacell to express the gene, but also be low enough to account for dropout[115]. Genes with $>10^{0.5}$ (3.16) fold-change (logFC) expression in the same cell type between two regions (pairwise) were considered rDEGs.

For marmoset, at the 30-month timepoint we restricted this analysis to the 2 30-month donors (bi005 and bi007, from our previous study[7]) that were represented in each regions. Because we averaged and normalized gene expression across cells within a region/age/species during metacell creation to calculate differential expression, we do not explicitly account for the relative contribution of each biological donor and/or replicate to differential expression. For this reason, for marmoset rDEGs, where the biological donors are balanced within each developmental time point (i.e., each animal donated each brain region), we filter rDEGs to those shared by at least 2 biological donors. For the mouse data, in which some regions were donated by different animals within a time point due to the aforementioned failures, and for age- and species- comparisons, which are inherently not replicate-balanced (no donor is represented in multiple ages or species), we do not enforce this criterion. Thus, for mouse rDEGs, aDEGs, and sDEGs, , we merged all biological replicates within a group into one metacell. As an alternative source of replicate specificity information, we calculated the fraction of replicates in each group that had at least 33% of cells expressing each gene.

To plot rDEG expression heatmaps as in **Figs. 2-3B**, we plotted the rDEG lists from fetal, early adolescent, and adult time points, ordered first by the age the gene was differentially expressed and second by the region(s) that the gene that was more highly expressed in, including repeats if an rDEG was detected at multiple time points. Marmoset rDEGs were only included if they were present in both replicates of a given time point, while mouse rDEGs did not have a replicate restriction.Because of the high heterogeneity in the mouse striatal astrocytes due to the multiple immature populations, these cells were reordered to place cells of similar populations together in the heatmap. To do so, striatal astrocytes were clustered at a low leiden resolution (0.1) and these clusters were ordered using a combination of the pseudotemporal ordering and the ages present in the cluster. A similar logic was applied to order marmoset striatal nuclei in the rDEG expression heatmaps.

Raster plots underneath the rDEG expression heatmaps were generated as separate figures using custom Python scripts with the help of ChatGPT 4.0. In brief, we created a matrix encoding the age and region(s) for which each gene was an rDEG, and plotted a line of the corresponding color in the corresponding raster row. These separate figures were manually aligned below the scanpy-generated rDEG expression heatmaps to the best of our abilities, but the x-axes may not be exactly aligned. To create the UpSet[116] plots in **Figs. 2, 3C-D**, **Fig. 5C-D, Fig. 6F,** and **Fig. S16B-C**, we used the package UpSetPlot (https://github.com/jnothman/UpSetPlot). The "active" dots were manually colored according to the group's color scheme for clarity.

The pairwise region rDEG scatter plots in **Fig. S7** were generated as follows. First, we recalculated rDEGs for each species within each region pair using the same metacell method described above, with one modification to allow more lowly expressed genes to pass the filter: requiring a gene be expressed by a minimum of 33% of cells in *any* region of either cortex, striatum, or thalamus. We then plotted the log fold-changes (calculated as described above) for each gene in (x,y) with x being the logFC value for one region pair (e.g. cortex-striatum), and y being the logFC value for another region pair (e.g. cortex-thalamus). The sign on the logFC value was determine with respect to the region shared across the pairwise comparisons (e.g. cortex). Pearson's correlation coefficient, r, was calculated using the scipy stats package (https://docs.scipy.org/doc/scipy/reference/generated/scipy.stats.pearsonr.html). A gene was marked as rDEG in one region if it had greater than 0.5 magnitude logFC in that region, or both if it had greater than 0.5 magnitude logFC in both.

aDEGs **(Fig. 5)** were calculated in a similar manner, except for that the metacell axis was age instead of region, and that no cross-replicate consistency was imposed for either species, because no donor or biological replicate provided tissue for multiple developmental time points. For aDEG and rDEG calculation in marmoset, lateral septal GABAergic neurons and putative hippocampal Cajal-Retzius neurons were removed, as we could not assign their region to either cortex, striatum, or thalamus, before metacell generation. Similarly for mouse, we omitted lateral septal GABAergic neurons and glutamatergic neurons from the anterior olfactory nucleus before aDEG and rDEG calculation. To calculate species differentially expressed genes (sDEGs, **Fig. 6**), we first combined downsampled mouse and marmoset datasets of all developmental time points and brain regions using the intersection of genes with 1:1 orthologs converted to mouse gene IDs using **Table S29**. If no match was found in the table, the marmoset gene name was converted to sentence case. We then created metacells of each supercluster for each species, where species was the metacell axis, and compared gene expression between species within each supercluster. As for aDEGs, we did not impose a cross-replicate requirement on sDEGs for either species because no replicate can be a member of both species.

To aid the interpretation of region-shared vs. region-specific aDEG profiles, we divided astrocyte aDEGs into 3 groups: group CA-RS (cell type agnostic, region specific) are astrocyte aDEGs that are also aDEG in neurons and OPCs for a given brain region; group AS-RA (astrocyte specific, region agnostic) are shared between astrocytes of all brain regions; and group AS-RS (astrocyte specific, region specific) are specific to astrocytes in a given brain region. CA-RS aDEGs are shared between all analyzed cell types in a given region and not shared with other regions. We found very few (3 or less) CA-RS aDEGs within marmoset GD135-14-month comparisons. Full lists of all mouse and marmoset aDEGs and associated UniProt GO annotations (as for rDEGs) for P4 vs. P90 and GD135 vs. 14-month comparisons respectively, including CA-RS, AS-RA, and AS-RS lists, are provided in **Tables S19-20.**

Group AS-RA aDEGs reflect universal aspects of astrocyte transcriptional identity at each developmental stage, regardless of brain region. We found 74 of these within marmoset GD135 vs. 14-month aDEGs and 56 within mouse P4 vs. P90 aDEGs. Group AS-RS aDEGs reflect the brain region's influence on the maturation of astrocytes *only* in a given brain region. In contrast to group CA-RS (above) they are unique to astrocytes (vs. OPCs and glutamatergic and/or GABAergic neurons) in a given brain region, and (unlike group AS-RA) are not shared with astrocytes in other brain regions. We found 20 of these in striatum, 51 in cortex, and 125 in thalamus within GD135 vs. 14-month aDEGs. Plotted genes (**Fig. 5F**, **Fig. S16D**) were manually selected to include genes that are both developmentally up- and down-regulated, and which have varied annotated functions that may be of interest to those studying astrocyte biology and/or neuron-astrocyte communication, but are not meant to suggest increased interest or importance compared to other cortical group AS-RS aDEGs.

We calculated the overlap between rDEGs and aDEGs between mouse and marmoset (**Fig. 6B-C**) by converting marmoset gene names to mouse gene names based on 1:1 orthologs using **Table S29** (as before, if no match was found in the table, the marmoset gene name was converted to sentence case) and taking the intersection between the two lists. To calculate whether a marmoset astrocyte rDEG was more likely to be a mouse astrocyte rDEG and vice-versa, we performed a Fisher's exact test using scipy stats's "fisher_exact" (https://docs.scipy.org/doc/scipy/reference/generated/scipy.stats.fisher_exact.html). 2x2 contingency tables were generated at each time point as follows: first row, first column was the number of overlapping rDEGs. First row, second column was the number of mouse rDEGs that were not marmoset rDEGs. Second row, first column was the number of marmoset rDEGs that were not mouse rDEGs. Second row, second column was the number of non-rDEG (in either species) overlapping 1:1 orthologs that men minimum expression criteria (at least 10 transcripts per 100,000 in at least one metacell per region and be expressed in at least 33% of nuclei in the metacell) to be considered as rDEGs. The test was run analogously for aDEGs. These tests are implemented in the Jupyter notebook "rDEGs_overlaps_xSpecies_current.ipynb" and "aDEGs_overlaps_xSpecies_current.ipyb".

**Automated UniProt annotation and querying of SFARI Gene 3.0.**

Code to query UniProt programmatically was modified from the companion document to the publicly available "Programmatic access to UniProt using Python" webinar at https://colab.research.google.com/drive/1i9UtVqa4m9WQ4ZVJkbGdwWbto_W7zmP_. Our scripts are available on our GitHub repository with the "mine_uniprot" prefix within rDEG, aDEG, and sDEG subfolders. Briefly, gene symbols from rDEG/aDEG/astrocyte sDEG lists and the appropriate species taxonomy ID were used to search for the UniProt accession number for each gene. The first result's primary accession number was then used to return the full protein name, GO Cellular Component, GO Biological Process, and GO Molecular Function for each protein. Multiple results were concatenated with commas and populated in a data frame for export with r/a/sDEG lists. Genes with no results were populated with "NaN" (empty) values. Genes beginning with the prefix "LOC" (unnamed) or "MT-" (mitochondrial) were ignored.

SFARI 3.0 genes from the 4/3/25 release were downloaded from the SFARI Human Gene Module (https://gene.sfari.org/database/human-gene/). Telencephalic and diencephalic sDEGs that were higher in marmoset (vs. mouse, hereafter marmoset sDEGs) were converted to human gene symbols based on 1:1 orthologs using **Table 29** (if no match was found in the table, the marmoset gene name was converted to all capital letters). For each set of marmoset sDEGs (diencephalic and telencephalic astrocytes), the following contingency table was constructed: number of SFARI genes that marmoset sDEGs in first row, first column; number of SFARI genes that were not marmoset sDEGs in first row, second column; number of non-SFARI marmoset sDEGs in second row, first column; and number of non-SFARI, non-sDEG genes (assuming 20,000 protein-coding genes in the human genome) in the second row, second column. A Fisher's exact test was run on this contingency table using Scipy stat's "fisher_exact" function in Python.

**Astrocyte subclustering within each brain region.**

We conducted sub-clustering for subsetted astrocytes from each brain region for each species separately (**Fig. S12-13**). For marmoset, using the subsetted cell x gene matrix for each region (all cell types), we removed small 10x Chromium batches with smaller than 500 cells (from reassigned regions), highly variable genes were recalculated (minimum number of batches equal to the floor of the total number of 10x Chromium batches divided by 10, to avoid batch-specific differentially expressed genes while including region-specific variation), the scVI model re-trained, and the neighborhood graph and UMAP coordinates re-calculated on subsetted astrocytes as described above. For mouse, highly variable genes were recalculated per region based on the original dissected region rather than the reassigned region due to a known issue with small batch sizes with the sc.pp.highly_variable_genes function. Our sub-clustering procedure was inspired by earlier methods[62]. Scanpy's "tl.leiden" function was used to identify clusters over a range of decreasing resolution parameters, purposefully starting with an intentionally high resolution that led to over-clustering. At each resolution, the minimum number of pairwise and one-versus-rest (where one group is compared to a metacell of all other groups combined) differentially expressed genes (DEGs) for each cluster and pair of clusters was calculated using the metacell method described above for rDEGs/aDEGs/sDEGs, with metacells calculated for each cluster, not incorporating replicate information. We first conducted a course resolution scan with increments of 0.05, followed by a fine-grained resolution scan in the target resolution range with increments of 0.01. We chose a clustering resolution that resulted in at least 3, but often 10 or more, pairwise and one-vs-rest DEGs for each subcluster and subcluster pair. Subclusters were named according to the following convention: [abbreviated_region]_Ast[subcluster number]_[known subtype name (immature, fibrous, and/or protoplasmic)]_[top marker genes]. More details are available in the scripts used to generate subclusters on our GitHub repository, with the prefix "subcluster_mouse_astrocytes" for mouse, or "integrate_xAge_subcluster" for marmoset.

**Pathway analysis with WebGestalt.**

We employed WebGestalt 2024[54,55] (https://www.webgestalt.org/) over-representation analysis (ORA) for pathway analysis. For the reference gene set, we used either all genes present in the final mouse counts matrix ("adata.var_names"), all genes present in the fetal, neonate, 7-month, 14-month, and aged marmoset data (generated in the current study with the mCalja1.2.Pat.X reference), or all genes present in the adult (29-32 month) marmoset data (generated previously[7] and aligned to cj1700). We used human as the host species for marmoset, mouse as the mouse species for mouse, the Gene Ontology (GO) Biological Process[117,118] noRedundant and KEGG[119] pathway functional databases, weighted set cover redundancy reduction for pathway display, and default advanced WebGestalt settings. Because differentially expressed genes were calculated in a pairwise manner, pathway analyses were performed on genes differentially expressed in both directions (i.e., either up- or down-regulated, as opposed to unidirectionally). Lollipop plots were generated from "Description", "Ratio", and "FDR" columns of the weighted set cover redundancy-reduced pathway table using custom Python scripts with the assistance of ChatGPT 4.0.

**Cross-species integration with scANVI[73].**

scANVI[73] is a semi-supervised variational autoencoder variant of the previously described scVI model that utilizes cell type label information in its latent space. We used a random downsample of 20,000 cells from any age or region for each species. Next, we added a less granular “supercluster” annotation to each cell based on its leiden cluster annotation. For example, two marmoset cowrtical excitatory neuron L6IT clusters were combined into one supercluster. In some cases, the leiden cluster to supercluster mapping was one:to:one, as for OPCs, lateral septal inhibitory neurons, diencephalic and telencephalic astrocytes, and others. To create the embedding space, we used a small subset of highly variable genes for cross-species integration that was inspired by an earlier approach[62]. Briefly, we converted the marmoset gene names to their mouse orthologs and subsetted both datasets to the intersection of shared genes. Next, we took the top 50 most highly expressed genes in each supercluster (as determined by a Wilcoxon rank-sum test in scanpy) in each species separately, then used the intersection of those lists as the highly variable genes entered into the model. We created a scANVI model using the superclusters as the labels key, the dispersion set to “gene-batch”, and calculated on the raw counts layer. Finally, we calculated the neighborhood graph and UMAP using scanpy’s previously mentioned functions.

**Cross-species integration with SATURN[74].**

SATURN is a deep learning method used to integrate cells from multiple species in the same low-dimensional space. It utilizes the ESM2[120] language model and reference genomes from Ensembl (for marmoset: https://useast.ensembl.org/Callithrix_jacchus/Info/Index, https://useast.ensembl.org/Mus_musculus/Info/Index for mouse, as in the original publication) to generate protein embeddings that predict similarity of genes across species. The marmoset protein embedding space was generated by the lead author of the SATURN study and is linked at https://github.com/snap-stanford/SATURN/issues/19, and the mouse protein embedding is available at http://snap.stanford.edu/saturn/data/protein_embeddings.tar.gz. SATURN uses a combination of cell type annotations from the user and the protein embeddings to create “macrogenes”, groups of genes that are predicted to be “functionally related” and “coexpressed across species”[74]. It then uses a weakly supervised autoencoder to refine the macrogene space by using a triplet loss function that incorporates within-species cell type annotations. Before running SATURN, we first randomly downsampled the annotated cell x gene matrices for both species to 100,000 nuclei total to reduce computational burden. We provided SATURN with a mapping between mouse and marmoset supercluster names, which were identical except for 2-3 unique superclusters per species. We trained the SATURN model as instructed in the tutorial (https://github.com/snap-stanford/SATURN/blob/main/Vignettes/frog_zebrafish_embryogenesis/Train%20SATURN.ipynb), using raw (not normalized) counts, 2,000 macrogenes, 8,000 highly-variable genes, and our supercluster mapping as the cell type mapping file. The resulting SATURN embedding in UMAP space was used to generate the plots in **Fig. S17**.

We performed SATURN cross-species integration on the same downsampled, supercluster-annotated cell x gene expression matrix as for our scANVI approach (with the exception of L5IT cortical excitatory neurons labeled separately for marmoset). By visual inspection, the species-integrated UMAP calculated from the SATURN embedding yielded similar results as our scANVI approach: broad conservation with a few species-specific clusters as described above (**Fig. S17C**). However, because SATURN does not explicitly rely on cell type annotation to group cells from different species, it was able to merge cortical glutamatergic L5IT supercluster mouse neurons with the corresponding population in marmoset (**Fig. S17C**, dark green cluster with the asterisked label), despite the fact that L5IT mouse neurons were not separately annotated as such, but grouped with L4/5IT cortical glutamatergic neurons. The scANVI approach did not result in this merging unless we grouped L5IT and L4/5IT neuron superclusters in both species, which is the approach we adopted for the results shown in **Fig. 6**. However, the relative position of superclusters in the SATURN UMAP seems to be less meaningful than for the scVI/scANVI embeddings: related neuronal clusters are no longer adjacent in UMAP space, OPCs and MOLs are far away from one another, and microglia were well integrated, despite species differences apparent in the scANVI integration and past studies suggesting they should be species-divergent[77,121]. Closer examination of the SATURN-integrated astrocytes illustrates concordant results with the scANVI integration: telencephalic marmoset and mouse astrocytes were well-integrated, diencephalic astrocytes were more separated, and immature astrocytes were almost completely separated (**Fig. S17D**).

**Cell-cell communication analysis.**

We performed cell-cell communication analysis on a per-region (prefrontal and motor cortex pooled), per-age, per-species basis in Python using CellPhoneDB[65] via Liana[122], a scanpy-friendly package that integrates several cell-cell interaction inference methods. To avoid spurious findings due to differences in cluster proportion (i.e., bias towards or away from rare cell types), we first randomly downsampled all clusters to a maximum of 1,000 nuclei and dropped all clusters with fewer than 100 (thalamus and cortex) or 70 (striatum, to avoid dropping the sparse *CHAT+* neuron cluster) nuclei. As recommended in the Liana documentation, we ran CellPhone DB on the log1p-transformed normalized counts matrix, grouping by leiden cluster, using the "consensus" resource for marmoset and the "mouse consensus" resource for mouse, and requiring that 33% of nuclei in a cluster express a gene at nonzero levels for it to be considered for ligand-receptor enrichment. CellPhoneDB generated two outputs of interest: 1) "lr_means", a measure of interaction magnitude, and is simply the mean expression of the ligand in the source cluster averaged with the mean expression of the receptor in the target cluster, and 2) the permutation-based p-value, a measure of interaction specificity. Briefly, the specificity p-value for a ligand-receptor pair is calculated as the proportion of null distribution (generated by randomly permuting the cluster labels of all cells) means that are greater than or equal to the actual mean expression calculated. For all ligand-receptor pairs shown, we required a CellPhoneDB p-value of less than $10^{-6}$. Because many ligands and receptors are expressed by both neurons and astrocytes, we ran CellPhoneDB on only the neuronal and astrocytic clusters to increase specificity by removing cell types that could deflate the specificity p-value. To generate the dotplots shown in **Fig. 4A-B** and **Fig. S14A-B**, we restricted the plot to only those ligand-receptor (L-R) pairs that were near-unique (p-value below $10^{-6}$) to 2 or fewer (except for mouse P90, which was relaxed to 3 or fewer) neuronal clusters and the dominant astrocyte subtype (or vice-versa). Otherwise, the top 25 L-R pairs shown are mostly shared across all neuronal subtypes.

For the upset plots shown in **Fig. 4C-D** and **Fig. S14C-D**, non-filtered (non near-unique) L-R pairs were used. Because immature versions of the most abundant neuronal and astrocytic cluster dominated at early developmental time points in both species, L-R pairs between these immature clusters were used at these time points only, and only if their abundance was greater than the corresponding mature cluster. For example, in neonate mouse, we used L-R pairs between the 'Neuron_Excit_Ctx_Immature_L23IT_Ptprk' Leiden cluster and the 'Astrocyte_Immature_Ptprz1' Leiden cluster, while in adult mouse, we used L-R pairs between 'Neuron_Excit_Ctx_L23IT_Cam2ka' and 'Astrocyte_Telen_Mature_Slc1a2'. Details are available in the Jupyter notebooks in our GitHub repository (https://github.com/Feng-Lab-MIT/AstrocyteHeterogeneity).

We ran three analyses at a single developmental time point (14-month marmoset) to check whether differences in neuron-astrocyte and astrocyte-neuron L-R pair expression across regions were driven solely by neuronal expression differences across regions. We first ran a region-scrambled CellPhoneDB analysis that included local neuronal and astrocytic populations from all regions in the same run of CellPhoneDB (as opposed to running CellPhoneDB separately by region as in the manuscript **Fig. 4** and **Fig. S14**). See "CCC_check_specificity_toneuron_astrocyte_shuffle_nofilter_allnas_current.ipynb" in our GitHub repository for more details. We compared the magnitude and specificity of N-A L-R pairs across regions by examining L-R pairs between local interneuron clusters (cortical MGE-derived *PVALB+* interneurons, striatal MGE-derived interneurons, and thalamic midbrain-derived interneurons) and local astrocyte populations (cortical telencephalic astrocytes, striatal telencephalic astrocytes, and thalamic diencephalic astrocytes). Our results showed that neuron-astrocyte L-R enrichment patterns are not the same across all astrocyte subtypes for a given local neuronal subtype. In a second analysis, we compared the current astrocyte-neuron/neuron-astrocyte L-R pairs, and their sharing/divergence across regions, to OPC-neuron/neuron-OPC L-R pairs. See "CCC_check_specificity_toneuron_opc_shuffle_nofilter_allnos_current.ipynb" in our GitHub repository for more details. We found that only 40-60% of L-R pairs overlapped between astrocytes and OPCs within a given region. Finally, we quantified the proportion of genes in L-R pairs of a given neuron-astrocyte pairing that are also DEGs in that neuron cluster, or astrocyte regional subtype, relative to others. Our analysis showed that N-A L-R pairs contained DEGs from both neurons and astrocytes, with the proportion of ligands or receptors that were also cluster DEGs ranging from ~20-70%. In fact, astrocytes had a much larger proportion of overlap between ligand (when source) or receptor (when target) and leiden-level cluster DEGs. However, the increased astrocyte DEG presence in L-R pairs should be interpreted with caution, because there are many more neuronal than astrocytic Leiden clusters, potentially leading to more general "astrocytic" genes being DEGs, while neuronal DEGs are more leiden cluster-specific.

**Pseudotime analysis and calculation of gene change scores.**
We performed pseudotime analysis using Palantir[68], a scanpy-friendly package that orders cells along pseudo-temporal trajectories based on diffusion space and assigns each cell a probability of differentiating into each user-defined terminal state based on a Markov chain. Importantly, unlike single-cell velocity based methods that rely on estimates of spliced and unspliced counts[123–125], Palantir runs diffusion maps in a latent space calculated from the regular counts matrix, which is in our case the scVI latent space (pca_key= "X_scVI", n_components=5). The "determine_multiscale_space" parameter n_eigs was set to 5 to avoid a documented error (https://github.com/dpeerlab/Palantir/issues/84). Palantir imputes missing gene expression data in log1p-transformed counts per million space using MAGIC[126], which is useful for visualizing gene expression. The root and terminal cells were manually specified for each cell type (oligodendrocyte lineage or astrocyte) based on their location in UMAP space combined with cluster and age information, and 500 waypoints were used. We calculated pseudotime trajectories for the astrocyte and oligodendrocyte lineages on a per-species region-combined basis. To reduce computational load, marmoset oligodendrocytes were randomly downsampled to 100,000 nuclei. We first ran Palantir pseudotime analysis on the oligodendrocyte lineage as a "sense check" for the algorithm, as the biological ground truth of oligodendrocyte lineage differentiation is well known (OPC→COP→NFOL→MFOL→MOL[69]), and found that Palantir's calculated pseudotime was able to recapitulate the known differentiation trajectory.

We then used Mellon[70] to identify genes whose expression changes significantly in pseudotime transition periods, identified as regions of low cell-state density, for each pseudotime trajectory branch (AST-DI and AST-TE). We followed the basic Mellon tutorial "Density Estimator for scRNA-seq data" (https://mellon.readthedocs.io/en/latest/notebooks/basic_tutorial.html) and the "Gene change analysis" tutorial (https://mellon.readthedocs.io/en/latest/notebooks/gene_change_analysis_tutorial.html). Mellon, a companion algorithm to Palantir, estimates cell-state "densities", that is, density of cells in each state, from high-dimensional single cell data using a Bayesian model. It also computes "local variability" on a per-cell, per-gene basis as the maximum (across nearest neighbors) normalized (by the distance between cells in state space) difference in MAGIC-imputed expression of a gene in a cell compared to that of its nearest neighbors. Gene change scores are then calculated using a specified set of cells (e.g.., a pseudotime trajectory branch) as the inverse density-weighted (such that lower densities have higher weights) average local variability for a gene across all cells. As stated in the source paper, "gene-change analysis quantifies the influence of a gene in driving state transitions in low-density regions". Briefly, we subsetted the integrated all-region, all-age data to astrocytes, calculated the diffusion maps, calculated densities, calculated pseudotime using Palantir (as described above), computed local variability, and calculated gene change scores separately for each pseudotime branch (AST-TE and AST-DI). We then computed gene expression trends from the MAGIC-imputed expression data and visualized the trends over pseudotime for the top 25 highest change-scoring genes (**Fig. S15C**). We also plotted the average trend overlaid on the per-cell trend scatterplot vs. pseudotime for each of these 25 genes, and selected 1 example gene from each trajectory in **Fig. S15D**. These plots are available in our GitHub repository for marmoset in the "marmoset_mellon_gene_change_allages_regcombined_current.ipynb" notebook and for mouse in the "mouse_mellon_gene_change_allages_regcombined_current.ipynb" notebook. A list of the highest change-scoring genes is provided for both species in **Table S18.** We gratefully acknowledge the assistance of Dominik Otto on this analysis (https://github.com/settylab/Mellon/issues/13).

**RNA fluorescence in situ hybridization (FISH).**
Neonate (P4) or adult mice were deeply anesthetized via hypothermia (neonates) or isoflurane overdose (adults) and rapidly decapitated. Brains were extracted and frozen in OCT compound (Tissue-Tek) over dry ice and stored at -70°C. Briefly, neonate and adult marmosets were deeply sedated by intramuscular injection of ketamine (20–40

mg/kg) or alfaxalone (5–10 mg/kg), followed by intravenous injection of sodium pentobarbital (10–30 mg/kg). When the pedal with-drawal reflex was eliminated and/or the respiratory rate was diminished, animals were trans-cardially perfused with ice-cold sterile PBS. Whole brains were rapidly extracted into fresh PBS on ice for transfer to the lab (<30mins), then frozen in OCT over dry ice and stored at -70°C. Brains were sectioned sagittally on a cryostat (Leica) at -16um with a cutting temperature between -15 and -17°C and mounted on SuperFrost Plus (VWR/EMS) slides and stored at -70°C.

Multiplexed smFISH was performed using the RNAscope™ HiPlex kit and protocol (Advanced Cell Diagnostics). Briefly, sections were removed from 70°C, placed directly in 4% PFA, and fixed for 1 hour at room temp. Next, sections were dehydrated via an ethanol series of 50%, 70%, 100%, and 100% EtOH in water, each immersion for 5 minutes at room temp. Samples were then treated with protease (Protease Plus for neonate mouse, Protease III for neonate marmoset, adult marmoset, and adult mouse) for 30 minutes at room temperature. Probes (at a 1:50 dilution factor) were hybridized for 2 hours at 40°C. Following the application of amplifiers and fluorophores at 40°C, adult marmoset sections were incubated with TrueBlack Plus (Biotium) at a 1:30 dilution factor (1.5x) for 10 minutes. Sections were counterstained with DAPI and coverslips were affixed with ProLong Diamond Antifade Mountant (Thermo Fisher). Following each round of imaging, coverslips were removed by soaking slides in 4x SSC, fluorophores were cleaved, and the next set of tails were applied, followed by additional TrueBlack (for adult marmoset sections) and DAPI application (every other round for mouse, every round for marmoset) prior to mounting and acquiring images for the next round. The tail application and imaging order was as follows: T1-3, T4-6, T7-9, and T10-12 for mouse; T7-9, T4-6, T10-12, and T1-3 for marmoset. We changed the round order for marmoset after observing low signal-to-noise for T7-9 when imaged in the third round. Imaging control probes (housekeeping genes *POLR2A*, *PPIB*, and *UBC*) also showed significant loss of signal in the third round of imaging in adult marmoset brain.

Single-round FISH for marmoset rDEGs and sDEGs in both species was performed using the RNAscope™ Multiplex Fluorescent V2 kit and protocol (Advanced Cell Diagnostics. Briefly, sections were removed from 70°C, placed directly in 4% PFA, and fixed for 1 hour at room temp. Next, sections were dehydrated via an ethanol series of 50%, 70%, 100%, and 100% EtOH in water, each immersion for 5 minutes at room temp. Samples were then treated with Protease Plus for 30 minutes at room temperature. Probes (mixed at 1:1:50 ratios of C2, C3, and C1 respectively) were hybridized for 2 hours at 40°C. Following the application of amplifiers (30 min for Amp1, 30 min for Amp2, and 15 min for Amp3), HRP-C1 (TSA fluorescein), HRP-C2 (TSA Cyanine 5), and HRP-C3 (Cyanine 3) signals were developed. Adult marmoset sections with incubated with TrueBlack Plus (Biotium) at 1:30 dilution factor (1.5x) for 10 minutes. Sections were counterstained with DAPI and coverslips were affixed with ProLong Diamond Antifade Mountant (Thermo Fisher).

Images were acquired using an Olympus Fluoview FV3000 confocal microscope using the multi-area time-lapse (MATL) module to record stage positions for re-use in subsequent imaging rounds. Fluorophores were excited with 405nm, 488nm, 561nm, and 640nm lasers. Imaging settings were adjusted on a per-experiment (and in rare cases, per-sample) basis due to batch-level variations in signal intensity so as to maximize the dynamic range of pixel intensities. All brain regions within a slide were imaged with the same laser power and voltage settings. Fields of view consisting 3x3 grids or smaller (for smaller regions in neonate tissue) with the 20x magnification objective lens were obtained in each brain region of interest (PFC, striatum, and thalamus). We obtained a z-stack covering the entire axial extent of the tissue section with a z-step size of 2μm. Raw images from the microscope were converted to .tif format, flattened using a maximum intensity projection, and separated into individual channels. For HiPlex images, round 2-4+ images were registered to round 1 using the DAPI channel and cropped to mutually overlapping area using the HiPlex Image Registration Software v2.1 (ACD, https://acdbio.com/rnascope%E2%84%A2-hiplex-image-registration-software-v21), or, in cases where v2.1 failed to register most of the field of view, HiPlex Image Registration software v1.0.0 (ACD, provided by their technical support team).

RNA quality was assessed using positive control probes targeting housekeeping genes for each species. These included *POLR2A*, *PPIB*, and *UBC* for marmoset and *Polr2a*, *PPIB*, *Ubc*, *Hprt*, *Actb*, *Tubb3*, *Bin1*, *Ldha*, *Gapdh*,

*Pgk1*, *Bhlhe22*, and *Cplx2* for mouse (RNAscope HiPlex12 Positive Control Probe - Mm). For HiPlex datasets, control experiments were always performed simultaneously with the experiments for rDEG probes of interest, using the same reagents and brain slices of minimal stereotaxic distance to the slice used for rDEG probes. Control probe signal was not quantified, but the brightness and density of probes that are expected to be expressed ubiquitously in most/all cells were manually observed to judge RNA quality for the inclusion or exclusion of experiments. Control probes were omitted for RNAscope v2 experiments, as sufficient RNA quality in neighboring slides with tissue from the same animals had already been confirmed.

**FISH quantification with CellProfiler.**
Quantification of RNAscope HiPlex FISH images was preregistered during or after data collection on the Open Science Framework (https://osf.io/crs7v/; https://doi.org/10.17605/OSF.IO/6KUB2) and conducted with the analyzer blinded to rDEG identity, for mouse (for all genes except Sparc, for which we modified the analysis pipeline post-hoc, see Supplementary Note 1). The following text has been adapted from the preregistration, with major deviations from the preregistration being noted here. CellProfiler 4.2.5 was used to quantify RNA signal in each nuclei of different regions. Variations between experiments such as tissue quality or probe freshness caused variations in fluorescence, so it was necessary to use separate CellProfiler pipelines for each experiment for mouse (with the exception of one adult mouse experiment that needed to be split by replicate) and neonate marmoset to account for those differences. The adult marmoset tissue had more dramatic difference in quality between replicates, requiring a separate pipeline for each replicate. Crucially, the CellProfiler pipelines were always identical within each slice of tissue, allowing for each replicate to have unbiased comparison between regions.

We first created binary masks to eliminate large artifacts (e.g., very large/bright debris), large tears in the tissue, areas of very high autofluorescence, out-of-focus areas, poorly registered areas, and/or nearby brain regions that were not the region of interest. In most cases, the cropped region of interest was greater than ~80% of the original. For adult marmoset tissue, which exhibited persistent lipofuscin autofluorescence in later rounds despite quenching with TrueBlack Plus, we automatically generated masks to eliminate this punctate fluorescence as follows: minimum intensity projection of all three channels in the third round, gaussian filter with sigma = 2, intensity thresholding at 1.5 standard deviations above the mean, and median filtering with radius of 2 pixels. We then combined this autofluorescence mask with the large artifact mask mentioned above. This quality control step was not mentioned in our preregistration, largely because we did not anticipate the extent of remaining autofluorescence. The registered, cropped, masked max-projected images were processed through the CellProfiler quantification pipeline.

In CellProfiler, all images were scaled to stretch to the full intensity range and masked to regions of interest. The lipofuscin mask was used to filter nuclei covered more than 20% by lipofuscin and crop any smaller areas within remaining nuclei. Some regions had a higher background intensity than others, so the images were binarized to eliminate background noise. Binarization was achieved using the CellProfiler "Threshold" module, with the threshold set manually after using the Otsu algorithm to suggest options for values that would separate foreground and background. The manual threshold was used instead of Otsu, so that the definition of foreground and background would be consistent between regions. The exception to this was *Slc1a3*, our astrocyte marker, because its signal was known to be different between regions. Using this binarized signal, we were able to quantify the fluorescence from RNA in each nucleus. There were two rDEG probes in the adult mouse that required additional processing steps prior to binarization: *Sparc* and *Clmn* were particularly noisy in the thalamus specifically, with punctate signal outside of nuclei. Those genes required the "EnhanceOrSuppressFeatures" to suppress speckles. We also applied the "Smooth" module to *Sparc* in all adult mouse regions to reduce noise. From the binarized signal, we calculated mean intensity, or the fraction of the nucleus that is covered by the probe signal, and integrated intensity, the number of thresholded pixels within the area of the nucleus.

All nucleus objects identified by DAPI signal were expanded by 3 pixels to reflect RNA signal likely also existing outside of the nucleus, with the exception of P4 mouse nuclei, as they were more tightly packed together. To identify astrocytes, we first filtered by *SLC1A3/Slc1a3* mean intensity, using 5% for neonate mouse astrocytes, 6.5% for adult mouse astrocytes, 12.5% for neonate marmoset astrocytes, and 8% for adult marmoset astrocytes. This value was determined to reflect expected expression levels and similar fraction of astrocytes out of total cells based on the snRNAseq data. Next, we filtered out the astrocytes that had at least 50% mean intensity for neuronal markers *GAD2/Gad2*, *SLC17A6/Slc17a6*, and *SLC17A7/Slc17a7*. We did not filter based on *OLIG2/Olig2* because a subset of astrocytes are known to express *OLIG2/Olig2*[127]. We then measured the intensity of the binarized signal in our rDEG probes inside astrocytes and in all nuclei. To count a cell positive for a probe, the cell needed to either have at least 0.03 mean intensity or 6 integrated intensity. Both measurements were used to account for variations in cell size, as smaller cells would likely have lower integrated intensity and larger cells would likely have lower mean intensity. To compare rDEG expression between regions, we used the fraction of cells counted positive for a probe and the average mean intensity of all cells within the region. For mouse, these values were averaged within slices of the same replicate and then treated as a single data point.

We used DAPI signal, positive control probes, and cell type marker probe signals to qualitatively assess RNA quality (e.g., degraded or not degraded based on the brightness of fluorescence intensity) before running images through the analysis pipeline. If we observed unacceptably low tissue (e.g. over-digested, damaged, or folded slices or extremely high background) or RNA quality (little to no signal) for a given sample, we collected more images from a different slice or animal to maintain sample size prior to starting analysis. Given the low number of samples per region/age/species, we did not remove outliers, as we were not adequately powered to detect outliers post-hoc. Of note, changes were made to marmoset analysis pipelines after unblinding due to unrealistic detection of *FOXG1* expression in the thalamus. This gene is known to be a telencephalic patterning factor[128], a finding confirmed *in situ* by other groups (see for example the RIKEN Marmoset Gene Atlas[129,130] at https://gene-atlas.brainminds.jp/gene-image/?gene=370-6), triggering reevaluation of the lipofuscin masks and *FOXG1*'s binarization threshold. Ultimately, we excluded *FOXG1* from our quantification results because no version of our pipeline could overcome the large variations in signal-to-noise between brain regions. Another notable change to the pipeline after unblinding was an increase of the intensity threshold for binarizing *Sparc* images in neonate mouse,, as the minimal expression of the gene at the neonate timepoint led to an erroneously low intensity threshold under blinding. The *Sparc* mouse neonate analysis was also rerun with altered masks to remove the pial and ventricle surfaces of the PFC and striatum (see **Supplementary Note 1**).

For the RNAscope v2 data, the images did not require advanced masking to account for lipofuscin and were only masked for the regions of interest. Because there was no labeling for other cell type markers to filter the cells, the threshold of mean *SLC1A3* intensity for astrocytes was increased to 15% and the fraction of astrocytes per region was comparable to the HiPlex quantification. Because of differing signal-to-noise ratios of the RNAscope protocols, the threshold for a positive cell was increased to at least 5% mean intensity or 10 integrated intensity.

Because we did not have sufficient sample size to test for statistical significance in marmoset, we report only observations and trends. For mouse data, we used 2 univariate ANOVAs with Benjamini–Hochberg p-value correction for each measure presented: fraction of astrocytes positive for a probe and mean fraction of the astrocyte covered by the probe (i.e., mean intensity). If the effect of brain region was significant overall, Tukey's multiple comparisons (i.e., the Tukey HSD method) test was performed separately on each variable to test for the significance of pairwise differences between brain regions. Statistical analyses were performed in GraphPad Prism. Contrary to our preregistration, we did not first use a one-way repeated measures *multivariate* ANOVA, as we could not find a package to do so, but we believe this divergence to be minor given that we only report two outcome variables.

**Quantification of mouse astrocyte rDEG expression in the ABCA Whole Mouse Brain atlas.**
We utilized the Allen Brain Cell Atlas MERFISH spatial transcriptomics dataset[1] available at https://knowledge.brain-map.org/abcatlas and using the tutorial posted at https://alleninstitute.github.io/abc_atlas_access/descriptions/MERFISH-C57BL6J-638850.html. Next, we subsetted the data to astrocytes only by subclass, only '318 Astro-NT NN' or '319 Astro-TE NN'. To subset the data for the regions of interest, we selected striatum, thalamus, hypothalamus, midbrain, and cerebellum cells by the parcellation divisions 'STR', 'TH', 'HY', 'MB', or 'CB' respectively.

We selected PFC cells by parcellation structure being 'AId', 'PL', 'ORBl', 'AIp', 'ORBvl', 'ORBm', 'AIv', 'ILA', or 'FRP'. We selected motor cortex cells by parcellation structure 'MOp', or 'MOs.' Finally, we selected somatosensory cortex cells by parcellation structures of 'SS-bfd', 'SSp-ll', 'SSp-m', 'SSp-n', 'SSp-tr', 'SSp-ul', 'SSp-un',' or 'SSs.' We then averaged the gene expression for each gene of interest per region and plotted a heatmap of the resulting values.

**Mouse astrocyte labeling and ExR for mouse tissue.**
We stereotaxically injected 10 week-old adult C57 Bl/6J mice with AAV2/5 CAG-flex-GFP-4x6T (Addgene 196418) and AAV2/5 gfaABC1D-Cre-4x6T (Addgene 196410) as in ref[131] in prefrontal cortex, striatum, and thalamus (**Fig. 7A**). After 3 weeks of viral expression and pre-expansion staining for GFP, astrocytes were clearly labeled in our regions of interest (**Fig. 7B**). We found that a single expansion step, yielding an expansion factor of ~3.5x, was sufficient to visualize complex astrocyte morphology in gray matter regions of the mouse PFC, striatum, and thalamus (**Fig. 7C**, **Movies S1-18**). For reference, the effective resolution of ~4x expansion microscopy is ~70nm[132]. Though not the super-resolution afforded by other techniques such as electron microscopy, ~4x ExR is advantaged by rapid sample preparation, compatibility with conventional antibody staining, and rapid imaging of large volumes on a confocal microscope.

All AAVs in this study were packaged in-house as previously described[133]. Briefly, for each 150mm culture dish of HEK293 cells, 5.7µg of construct DNA was transfected with 22.8µg of capsid plasmid and 11.45µg of pADDeltaF6 using polyethylenimine 25K MW (Polysciences, 23966-1). Collection of cells and media for AAV harvesting began 72 hours after transfection, followed by iodixanol gradient ultracentrifugation purification using a Type 70 Ti Fixed-Angle Titanium Rotor (Beckman-Coulter, 337922). Titer was calculated using droplet digital PCR (ddPCR) as described by Addgene (https://www.addgene.org/protocols/aav-ddpcr-titration/)[134] using the QX200 AutoDG Droplet Digital PCR System (BioRad, 1864100).

For stereotactic intracranial injection of AAVs, C57BL/6J mice (3 females, 5 males) aged 10 weeks old were anesthetized with isoflurane (5% induction, 1.5% maintenance) and kept on a heating pad. Depth of anesthesia was confirmed via breathing rate and bilateral toe pinch. Mice were given subcutaneous sustained-release buprenorphine (1mg/kg) on the day of surgery and subcutaneous meloxicam (5mg/kg) on both the day of surgery and 24 hours after. Viruses were delivered using a pulled glass micropipette. A total volume of 500nL of AAV2/5 CAG-flex-GFP-4x6T and AAV2/5 GfaABC1D-Cre-4x6T[131], each at final a titer of 1 x $10^{12}$ vg/mL, were co-injected at 150 nL/min to the PFC (2.68 AP, 0.75 ML, 2 DV), striatum (1 AP, 1.7 ML, 3.5 DV), and thalamus (-1.46 AP, 1 ML, 3.5 DV). Mice were monitored for 3 days post-surgery to ensure proper recovery.

3 weeks following surgery for adult C57 Bl/6J mice, animals were perfused for ExR as described[37]. Briefly, animals were deeply anesthetized using isoflurane and transcardially perfused with ice-cold 2% acrylamide in PBS followed by ice-cold 30% acrylamide and 4% paraformaldehyde in PBS (by initial volume: e.g., for two mice, we dissolved 15g of acrylamide in 38.75mL deionized water, added 5mL 10x PBS, and 6.25mL 32% PFA). Brains were post-fixed in the same fixative solution overnight, transferred to 100mM glycine for 6 hours, and stored in PBS at 4°C until sectioning. Brains were sectioned coronally at 150µm on a vibrating microtome (Leica) and stained for GFP (primary antibody, Abcam chicken-anti-GFP at 1:1000, secondary antibody, AcX-conjugated goat-anti-chicken Alexa Fluor 488 at 1:200, see **Table S30** for antibody product information). Briefly, sections were permeabilized in 1x PBS + 0.05 Triton X-100 solution for 10 minutes at RT, blocked for 2 hours at RT in blocking buffer (5% normal goat serum (NGS) + 0.5% Triton X-100 in 1x PBS), incubated with primary antibodies in carrier solution (5% NGS + 0.25% Triton X-100 in 1xPBS) for 12-24 hours at 4°C, washed in PBST (1x PBS + 0.1% Triton X-100) 3 times for 10 min each at RT, incubated with secondary antibodies in carrier solution for 12-24 hours at 4°C, and washed in PBST 3 times for 10 min each at RT. Slices containing regions of interest (PFC, striatum, and thalamus) were identified using the online Allen Brain Institute adult mouse reference atlas with coronal sections.

One hemisphere containing each region of interest (the hemisphere that was injected, from the slice with the brightest GFP signal) was expanded at ~3.5x expansion followed by staining for GFP, Lectin, and GFAP for morphology characterization. For 2 of the 8 mice, another hemisphere from a neighboring slice was expanded ~18x for super-resolution imaging of astrocytic processes with synaptic proteins and rDEGs. Gels were generated using the ExR protocol[37]. Briefly, tissues were incubated in the first gelling solution for 30 min at 4°C followed by 37°C for 30 min to 2 h. To preserve blood vessel morphology, gels were treated with 0.5 kU/mL Collagenase VII overnight at 37°C, a variation on our lab's previous protocol[135] (**Fig. S20A**). After collagenase treatment, tissue-embedded gels were incubated in ExR denaturation buffer for 1h at 95°C. Denatured gels were fully expanded in deionized water by washing 2-4 times for 15-45 min each. ~18x-expanded gels were generated without collagenase treatment[37]. We omitted collagenase treatment for the ExR samples for two reasons. First, the goal of this experiment was not to capture astrocyte morphology, which might be locally disrupted by blood vessel breakage (indeed, most of the fields of view we imaged contained no blood vessels). Nevertheless, despite some broken blood vessels, astrocyte morphology appeared largely continuous in non-collagenase treated samples, except specifically at astrocyte contact sites with blood vessels (**Fig. S20B**). Second, collagenases could in principle be contaminated with other proteases, which might degrade sensitive epitopes and reduce signal captured by ExR.

For the ~3.5x-expanded gels for morphology analysis: After full expansion, we shrunk the gels for easier handling by incubating in 10x PBS for 10-30min. We then transferred gels to blocking buffer (5% NGS and 0.5% Triton X-100 in 1x PBS) and incubated for 90 mins - 2 hours at room temperature. Primary antibodies (chicken-anti-GFP and mouse-anti-GFAP, see **Table S30** for antibody product information) were incubated at 4°C for 16-24 hours in antibody carrier solution (5% normal donkey serum (NDS) and 0.25% Triton X-100 in 1x PBS) at a dilution factor of 1:200. Gels were washed in 0.01% Triton X-100 in 1x PBS 6 times for 15 minutes each at room temperature, and then incubated with secondary antibodies (donkey-anti-goat AF488, donkey-anti-chicken AF488, donkey-anti-mouse AF555, and Lycopersicon Esculentum (Tomato) Lectin (LEL, TL), DyLight 649, see **Table S30** for antibody product information) were incubated at 4°C for 16-24 hours in antibody carrier solution (5% normal NDS and 0.25% Triton X-100 in 1x PBS) at a dilution factor of 1:200. Gels were washed in 0.05x PBST (e.g., 500uL Triton X-100 in 1xPBS, 25mL of 1x PBS, up to 500mL of DIW) 6 times for 15 minutes each at room temperature to expand the gels for imaging.

For the ~18x-expanded gels for rDEG protein expression analysis: After full expansion, we shrunk the gels for easier handling by incubating in 10x PBS for 10-30min. We then transferred gels to blocking buffer (5% NGS and 0.5% Triton X-100 in 1x PBS) and incubated for 90 mins - 2 hours at room temperature. Primary antibodies (chicken-anti-GFP and mouse-anti-GFAP, see **Table S30** for antibody product information) were incubated at 4°C for 12-24 hours in antibody carrier solution (5% NDS and 0.25% Triton X-100 in 1x PBS) at a dilution factor of 1:50 to 1:200. Gels were washed in 0.01% Triton X-100 in 1x PBS 6 times for 15 minutes each at room temperature, and then incubated with secondary antibodies (donkey-anti-goat AF488, donkey-anti-chicken AF488, donkey-anti-mouse AF555, and Lycopersicon Esculentum (Tomato) Lectin (LEL, TL), DyLight 649, see **Table S30** for antibody product information) were incubated at 4°C for 12-24 hours in antibody carrier solution (5% NDS and 0.25% Triton X-100 in 1x PBS) at a dilution factor of 1:200. Gels were washed in 0.05x PBST (e.g., 500uL Triton X-100 in 1xPBS, 25mL of 1x PBS, up to 500mL of DIW) 6 times for 15 minutes each at room temperature to expand the gels for imaging.

The expansion factor for each gel in the morphology set was measured before and after expansion (in 0.05x PBST, the buffer used for washing before imaging) using a ruler (pre-expansion) and landmarks in the gel from tiled overview images obtained at 4x magnification (post-expansion). Measurement lengths smaller than 0.1cm, the smallest demarcation on the ruler, were estimated. We used the average expansion factor across samples for a given brain region or experiment to convert from physical (pre-expansion) to biological (post-expansion) units for scale bars, area, and volume calculations (**Table S31**). For the low expansion factor morphology dataset, final expansion factors for PFC, striatum, and thalamus were 3.65, 3.43, and 3.34, respectively. Images were obtained on an inverted Nikon w1 confocal microscope with a 40x water magnification lens. Images for the lower expansion factor morphology dataset were obtained at a 0.5µm z-step with 200ms exposure and 100% laser power for each optical channel. For some image stacks, the Lectin channel (640nm) was inadvertently obtained with 50ms exposure (as noted in **Table S32**), but this channel was not used for quantification, and only one of such astrocytes is shown in **Fig. S7** (for this astrocyte, the maximum Lectin contrast was decreased by 50% so that its Lectin contrast is similar to the others in the figure). We imaged the full z-stack for each channel separately using the Ti

Z-drive. When possible, we imaged the astrocytes that were: 1) sufficiently brightly labeled with GFP, 2) fully contained within the gel (volume not cut off, though many astrocytes were partially cut off for the experiment in **Fig. 7**, as noted in **Table S32**), 3) not overlapping with other brightly labeled astrocytes (though many fields of view did contain part of a second astrocyte, often more dimly labeled, for the experiment in **Fig. 7**, as noted in **Table S32**).

The expansion factor for the twice-expanded ExR gels for rDEG visualization was estimated at ~18x in 0.05x PBST based on measurements using the same protocol in  our prior studies[37,136]. Images for ~18x expanded ExR gels were obtained at 0.5μm z-step with 1s exposure and 100% laser power for each optical channel on a Nikon SoRa confocal microscope. We imaged 2048x2048x51 voxel volumes, with each z-step incrementing only after all 3 channels were imaged. In several fields of view, there are portions of empty gels lacking tissue, likely resulting from tissue being improperly anchored to the gel (see **Table S32**). These regions of empty did not affect our quantification of rDEG intensity in astrocytes across brain regions, as they were masked out during the GFP segmentation process (see "**Analysis of high-expansion factor ExR rDEG target images**" section below). Furthermore, we only included fields of view containing mostly tissues for quantification.

**Analysis of low-expansion factor ExR morphology images**.

Parameters for the following analyses were determined in advance of results compilation and statistical testing. All ExR images were background-subtracted in Fiji using the rolling ball algorithm, radius of 50 pixels, as in our previous studies[37,136]. To quantify astrocyte morphology from ~3.5x expanded astrocytes, images were processed as follows after background subtraction: conversion to grayscale, gaussian filtering with sigma = 2 using MATLAB's "imgaussfilt3", binarization with an intensity threshold at 1 standard deviation above the mean across the whole stack, and size filtering with a minimum of $10^6$ voxels using MATLAB's "bwareaopen". Importantly, the resulting segmentations allowed our morphological profiling to be robust to variations in GFP intensity observed in the astrocytes we imaged (**Fig. 7C**, **Movies S1-36**). If no astrocytes were detected after thresholding and filtering, the intensity threshold was lowered to 33.33% of the original (i.e., 0.3333 standard deviations above the mean intensity of the whole stack) and size filtered at the same minimum size. If no astrocytes were detected after lowering the intensity threshold, no segmentation was created for the image and it was excluded from further analysis. If there were two connected components (MATLAB's "bwconncomp", connectivity of 26) over the minimum size filter, a new size filter was set at 10 voxels less than the volume of the largest connected component, so that only the largest astrocyte remained. Volume, surface area, and equivalent diameter were then calculated from the largest connected component using MATLAB's "regionprops3", and converted to cubic and square microns using a weighted average of the physical pixel size divided by the expansion factor (e.g., ((⅔)*(0.1625/expansion_factor) + (⅓)*(0.5/expansion_factor))$^3$ for volume). Aspect ratio was calculated as the length of the first principal axis divided by the length of the second principal axis of the ellipsoid with the equivalent normalized second central moments as the connected component, calculated using MATLAB's "regionprops3".

After segmentation, we excluded 7 astrocytes that  had a large-volume second astrocyte in the 3D binary segmentation, and/or had low-quality 3D segmentations due to sparse signal (for 2 of such astrocytes, no 3D binary segmentation was created). Notes on the quality of each imaged astrocyte region of interest are provided in **Table S32**. We assessed whether the results of our statistical testing (see below) held up when more stringent quality control quality control (QC) criteria were applied to the 3D astrocyte segmentations being analyzed. Specifically, we removed astrocytes with sparse segmentations (6), segmentations including portions of second astrocytes in the field of view (22), cracked blood vessels (27), or which were partially cut off in the z-dimension (63). This higher-quality subset of the data included 74 astrocytes, most of which came from the thalamus (38%), followed by striatum (36%), and lastly PFC (26%). We repeated our linear mixed effects modeling on this dataset, and report the results in **Fig. S19A** and **Table S25**).

Sholl analysis was run using Fiji's SNT[137] Sholl Analysis[81] plugin, installed and accessed via the Neuroanatomy plugin, as described: https://imagej.net/plugins/snt/#installation. The center of each soma was manually annotated using the point selection tool and saved as an overlay. Sholl analysis was run in batch mode for each image using a custom macro that calls the legacy version of SNT's Sholl Analysis, with the following parameters: start radius of

2μm, end radius of 60μm, step size 2μm, and no polynomial fitting (see "batch_sholl_frombin.ijm" in our GitHub repository). The number of intersections at each radius for each cell was averaged over all non-excluded astrocytes in each region to create the plots in **Fig. 7E(vi)**.

Analysis of fractal dimension was performed in MATLAB on 3D binary segmentations using the box counting method via "boxcount", available on the MATLAB file exchange at https://www.mathworks.com/matlabcentral/fileexchange/13063-boxcount (written by Frederick Moisy, accessed August 2024). Briefly, a fractal set (in this case, a binary image of an astrocyte) is one that exhibits self-similarity at progressively smaller scales. The box counting method can be used to characterize the extent to which a set is fractal by counting the number of boxes of size R in a grid needed to cover the edges (in our case, transition from black to white) of an image at progressively smaller grid sizes[80]. A fractal image will have exponentially more detail (edges) at smaller grid sizes, require exponentially more boxes of size R to cover the set, and therefore will have a negatively sloped line on the log-log plot of N, the number of boxes vs. R, the size of each box in the grid. For a straightforward explanation of box counting, see https://fractalfoundation.org/OFC/OFC-10-5.html. The local fractal dimension can therefore be calculated as the slope of the log(N) vs. log(R) plot (df = -diff(log(n))./diff(log(r)), as in the boxcount function "Examples" tab). We took the mean fractal dimension over all 12 local slopes on the line to arrive at a single measure of fractal dimension for each astrocyte.

Statistical significance of differences between astrocytes from different brain regions was determined using a linear mixed effects model via the scipy statsmodels package's "mixedlm" function (https://www.statsmodels.org/stable/mixed_linear.html), where the outcome variable was the measure of interest, the coefficient was region assignment, and "animal" was the random effect group variable (see the associated Jupyter notebook, "analyze_ExR_4x_astromorph_results.ipynb" on our GitHub repository). P-values on the coefficients from striatum and thalamus were corrected for multiple comparisons with the Benjamini-Hochberg method using the scipy stats package's "false_discovery_control" function (https://docs.scipy.org/doc/scipy/reference/generated/scipy.stats.false_discovery_control.html). For the analysis where more stringent QC criteria were applied, the model failed to converge for fractal dimension. Results are provided in **Table S25**.

**Analysis of high-expansion factor ExR rDEG target images.** All ExR images were background-subtracted in Fiji using the rolling ball algorithm, radius of 50 pixels, as in our previous studies[37,136]. To create a segmentation of astrocyte processes (and if applicable, soma), the GFP channel was processed as follows: gaussian filtering with sigma = 10 using MATLAB's "imgaussfilt3", binarization with an intensity threshold at 0.75 standard deviations above the mean across the whole stack, median filtering with size 5x5x3 voxels, and size filtering with a minimum of 200 voxels using MATLAB's "bwareaopen". We then calculated the mean intensity (**Fig. 7F**) of the target channel (either Glast or Gat3) in the GFP+ region as follows: divided the raw pixel value of the target image by 2e16 (images were 16-bit), multiplied the resulting target image by the binary segmentation of the GFP channel, and divided the sum of this image by the number of nonzero pixels. We did the same for the reference synaptic protein channel, Cav2.1, which can be used to normalize the intensity values for the target channel (**Fig. S19B(i)**). The enrichment ratio was calculated as the mean intensity (of either the target channel or Cav2.1) within the GFP+ segmentation to the mean intensity outside the GFP+ region (**Fig. S19B(ii)**). Fields of view with poor Cav2.1 staining or lots of empty gel were excluded from analysis (see **Table S32**). Results are provided in Table S26.

**Generative AI.** ChatGPT (OpenAI, GPT-3.5 or 4.0) and GitHub Copilot (in Visual Studio Code) were used to aid computational scripting and debugging. On these occasions, we prompted it to interpret error messages and some lines of code generated by others, generate bash commands and scripts, accelerate existing Python code, generate some Python functions and code chunks for niche tasks (e.g., generating Venn diagrams and raster plots), and/or used the Copilot autocompletion function. ChatGPT and CoPilot-generated code blocks are annotated as such in our Python notebooks. However, generative AI did not meaningfully contribute to the intellectual development of experimental design, data analysis, or data interpretation of this manuscript. No generative AI was used to write or edit the manuscript.

**Supplementary Note 1**

The reader will note that not all marmoset astrocyte rDEGs followed the predicted expression pattern by RNAscope HiPlex FISH, and that there is a high degree of variability between the two replicates (**Fig. S9**). For example, *FOXG1* expression was spuriously detected in adult marmoset thalamus (at lower levels than PFC and striatum) by our CellProfiler pipeline, perhaps due to low and variable signal-to-noise. Nevertheless, *FOXG1* expression appears negligible in thalamic astrocytes when compared to PFC and striatum in our higher-quality marmoset sample, highlighting the limitations of a thresholding-based quantification approach (**Fig. S8**). In another example, the fraction of astrocytes positive for *KCND2* differs by several fold in the PFC between the two adult marmoset replicates. Our inability to quantitatively validate rDEG patterns in some replicates is likely due to variable tissue quality (even between regions, with striatum usually exhibiting less autofluorescence than PFC or thalamus), sampling a different position on the medial/lateral axis (especially for neonate marmoset samples), decreased signal-to-noise in later rounds of the RNAscope HiPlex protocol, overlapping RNA signal from adjacent cells of other cell types, or a combination of these factors. We found the RNAscope HiPlex protocol to produce variable results in marmoset tissue, especially in later rounds, even after several attempts at optimization. For this reason, we probed for *SPARC* and *KCNH7*, two probes with lower signal-to-noise in the HiPlex protocol, using RNAscope Multiplex Fluorescent v2 protocol, and produced much higher quality data. We suspect that the signal-to-noise for other rDEGs would be similarly improved, and some of the aforementioned issues resolved, if we used the RNAscope v2 protocol for all rDEGs though we did not perform these experiments ourselves.

In mouse, our quantification suggested *Gfap* expression was higher at both time points than predicted by snRNAseq, as was *Sparc* expression in neonates (**Fig. S11D**), findings which were not readily apparent by visual inspection of the images (**Fig. S10**). For *Gfap*, this discrepancy may reflect under-detection in the snRNAseq dataset, as evidenced by higher *Gfap* expression in the mouse ABCA MERSCOPE v1 dataset (**Fig. S11B**). In this and other cases, discrepancies between snRNAseq expression and FISH signal may be due to distinct gene capture strategies, or the use of probe target sequences on exons not highly expressed in astrocytes. For *Sparc*, overall expression in astrocytes at the neonate timepoint in the snRNAseq dataset is low (even in the thalamus, mean logCPM in astrocytes = 1.5 and fraction of cells in group for astrocytes < 30%). Low overall expression reduces the signal-to-noise ratio of FISH and makes our quantification methods more vulnerable to false positives (especially because the analyzer was blinded to gene identity, and did not know *a priori* that *Sparc* expression was predicted to be low overall in neonate astrocytes). At P4 in mouse, we noticed that *Sparc* expression was particularly high in endothelial cells, and what look to be pial astrocytes or other pial-associated non-neuronal cells lining the edges of the PFC, as well as in ependymal cells bordering the striatum (**Fig. S10E**). We suspected that co-localization of *Slc1a3* (which is not exclusive to astrocytes) and *Sparc* in these brain border- and vascular-associated cells was driving a discrepancy between the snRNAseq data and our FISH image quantification. We verified this by repeating *Sparc* quantification using a mask that removed the thin border sections (pial surface, ventricle) of the tissue in the PFC and striatum and increased intensity threshold in all regions, and found expression trends that more closely align with our snRNAseq predictions. Accordingly, we have replaced our Sparc quantification data (Fig. S11E) with these, and note that masking was done with the analyzer unblinded to rDEG identity and as such was not consistent with our OSF preregistration (see **Methods**).

Despite these limitations, taken together, our multiplexed FISH results confirm the divergent gene expression patterns of telencephalic and diencephalic astrocytes, with notable changes occurring over postnatal development.
