## Supplementary material for "Astrocyte regional specialization is shaped by postnatal development": Schroeder_2025_SIMovie_Links

**Supplementary Movie Dropbox Links**

1. Supplementary Movie 1: <https://www.dropbox.com/scl/fi/27jv3aavk2zfsqdkrv6zq/SI_Movie1_mouse1_pfc_40x_astro002_GFP.avi?rlkey=zzqovobezwcms8n8ex9vkcfxl&dl=0>
2. Supplementary Movie 2: <https://www.dropbox.com/scl/fi/qdp5dqs9kqnds3aalqj7e/SI_Movie2_mouse2_pfc_40x_astro001_GFP.avi?rlkey=xbe6yq3fvvm1dfyzexu7g2eto&dl=0>
3. Supplementary Movie 3: <https://www.dropbox.com/scl/fi/fpqyrope8jvkpogpc7j3f/SI_Movie3_mouse7_pfc_40x_astro001_GFP.avi?rlkey=3egqrt8jl5np6fglcvclrdv9x&dl=0>
4. Supplementary Movie 4: <https://www.dropbox.com/scl/fi/iiw4iwpduhll74p8a1qp0/SI_Movie4_mouse8_pfc_40x_astro002_GFP.avi?rlkey=f5djr5xwih3xjbwoz15xmmu7x&dl=0>
5. Supplementary Movie 5: <https://www.dropbox.com/scl/fi/15s9xjtnjvg20zxktlwua/SI_Movie5_mouse3_pfc_40x_astro003_GFP.avi?rlkey=z2gy0jmplmjhk4k6i4c9llugv&dl=0>
6. Supplementary Movie 6: <https://www.dropbox.com/scl/fi/0et2op9dn3sj0ji0ph2oj/SI_Movie6_mouse6_pfc_40x_astro002_GFP.avi?rlkey=zu26608slda4hroc24gw8zhrk&dl=0>
7. Supplementary Movie 7: <https://www.dropbox.com/scl/fi/barfvkfgfqc6zst21eljp/SI_Movie7_mouse1_str_40x_astro005_GFP.avi?rlkey=7v2h5icsbwmo9ctnifmyuy2wg&dl=0>
8. Supplementary Movie 8: <https://www.dropbox.com/scl/fi/p5x1shajrd9eppmq83igw/SI_Movie8_mouse2_str_40x_astro005_GFP.avi?rlkey=9nc32yveh1ruv3e63cena7zra&dl=0>
9. Supplementary Movie 9: <https://www.dropbox.com/scl/fi/q7l57ol2zgqw7zu9s5rwz/SI_Movie9_mouse5_str_40x_astro001_GFP.avi?rlkey=ayyv53pid4ujhedqlzcchaam9&dl=0>
10. Supplementary Movie 10: <https://www.dropbox.com/scl/fi/7lwzjdzt08i0pml2lrsaf/SI_Movie10_mouse8_str_40x_astro001_GFP.avi?rlkey=815x02vesrw48cvxf15kmwdyr&dl=0>
11. Supplementary Movie 11: <https://www.dropbox.com/scl/fi/xah9u6l9ywvyv6y0ixapt/SI_Movie11_mouse7_str_40x_astro001-2_GFP.avi?rlkey=d4fiy7rb1hlb0rpxl652je1h6&dl=0>
12. Supplementary Movie 12: <https://www.dropbox.com/scl/fi/aog91ge0n14bmhe9cid2e/SI_Movie12_mouse6_str_40x_astro004_GFP.avi?rlkey=f7jt7w0s96g1981f8gz93k4o5&dl=0>
13. Supplementary Movie 13: <https://www.dropbox.com/scl/fi/sil9neqq0vq4152y6i6u4/SI_Movie13_mouse1_thal_40x_astro008_GFP.avi?rlkey=3m3mwy7se94w9ah6xyw1y908i&dl=0>
14. Supplementary Movie 14: <https://www.dropbox.com/scl/fi/aigdxl733louvhn85dkrz/SI_Movie14_mouse2_thal_40x_astro002_GFP.avi?rlkey=bdr5c0ylpzk0j595ma3i0597m&dl=0>
15. Supplementary Movie 15: <https://www.dropbox.com/scl/fi/6k1u00q06ufhe7otnorra/SI_Movie15_mouse5_thal_40x_astro002_GFP.avi?rlkey=qlopmvplvtesjj30mzz4bzvj4&dl=0>
16. Supplementary Movie 16: <https://www.dropbox.com/scl/fi/zxsctomerza2rxqkb48jt/SI_Movie16_mouse8_thal_40x_astro004_GFP.avi?rlkey=3fa7px41sue702f0i3xrxgg11&dl=0>
17. Supplementary Movie 17: <https://www.dropbox.com/scl/fi/ndvluav7ssq9hv2va2fda/SI_Movie17_mouse7_thal_40x_astro001_GFP.avi?rlkey=8g8u6mf4009lmg5p5ie6vpchz&dl=0>
18. Supplementary Movie 18: <https://www.dropbox.com/scl/fi/211f7dlf2glv266hhvonk/SI_Movie18_mouse4_thal_40x_astro002_GFP.avi?rlkey=nf5qat2pm0ojwnu0rdc1wtijr&dl=0>
19. Supplementary Movie 19 (Related to Movie S1): <https://www.dropbox.com/scl/fi/72yqmkdzgk1pmh5oh4ea4/SI_Movie19_mouse1_pfc_40x_astro002_GFP_segmentation.avi?rlkey=19ipip25ekbk3zd0i1nqxiqb3&dl=0>
20. Supplementary Movie 20 (Related to Movie S2): <https://www.dropbox.com/scl/fi/kmlnzfgjofw88gck471sm/SI_Movie20_mouse2_pfc_40x_astro001_GFP_segmentation.avi?rlkey=zwjei510zd1d3wf6bzc9xt2be&dl=0>
21. Supplementary Movie 21 (Related to Movie S3): <https://www.dropbox.com/scl/fi/q4pf5wg1ytwin9ppla9dx/SI_Movie21_mouse7_pfc_40x_astro001_GFP_segmentation.avi?rlkey=qvrinbxpp0lr8shh20cbszkqw&dl=0>
22. Supplementary Movie 22 (Related to Movie S4): <https://www.dropbox.com/scl/fi/9njziq3klo5j6261jq1hs/SI_Movie22_mouse8_pfc_40x_astro002_GFP_segmentation.avi?rlkey=2e32iu43s849veyxckeju0jo3&dl=0>
23. Supplementary Movie 23 (Related to Movie S5): <https://www.dropbox.com/scl/fi/vy0nkhoo2a9xjj4mme5jz/SI_Movie23_mouse3_pfc_40x_astro003_GFP_segmentation.avi?rlkey=8ow13fhrs3g7o3vt38262fv90&dl=0>
24. Supplementary Movie 24 (Related to Movie S6): <https://www.dropbox.com/scl/fi/escnqcgkwwe4z72a2hpeh/SI_Movie24_mouse6_pfc_40x_astro002_GFP_segmentation.avi?rlkey=blg3s0ekhnnp73kxpcekyp9nh&dl=0>
25. Supplementary Movie 25 (Related to Movie S7): <https://www.dropbox.com/scl/fi/mai7lagx9qdfle9ra30t2/SI_Movie25_mouse1_str_40x_astro005_GFP_segmentation.avi?rlkey=fktwgz0jh9xf61oysgs7109s3&dl=0>
26. Supplementary Movie 26 (Related to Movie S8): <https://www.dropbox.com/scl/fi/9jm4g82iiaezc1i2o0y07/SI_Movie26_mouse2_str_40x_astro005_GFP_segmentation.avi?rlkey=md65ltw0msllbj2e1f92hfssw&dl=0>
27. Supplementary Movie 27 (Related to Movie S9): <https://www.dropbox.com/scl/fi/9gzqb5oi4ad8v2ig1fazk/SI_Movie27_mouse5_str_40x_astro001_GFP_segmentation.avi?rlkey=vbv81ea7lsqo7e2ba3op3x78o&dl=0>
28. Supplementary Movie 28 (Related to Movie S10): <https://www.dropbox.com/scl/fi/k2q9y7u33elg2teuxt09y/SI_Movie28_mouse8_str_40x_astro001_GFP_segmentation.avi?rlkey=vzf4w4se1fypoyvxklx3e2h1f&dl=0>
29. Supplementary Movie 29 (Related to Movie S11): <https://www.dropbox.com/scl/fi/nhalzwb801tigoepfiomo/SI_Movie29_mouse7_str_40x_astro001-2_GFP_segmentation.avi?rlkey=26ocyd3eiamf2r7q6cegyte8r&dl=0>
30. Supplementary Movie 30 (Related to Movie S12): <https://www.dropbox.com/scl/fi/uisi0wmpbvhwhhgkwf2ng/SI_Movie30_mouse6_str_40x_astro004_GFP_segmentation.avi?rlkey=fl7xyhjpxhwdamsd7dc6uwfl0&dl=0>
31. Supplementary Movie 31 (Related to Movie S13): <https://www.dropbox.com/scl/fi/og63oymuqo8cnebre1huy/SI_Movie31_mouse1_thal_40x_astro008_GFP_segmentation.avi?rlkey=6w6wdrc6axw0uhdjnfhvafh48&dl=0>
32. Supplementary Movie 32 (Related to Movie S14): <https://www.dropbox.com/scl/fi/jxrbghtuafj26v3ct9rv0/SI_Movie32_mouse2_thal_40x_astro002_GFP_segmentation.avi?rlkey=kbyc8o3m0x0je0e1cz39uaeep&dl=0>
33. Supplementary Movie 33 (Related to Movie S15): <https://www.dropbox.com/scl/fi/cty4d775tr5e7gzsgm6ae/SI_Movie33_mouse5_thal_40x_astro002_GFP_segmentation.avi?rlkey=7w5qgn5y7688ywf5yhx19tlcr&dl=0>
34. Supplementary Movie 34 (Related to Movie S16): <https://www.dropbox.com/scl/fi/5507y7yc5d0z2i65smcku/SI_Movie34_mouse8_thal_40x_astro004_GFP_segmentation.avi?rlkey=cyhthqbmj01hrdgamly6uzvrm&dl=0>
35. Supplementary Movie 35 (Related to Movie S17): <https://www.dropbox.com/scl/fi/j3q8x5wc4lbfuel5i4xz9/SI_Movie35_mouse7_thal_40x_astro001_GFP_segmentation.avi?rlkey=o7t9bcaey600120b69eg6exak&dl=0>
36. Supplementary Movie 36 (Related to Movie S18): <https://www.dropbox.com/scl/fi/bu5rhfbujboudaq9djma0/SI_Movie36_mouse4_thal_40x_astro002_GFP_segmentation.avi?rlkey=8ro1rt8pobz0i9num6pkgrntg&dl=0>
